# The genome of North American beaver provides insights into the mechanisms of its longevity and cancer resistance

**DOI:** 10.1101/2020.06.25.171322

**Authors:** Quanwei Zhang, Gregory Tombline, Julia Ablaeva, Lei Zhang, Xuming Zhou, Zachary Smith, Alus M. Xiaoli, Zhen Wang, Jhih-Rong Lin, M. Reza Jabalameli, Joydeep Mitra, Nha Nguyen, Jan Vijg, Andrei Seluanov, Vadim N. Gladyshev, Vera Gorbunova, Zhengdong D. Zhang

**Affiliations:** Department of Genetics, Albert Einstein College of Medicine, Bronx, NY, USA; Department of Biology, University of Rochester, Rochester, NY 14627, USA; Division of Genetics, Department of Medicine, Brigham and Women’s Hospital, Harvard Medical School, Boston, MA, USA; Departments of Medicine and Developmental and Molecular Biology, Albert Einstein College of Medicine, Bronx, NY, USA

**Keywords:** Beaver genome, gene duplication, positive selection, longevity

## Abstract

The North American beaver (Castor canadensis) is an exceptionally long-lived and cancer-resistant rodent species, and thus an excellent model organism for comparative genomic studies of longevity. Here, we utilize a significantly improved beaver genome assembly to assess evolutionary changes in gene coding sequences, copy number, and expression. We found that the beaver *Aldh1a1*, a stem cell marker gene encoding an enzyme required for detoxification of ethanol and aldehydes, is expanded (~10 copies vs. two in mouse and one in human). We also show that the beaver cells are more resistant to ethanol, and beaver liver extracts show higher ability to metabolize aldehydes than the mouse samples. Furthermore, *Hpgd*, a tumor suppressor gene, is uniquely duplicated in the beaver among rodents. Our evolutionary analysis identified beaver genes under positive selection which are associated with tumor suppression and longevity. Genes involved in lipid metabolism show positive selection signals, changes in copy number and altered gene expression in beavers. Several genes involved in DNA repair showed a higher expression in beavers which is consistent with the trend observed in other long-lived mammals. In summary, we identified several genes that likely contribute to beaver longevity and cancer resistance, including increased ability to detoxify aldehydes, enhanced tumor suppression and DNA repair, and altered lipid metabolism.

## Introduction

Rodent species show considerable variation in their maximum lifespans. Recent studies of several long-lived rodents have provided new insights into the mechanisms of longevity. For example, studies of the naked mole rats found unique amino acid changes in proteins involved in DNA repair and cell cycle (Kim, Fang et al. 2011), insulin β-chain associated with insulin misfolding and diabetes (Fang, Seim et al. 2014), decreased expression of genes in insulin/Igf1 signaling in liver (Kim, Fang et al. 2011), and increased expression of genes involved in DNA repair signaling (MacRae, Croken et al. 2015). In blind mole rats, Tp53 protein was found with an amino acid change associated with human tumors. This mutation leads to partial inactivation of Tp53 function and promotes p53 binding to promoters of cell cycle arrest genes rather than apoptotic genes (Ashur-Fabian, Avivi et al. 2004).

North American beaver (Castor Canadensis) has a maximum lifespan over 24 years and, with adults weighing from 24 to 71 lbs, is the second largest rodent species. It also shows resistance to cancer, despite a large body size and a long lifespan. Studies of several animals with large body sizes, long lifespans, and yet low rates of cancer revealed their enhanced antitumor mechanisms. For example, elephant, the biggest land mammal, was found have enhanced Tp53 activity by Tp53 expansion (~20 copies) (Abegglen, Caulin et al. 2015), while bowhead whale, the possibly longest-lived mammal, shows signals of natural selection in aging and cancer-associated genes (Keane, Semeiks et al. 2015). How beaver evolved to have cancer resistance and longevity is not clear.

The beaver genome assembly was first released in 2017 (Lok, Paton et al. 2017). Before analyzing the beaver genome, we re-sequenced and re-assembled the beaver genome to improve the assembly quality in terms of both the contig length and the assembly completeness (Zhou, Dou et al. 2020). To find potentially novel mechanisms of cancer resistance and longevity, using our improved beaver genome assembly, we explored three major types of signals of natural selection in beaver genome compared to other rodents: changes in gene copy numbers, positive selection that favored changes in amino acid residues, which further affect gene structure, interaction, and function, and changes in gene expression. Through a systematic, comparative analysis, we identified a striking multiplication of *Aldh1a1*, a stem cell marker gene, by around 10 copies in the beaver genome and beaver-specific expansion of several genes, including *Hpgd*, a tumor suppressor gene, and *Cyp19a1*, a gene involved in estrogen biosynthesis and linked to human longevity (Corbo, Ulizzi et al. 2011). We also found several genes associated with lipid metabolism, oxidation reduction, cancer suppression under positive selection and increased expression of DNA repair genes in beaver compared to mouse, consistent with observations in other long-lived species. In addition, we discovered that genes associated with lipid metabolism were likely under natural selection through changes in coding sequences, gene copy numbers, and gene expression levels.

## Results

### Genome annotation and phylogeny

Our beaver genome assembly (Zhou, Dou et al. 2020) consists of 739,342 scaffolds, with scaffold N50 = 24.31 Mb (the maximum was 93.06 Mb). This beaver genome assembly has a much higher completeness, with 95% complete BUSCOs (Benchmarking Universal Single-Copy Orthologs) (Simao, Waterhouse et al. 2015), compared to 83.1% of the beaver genome published previously (scaffold N50 = 318 kb and the maximum was 4.2 Mb) (Lok, Paton et al. 2017).

For gene annotation and downstream analysis, we selected scaffolds longer than 300 bp. There were 250,435 such scaffolds, and together they count for 96.6% of the assembled sequences. Using the Maker2 pipeline (Holt and Yandell 2011), we predicted 26,515 beaver genes with support from either transcriptomes or protein sequences (**Figure 1A**). Among these genes, 20,670 (78%) were found with functional domains by InterProScan (v5.25) (Jones, Binns et al. 2014). A genome is considered well annotated if more than 90% genes have an annotation edit distance (AED) score, which measures the goodness of fit of each gene to the evidence supporting it, lower than 0.5, and over 50% proteins contain a recognizable domain (Campbell, Holt et al. 2014). In our beaver genome annotation, ~90.7% gene models have AED scores lower than 0.5 (**Figure 1 – Supplement Figure S1**) and 78% gene products contain domains.

**Figure 1.**
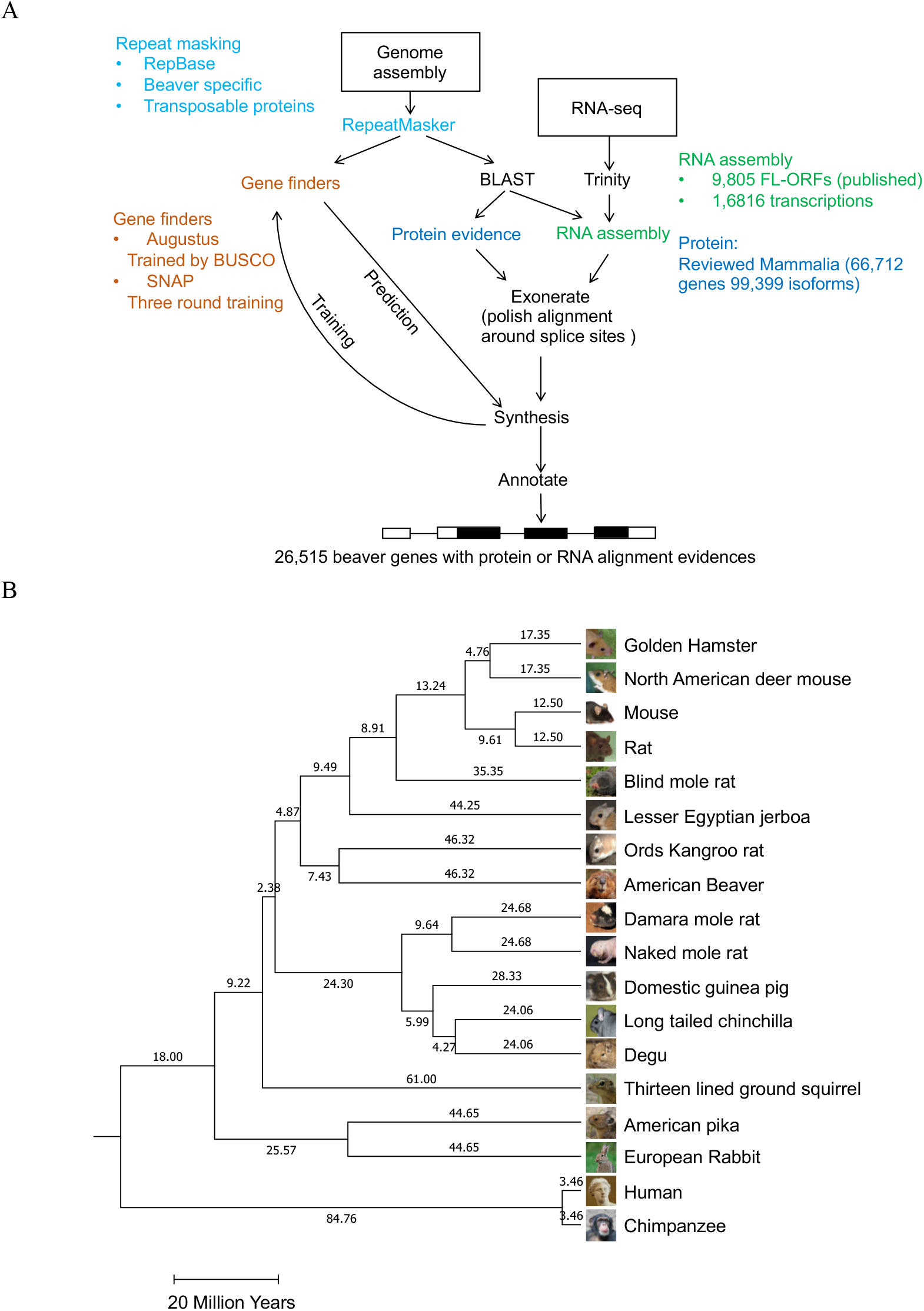
Genome annotation and phylogeny. **(A)** The beaver genome annotation pipeline. As a part of the annotation, we used 9,805 full-length open reading frames (FL-ORFs) from the published beaver genome (Lok, Paton et al. 2017). **(B)** Timed phylogeny. 5,087 single copy genes were used for the analysis.

A total of 14 rodent species, 2 rabbit species, and human and chimpanzee (as outgroup species), are included in our analysis. With 5,087 single copy genes across them, we generated their phylogeny and estimated that beaver and its evolutionally closest species in our set, Ord’s Kangroo rat, separated about 46 million years ago (**Figure 1B**).

### Significant expansion of beaver Aldh1a1 and its functional consequences

Using CAFE 3 (Han, Thomas et al. 2013), we identified significant (FDR<0.01) copy number increases of eight beaver genes (**Figure 2 – Supplement Figure S2**, see **Methods** for details). Most of these increases are not specific to beavers. One of them, the expansion of *Aldh1a1*, however, is striking in beaver, suggesting this expansion could be important for its cancer resistance and/or longevity. Therefore, we examined this gene in detail.

#### Expansion of beaver *Aldh1a1*

Compared with humans and chimpanzees, which have one copy of *Aldh1a1*, most rodent species have two copies of the gene (its close paralog in mouse is named as *Aldh1a7*). This indicates the duplication likely happened before the last common ancestor of rodents (**Figure 2 – Supplement Figure S3**). In contrast, 10 copies of *Aldh1a1* were predicted in the beaver genome, among which 2 copies are predicted as pseudogenes because of frameshift indels (**Figure 2 – Supplement Figure S4A, S4B**). One copy (ID: 266.2) has relatively lower annotation quality with two repeat fragments, which may indicate a prediction with a mixture from two copies of Aldh1a1.

We next examined every predicted copy of beaver *Aldh1a1* and carried out qPCR to experimentally quantify duplication events. We compared synteny of the *Aldh1a1* locus among beaver, human, and mouse (**Figure 2A**). Missing or unordered genes in a genomic region may indicate a poor assembly quality. A perfect match of the order of corresponding genes indicates good quality of genome assembly at this locus. After duplication, different gene copies evolved independently and accumulated differences in both sequence and gene structure. Using Apollo (Lee, Helt et al. 2013) and RNA-seq data, we manually improved the gene annotation of seven functional copies of beaver *Aldh1a1* (excluding two pseudogenes and the copy 266.2), which all have different gene structures (**Figure 2B**). Bona fide gene copies likely encode homologous proteins with different substitutions. We picked three sites in the coding sequences of exon 4 (**Figure 3C**), which can distinguish the seven copies from one another (**Figure 2C – Supplement Figure S4C**). The alignment of RNA-seq reads from beaver individuals indicates that those variable sites are genuine, and the different copies were transcribed simultaneously. While the RNA-seq read coverage at these variable sites was relatively low for the copy 266.5, we also analyzed another site, where the copy 266.5 is distinct from all other copies and with much higher RNA-seq reads coverage (**Figure 2 – Supplement Figure S4D**). To further validate the duplication of *Aldh1a1*, we performed qPCR analysis, which showed that there are around 10 copies of *Aldh1a1* in the beaver genome (**Figure 2D**).

**Figure 2.**
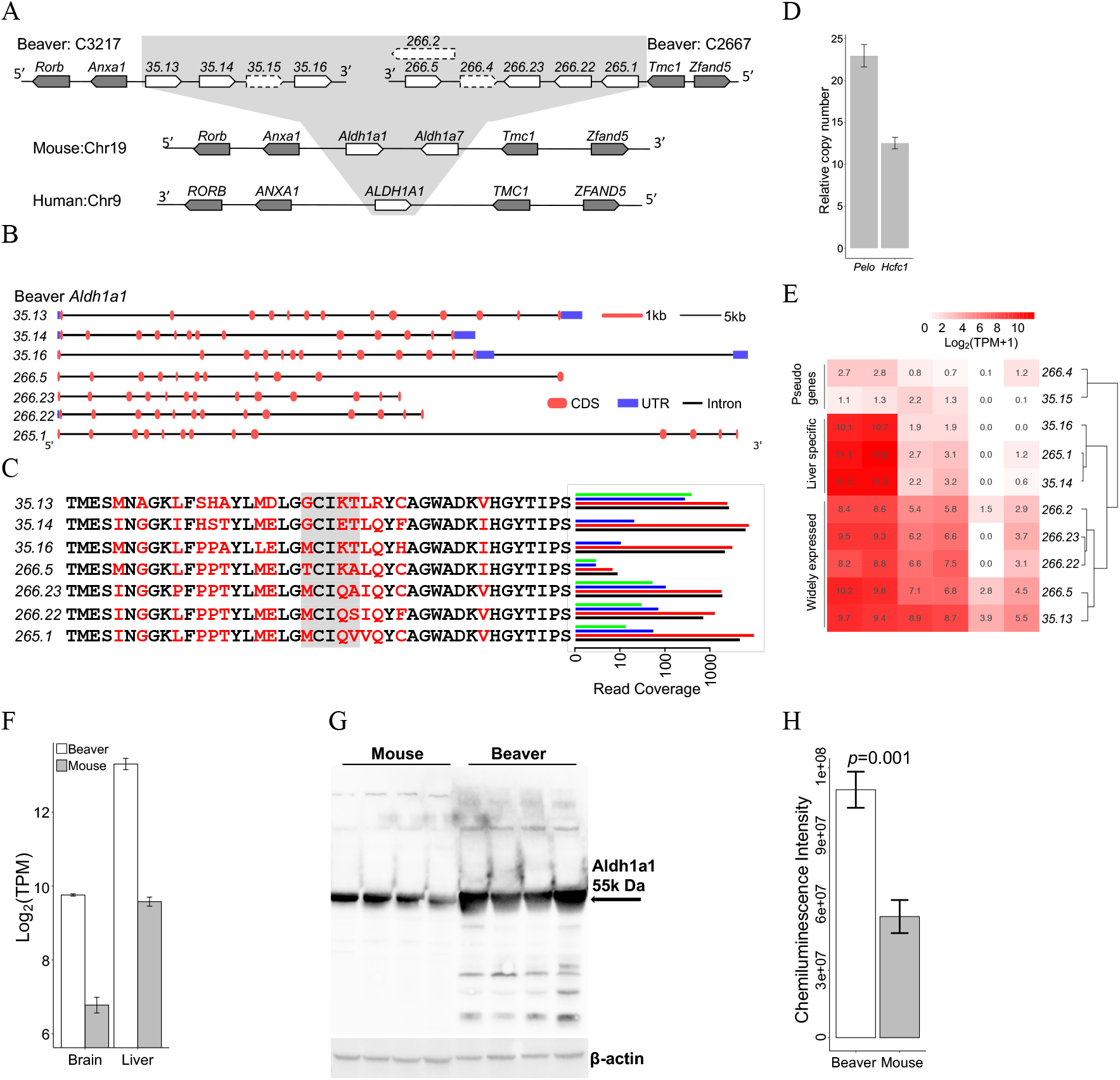
Expansion of *Aldh1a1* in the beaver genome. (**A**) The synteny of the *Aldh1a1* locus in the genomes of beaver, mouse, and human. We identified ten copies of *Aldh1a1* in the beaver genome, including two pseudogenes (35.15 and 266.4) and a low-quality copy (266.2). (**B**) Gene structure of the seven functional copy of *Aldh1a1*. (**C**) RNA-seq reads coverage at genomic locations of different amino acid residues among seven *Aldh1a1* gene products. Variable sites were highlighted in red, and the gray box indicates the selected sites where we checked the coverage of RNA-seq reads. We analyzed RNA-Seq reads from four beaver samples, B1-4 (**Supplementary Table 1**), separately. (**D**) Validation of the *Aldh1a1* copy number by qPCR. Two single-copy genes were used as references: *Hcfc1* on an autosome and *Pelo* on Chromosome X. (**E**) Expression profiles of *Aldh1a1* copies across several different beaver tissues. (**F**) *Aldh1a1* expression in liver and brain of both beavers and mice. For each tissue, we measured the expression of *Aldh1a1* in 2 beaver individuals and 8 mouse individuals.(**G**) Western blot of Aldh1a1 protein from beaver and mouse liver extracts of 4 different samples of each species. (**H**) Quantified Aldh1a1 levels in liver extracts. It is statistically higher (*P* = 0.001 by single side Welch Two Sample *t*-test) in beaver liver than in mouse liver.

**Figure 3.**
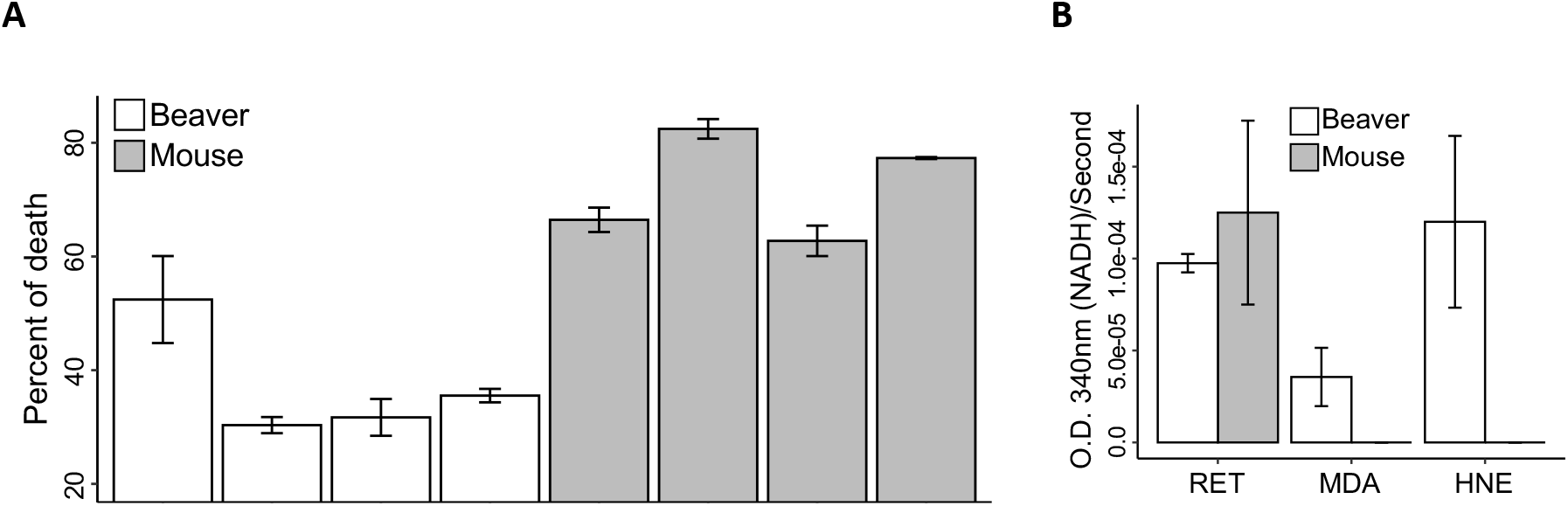
Functional characterization of *Aldh1a1*. (**A**) Cell viability. Beaver lung fibroblasts have lower percentage of death in the presence of high-concentration ethanol, normalized by corresponding controls (*p*=0.002 by mixed effects model, see **Methods**). (**B**) Aldehyde metabolic activity. Enhanced Aldh1a1 activity of beaver in aldehyde metabolism. Activities are normalized by corresponding controls without any addition (see **Methods**). RET: all-trans-retinal; MDA: malonaldehyde; HNE: 4-hydroxynonenal. Lung fibroblasts were used in experiments of panels A. Cytosolic extracts from hepatocytes were used for the experiment of panel B. 4 different biological samples of each species were used in experiments of **A)** and **B)**, with 3 replicates of each sample.

#### Duplicated *Aldh1a1* genes are transcribed and result in higher Aldh1a1 protein levels

All the predicted beaver *Aldh1a1* genes have regular gene structures with introns, and only two of them were identified as pseudogenes by GeneWise (Birney, Clamp et al. 2004) (see **Methods** and **Figure 2 – Supplement Figure S5**). At least seven copies of Aldh1a1 are of high annotation quality and transcribed in parallel in beaver individuals (**Figure 2C**). Based on their expression patterns in different tissues, beaver Aldh1a1 genes can be clustered into three groups (**Figure 2E**). As expected, the two pseudogenes show very low expression in all the tissues. While the highest expression of three copies in liver appears to be liver-specific, the expression of other copies is moderate in both liver and brain. To understand the overall transcriptional activity of *Aldh1a1* in beaver, we compared expression levels of *Aldh1a1* genes in liver and brain between beaver and mouse using RNA-seq data (see **Methods** for details). The overall expression of *Aldh1a1* was significantly higher in beaver liver (FDR = 2.23E-28, fold change = 10.04) and brain (FDR = 2.24E-12, fold change = 9.54) than that in the same mouse tissue, respectively (**Figure 2F**). We also examined protein level of Aldh1a1 by Western blotting. Using cytosolic extracts from liver cells of beaver and mouse, we found that there is more Aldh1a1 in beaver liver tissue than mouse liver tissue (**Figure 2G-H**). Together these results indicate that duplicated *Aldh1a1* copies are transcribed and result in higher protein levels in beaver tissue.

#### Potential positive selection of Aldh1a1 duplication

Gene duplications are often under relaxed purifying selection, as increased gene dosage tends to provide burden on the cell. The exceptional level of duplication of Aldh1a1 in beaver, however, suggests the presence of a positive (i.e., adaptive) selection. Aldh1a1 belongs to the Aldh super family: there are 19 and 21 Aldh genes in the human and mouse genomes, respectively. To investigate whether the duplication of Aldh1a1 in beaver is a compensation for loss of other Aldh genes or a result of natural selection, we manually checked all Aldh genes in the beaver genome (**Figure 2 – Supplement Figure S6**). Except Aldh1a1, beavers have the same set of Aldh genes as mice and humans with regular gene structures and open reading frames (**Figure 2 – Supplement Figure S7**). This conservation of other Aldh genes indicates that the significant expansion of beaver Aldh1a1 is likely a result of positive selection during evolution.

#### Beaver cells show enhanced tolerance of alcohol and aldehydes

Aldh gene products can protect organisms against damage from oxidative stress by processing toxic aldehydes generated as a result of lipid peroxidation (Singh, Brocker et al. 2013). Both alcohol and endogenous aldehydes can lead to DNA damage and increase mutations in stem cells (Garaycoechea, Crossan et al. 2018). We next tested the resistance of beaver cells to ethanol and aldehydes. When treated with 18% ethanol for 7 hours both beaver and mouse lung fibroblasts showed significantly decreased cell viability (**Figure 3A**). However, the reduction of cell viability was significantly lower for beaver lung fibroblasts than mouse cells (P = 0.002). This result suggests that higher Aldh1a1 levels in beaver cells lead to higher alcohol resistance. We also tested Aldh1a1 activity for three types of endogenous aldehydes: all-trans-retinal (RET), malonaldehyde (MDA) and 4-hydroxynonenal (HNE). Beaver and mouse liver extracts did not differ in their ability to process RET. However, beaver extract showed much stronger activity on MDA and HNE (***Figure 3B***). This result clearly shows that beaver liver possesses higher Aldh1a1 activity than mouse liver.

### Beaver-specific expansion of conserved genes

We identified 18 gene candidates for potential beaver-specific expansion and successfully validated five of them by qPCR (see **Methods** for details): *Hpgd* (*15-hydroxyprostaglandin dehydrogenase*), *Fitm1* (*Fat storage inducing transmembrane protein 1*), *Cyp19a1* (*Cytochrome P450 family 19 subfamily A member 1*), *Pla2g4c* (*Phospholipase A2 Group IVC*), and *Cenpt* (*Centromere protein T*) (**Figure 4A – Supplement Figure S8-S11**). There are two copies of *Hpgd*, a well-known tumor suppressor gene, in the beaver genome. The *Hpgd* loci in beaver, mouse, and human genomes show good synteny (**Figure 4B**), indicating a good assembly quality at this region. The beaver-specific duplication of *Hpgd* was validated by real-time qPCR (**Figure 4A**). We aligned beaver and mouse *Hpgd* sequences together (**Figure 4C**). While the two copies of beaver *Hpgd* have a high degree of sequence similarity, there are several amino acid residue differences. RNA-seq reads from individual beavers covered variable sites in both *Hpgd* copies, indicating that both copies are transcribed (and likely functional) and sequence differences were not due to DNA sequencing errors but instead divergent evolution after the duplication (**Figure 4D**). *Hpgd* is a tumor suppressor of many cancer types, including cancer of liver (Lu, Han et al. 2014), colon (Myung, Rerko et al. 2006), lung (Ding, Tong et al. 2005), and breast (Wu, Liu et al. 2017). Our RNA-seq data analysis showed although *Hpgd* was not differentially expressed in brain between beavers and mice, it was expressed significantly higher in beaver liver (fold change = 4.39, FDR = 4.56E-40), where *Hpgd* is more transcriptionally active (**Figure 4E**). The duplication of *Hpgd* and its higher expression likely contribute to the cancer resistance of beavers.

**Figure 4.**
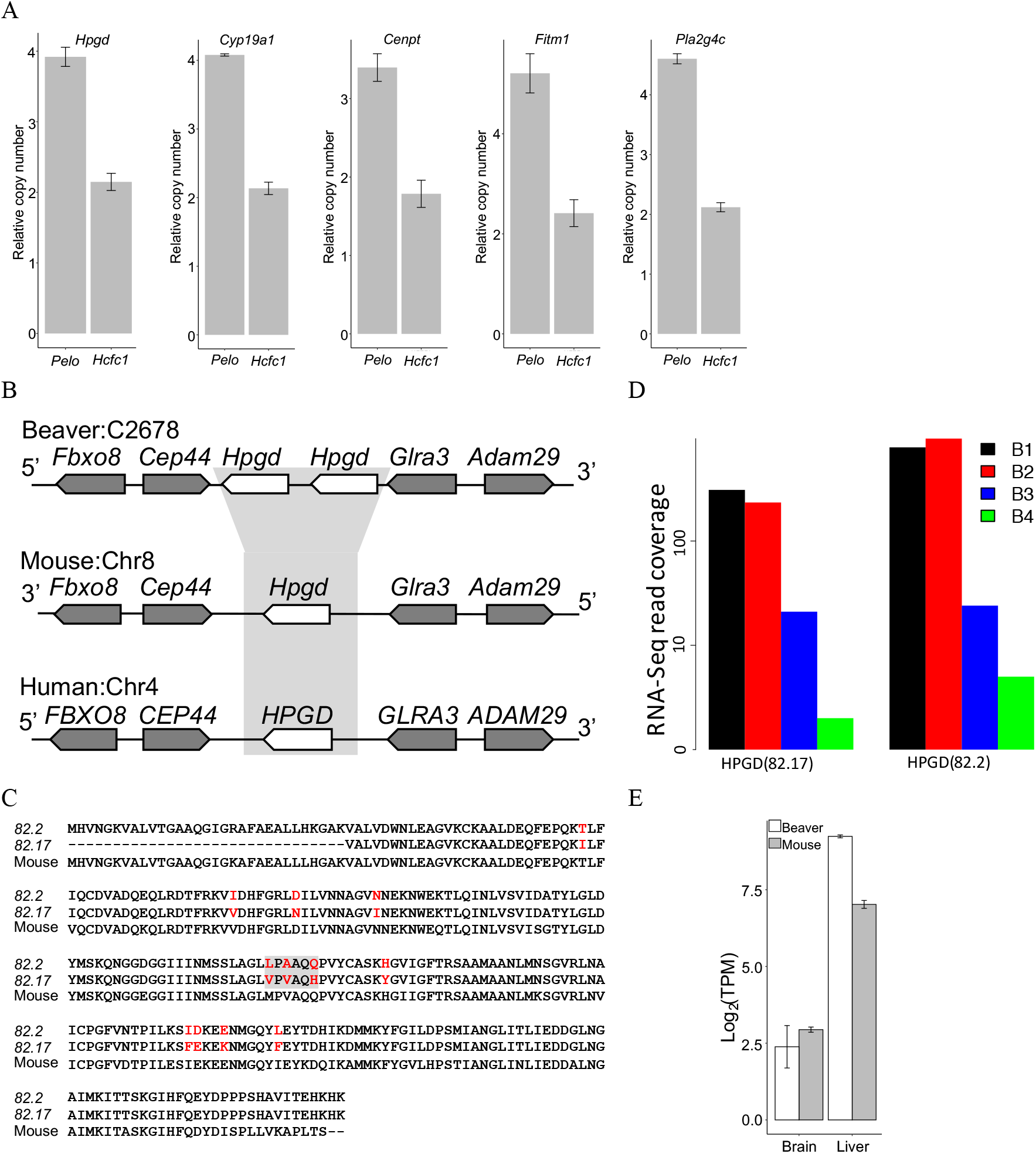
Beaver-specific gene expansion, including tumor suppressor *Hpgd*. (**A**) qPCR validation of the copy number of beaver-specific expanded genes. The copy numbers relative to the reference gene on chromosome X (i.e., *Pelo*) and on an autosome (i.e., *Hcfc1*) are shown for each candidate gene. (**B**) Synteny at the *Hpgd* locus. (**C**) Alignment of beaver and mouse Hpgd protein sequences. Variable sites are highlighted in red, and the gray box indicates the selected sites where we checked the coverage of RNA-seq reads. (**D**) Coverage of RNA-seq reads at the selected sites in individual beavers. (**E**) Expression of *Hpgd* in brain and liver of both beavers and mice.

### Beaver genes under positive selection are associated with tumor suppression and longevity

We identified 21 beaver genes putatively under positive selection (FDR < 0.01, **Table 1, Supplementary file 1**), using ‘branch site’ model (Zhang, Nielsen et al. 2005) conducted in PosiGene (Sahm, Bens et al. 2017) pipeline followed by manual curation (see **Methods** for details). Although not enriched at the pathway level, several top genes under positive selection are known to be associated with lipid metabolism (e.g., *Erlin2, Fabp3* and *Cilp2*), tumor suppression (e.g., *Vwa5a* and *Fabp3*), and oxidation reduction process (e.g., *Hsd17b1*, *Fabp3*, *Cox15*, *Cyb5a*, and *Aoc1*). Especially, several genes may be associated with aging/longevity. Different alleles of *Hsd17b1* were found significantly associated with human longevity in females (Scarabino, Scacchi et al. 2015). A single knockout of *Mtbp* in mice led to an extension of lifespan (Grieb, Boyd et al. 2016). Knockdown of *Mrpl37* increased lifespan of *C. elegans* by 41% on average (Houtkooper, Mouchiroud et al. 2013). Long-lived bats have greater abundance of Fabp3 in muscle mitochondria than that of short-lived mice, and thus its regulation of lipid may influence mitochondrial function (Pollard, Ingram et al. 2019). *FABP3* also acts as a tumor suppressor in human embryonic cancer cells and breast cancer (Song, Shen et al. 2012). In addition, Ptx3 is an inflammatory protein, which protects the organism against pathogens and controls autoimmunity, and is involved in tissue remodeling and cancer development (Doni, Stravalaci et al. 2019).

**Table 1.** Putative positive selection genes in beaver (FDR< 0.01)

| Gene | Gene name | Num. seq.<sup>1</sup> | Num. sites<sup>2</sup> | FDR |
| --- | --- | --- | --- | --- |
| *Mtbp* | MDM2 Binding Protein | 12 | 4 | 6.58E-04 |
| *Ptx3* | Pentraxin 3 | 16 | 6 | 1.56E-03 |
| *Tbxa2r* | Thromboxane A2 Receptor | 15 | 6 | 2.75E-03 |
| *Hsd17b1* | Hydroxysteroid 17-Beta Dehydrogenase 1 | 14 | 6 | 3.32E-03 |
| *Erlin2* | ER Lipid Raft Associated 2 | 18 | 3 | 3.86E-03 |
| *Fabp3* | Fatty Acid Binding Protein 3 | 17 | 7 | 3.86E-03 |
| *Adam19* | ADAM Metallopeptidase Domain 19 | 18 | 6 | 4.38E-03 |
| *Cilp2* | Cartilage Intermediate Layer Protein 2 | 14 | 30 | 5.07E-03 |
| *Chst12* | Carbohydrate Sulfotransferase 12 | 18 | 5 | 5.07E-03 |
| *Urb2* | URB2 Ribosome Biogenesis Homolog | 16 | 4 | 5.07E-03 |
| *Mrpl37* | Mitochondrial Ribosomal Protein L37 | 18 | 6 | 5.07E-03 |
| *Depdc7* | DEP Domain Containing 7 | 17 | 7 | 5.61E-03 |
| *Neu2* | Neuraminidase 2 | 12 | 3 | 6.07E-03 |
| *Vwa5a* | Von Willebrand Factor A Domain Containing 5A | 10 | 16 | 6.07E-03 |
| *Cox15* | Cytochrome C Oxidase Assembly Homolog COX15 | 18 | 4 | 6.64E-03 |
| *Gdf2* | Growth Differentiation Factor 2 | 17 | 9 | 7.31E-03 |
| *Sit1* | Signaling Threshold Regulating Transmembrane Adaptor 1 | 10 | 4 | 7.31E-03 |
| *Aoc1* | Amine Oxidase Copper Containing 1 | 17 | 9 | 8.02E-03 |
| *Cyb5a* | Cytochrome B5 Type A | 18 | 1 | 8.52E-03 |
| *Scn4b* | odium Voltage-Gated Channel Beta Subunit 4 | 16 | 6 | 8.64E-03 |
| *Il23a* | Interleukin 23 Subunit Alpha | 16 | 4 | 1.00E-02 |

**Notes:**
1. Num. seq.: the number of available orthologous sequences from a subset of our selected 18 species to build the high-quality alignment for detecting selection signals.
2. Num. sites: the number of amino acid residues that show selection signal in beavers.

Mtbp (Mdm2 Binding Protein) interacts with oncoprotein mouse double minute 2 (Mdm2) and enhances its stability, which promotes degradation of Tp53 (**Figure 5 – Supplement Figure S12**). Sites with positive selection signals were identified in exons encoding the “mid-domain” and the “C-domain” of the Mtbp protein (**Figure 5A**). In addition to codon changes (i.e., selection signals identified by PosiGene (Sahm, Bens et al. 2017)), there are also indels in beaver *Mtbp*, compared with the ortholog from other rodent species. Especially, there is a 15-bp insertion in exon 18 of beaver *Mtbp*, which resembles an insertion at a similar location in Mtbp of the long-lived naked mole rat. We explore potential functional effects of changes in codons and indels using PROVEAN (Choi and Chan 2015), PolyPehn2 (Adzhubei, Jordan et al. 2013) and CADD (Rentzsch, Witten et al. 2019) by considering changes from those in human genome (human sequences are the consensus sequence at those loci) to those in beaver genome (**Figure 5B**). Two sites under positive selection are consistently predicted to be deleterious, suggesting they could result in decreased Mtbp function and hence decrease the stability of Mdm2, leading to higher Tp53 activity. We also checked the top three sites under positive selection identified in the multiple sequence alignment of *Mtbp* from 62 mammals (**Figure 5 – Supplement Figure S12B-D**). We checked a few other sequence changes at the positive selection sites in other mammals and found only changes in beaver *Mtbp* were predicted as deleterious (**Figure 5 – Supplement Figure S12D**), which may contribute to its cancer resistance. Haploinsufficiency of *Mtbp* in mice delays spontaneous cancer development and extend lifespan by enhancing Tp53 function (Grieb, Boyd et al. 2016).

**Figure 5.**
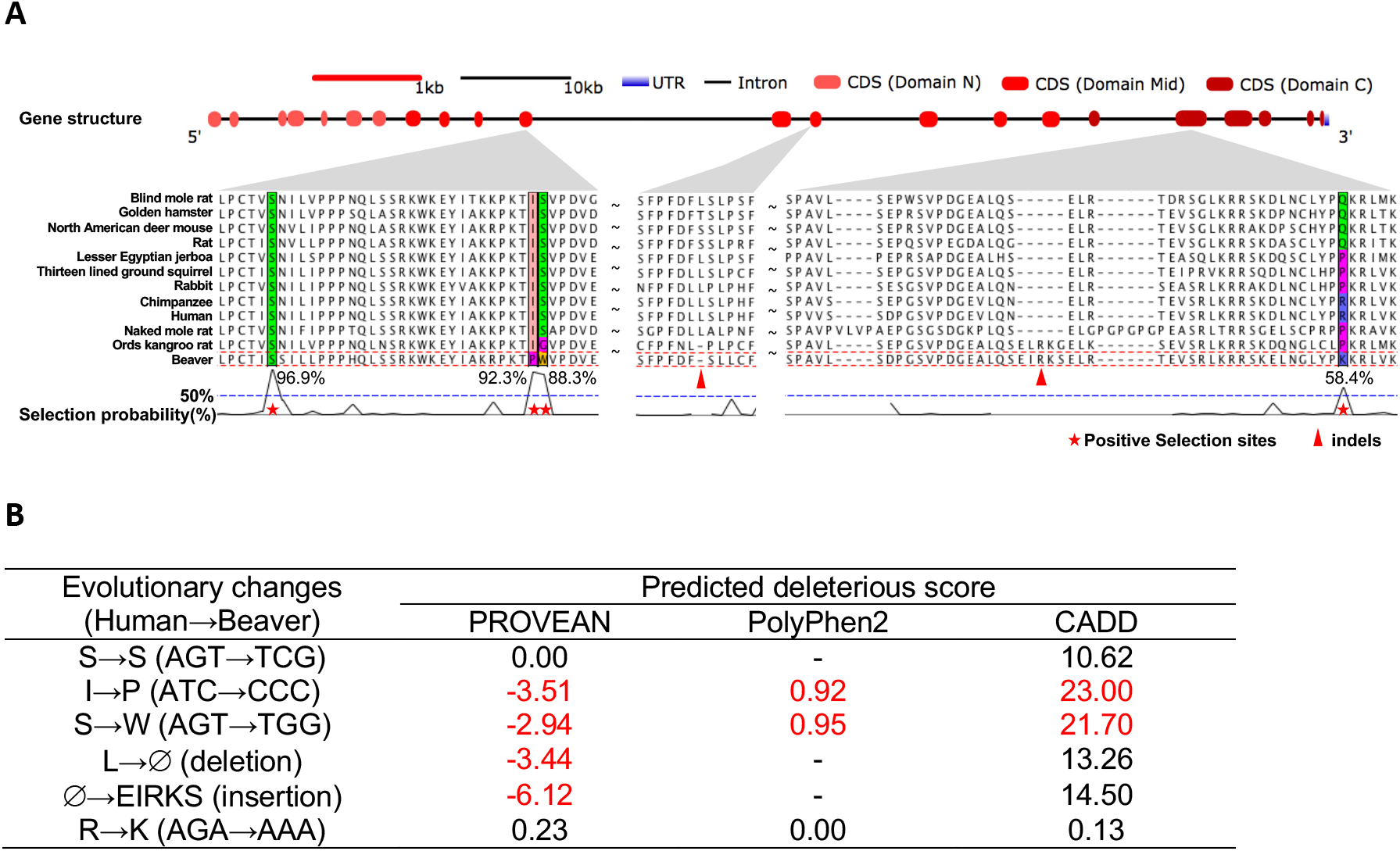
*Mtbp* is under positive selection. (**A**) Positive selection signals in *Mtbp* gene. We observed nucleotide positions and indels under positive selection in coding sequences of three exons, corresponding to the middle and the C domains of Mtbp. (**B**) Nucleotide sequence changes between the beaver and the human genomes. We predicted the deleteriousness of these changes based on their human genome annotation, using the following metrics and thresholds: PROVEAN score < −2.5, 0.85 < Polyphen2 score < 1.0, and CADD score > 20.

### Beaver displays enhanced expression of DNA repair genes and changes in genes involved in lipid metabolism

Among 12,090 genes with one-to-one orthology between beaver and mouse, we detected 2,892 (1,652 up-regulated and 1,240 down-regulated) and 2,765 (1,534 up-regulated and 1,231 down-regulated) differentially expressed genes (fold change > 2.5 and FDR < 0.001) in liver and brain, respectively, between beavers and mice (see **Methods** for details). Among REACTOM pathways, both up-regulated and down-regulated differentially expressed genes were enriched in lipid metabolism pathway in both liver and brain (**Figure 6**). Such result indicates significantly divergent evolution of lipid metabolism between beaver and mouse. Up-regulated genes in both livers and brains of beavers were also enriched in hemostasis, DNA repair, and cell cycle pathways. Especially, “nucleotide excision repair” and “base excision repair” pathways are enriched with up-regulated genes in beaver brains. Up-regulated genes in beaver livers are enriched in the Tp53 signaling pathway among 50 hallmark gene sets from the Molecular Signatures Database (Subramanian, Tamayo et al. 2005), which represent well-defined biological states or processes with coherent expression. Linking to human longevity and age-related diseases (Wolfson, Budovsky et al. 2009), overexpression of genes in “focal adhesion” (**Figure 6A, 6B**) may also contribute to beaver’s longevity.

**Figure 6.**
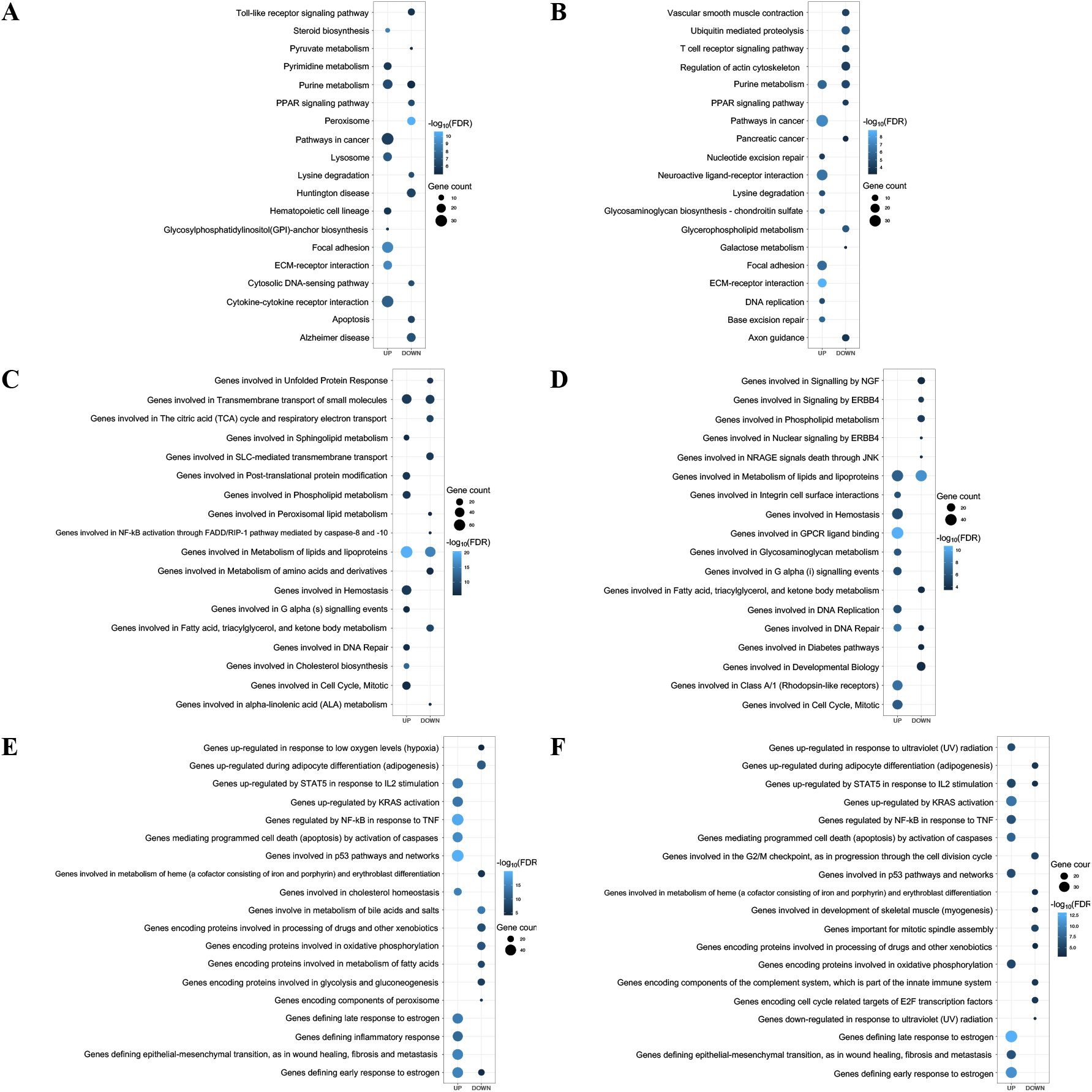
Gene sets enriched with genes differentially expressed between beavers and mice. (**A**) KEGG signaling pathways enriched with differential expression genes in liver tissue. (**B**) KEGG signaling pathways enriched with differential expression genes in brain tissue. (**C**) REACTOM pathways enriched with differential expression genes in liver tissue. (**D**) REACTOM pathways enriched with differential expression genes in brain tissue. (**E**) Hallmark gene sets enriched with differential expression genes in liver tissue. (**F**) Hallmark gene sets enriched with differential expression genes in brain tissue. UP: enriched pathways of up-regulated genes in beaver tissue comparing to mouse tissue. DOWN: enriched pathways of down-regulated genes in beaver tissue comparing to mouse tissue. Top 10 enriched pathways of up- and down-regulated genes are shown for each type of gene sets. Hallmark gene sets from the Molecular Signatures Database represent specific well-defined biological states or processes and display coherent expression (Liberzon, Birger et al. 2015).

Among differentially expressed genes, *Igf2* (insulin-like growth factor 2) showed much higher expression in beaver liver than in mouse liver (FDR = 3.06E-135, top 20th up-regulated gene in beaver liver), and its binding protein Igf2bp2 also shows significant up-regulation in beaver liver (FDR = 7.69E-07) **(Figure 6 – Supplement Figure S13**). Expression of *Igf2* significantly decreases after birth in liver for most mammals, including mouse and rat. Interestingly, *Igf2* and *Igf2bp2* show relatively higher expression levels in livers of naked mole rats and Damaraland mole rats (Fang, Seim et al. 2014, Ma and Gladyshev 2017), both long-lived rodents.

## Discussion

*Aldh* gene products process aldehydes generated as a result of lipid peroxidation, thus protecting the organism from the consequence of oxidative stress (Singh, Brocker et al. 2013). A recent study showed that both alcohol and endogenous aldehydes damage chromosomes and increase the mutation rate of stem cells (Garaycoechea, Crossan et al. 2018). We found that beaver cells exhibit better tolerance to ethanol than mouse cells and show strikingly enhanced capabilities for metabolizing malonaldehyde (MDA) and 4-hydroxynonenal (HNE) (**Figure 3**). While MDA is the most mutagenic aldehyde product of lipid peroxidation, HNE is the most toxic (Ayala, Munoz et al. 2014). The *Aldh* upregulation occurs in mammals in response to lipid peroxidation (Vassalli 2019), which is a major source of endogenous aldehydes. Polyunsaturated fatty acids (PUFAs) are more susceptible to oxidation than monounsaturated fatty acids, and beavers have a high proportion of PUFAs, which is unusual among mammals (Martysiak-Zurowska, Zalewski et al. 2009, Zalewski, Martysiak-Zurowska et al. 2009, Domaradzki, Florek et al. 2019). For example, beaver’s tail fat contains over 80% unsaturated fatty acids (Zalewski, Martysiak-Zurowska et al. 2009). Studies of diving mammals indicate potential function of PUFAs in oxygen conservation, through reducing heart rate, to enhance their diving ability (Trumble and Kanatous 2012). Consistently, the heart rate of beavers is 100 beats/minute during rest and 50 beats/minute while diving (Muller-Schwarze 2003), which is much lower than that (~250 beats/mins) of other rodents (Carpenter and Marion 2013). However, high level of PUFAs will result in increased lipid peroxidation, which is reversely correlated with lifespan of diverse species, including mammals (Hulbert, Kelly et al. 2014). Longevity of beavers indicates the presence of a potential protection mechanism against lipid peroxidation. Compared to *Aldh2* and *Aldh3a1*, *Aldh1a1* is the most important enzyme to oxidize aldehydes formed by lipid peroxidation in murine hepatocytes (Makia, Bojang et al. 2011). The higher susceptibility to lipid peroxidation and the associated oxidative stress in beavers may exert selective pressure for increased expression of *Aldh1a1*. Gene duplication is one way to increase gene dosage rapidly. From this point of view, *Aldh1a1* duplication in beaver is very likely a result of natural selection, which significantly improves its tolerance against oxidative stress, and so contributes to its longevity. In addition to gene duplication, our data also showed different expression patterns of the copies across tissues (**Figure 2E**), which warrants future studies to explore whether diverse selections have shaped regulation of *Aldh1a1* copies. While no difference in Aldh1a1 activity on RET was observed between beavers and mice, Aldh1a1 in beavers showed much higher activities on MDA and HNE than that in mice.

Increased Aldh1 activity is expected to increase organism’s resistance to oxidative stress. Enhanced tolerance for the oxidative stress has been found in long-lived fruit flies comparing those with normal lifespan (Deepashree, Niveditha et al. 2019). In human, centenarians have been found with less oxidative stress damages comparing to controls (Belenguer-Varea, Tarazona-Santabalbina et al. 2019). Linked to multiple age-related diseases, the oxidative stress is likely one of major contributors to aging (Liguori, Russo et al. 2018). Lipoxidation increases with age (Mitchell, Buffenstein et al. 2007), while reactive aldehyde, a known carcinogen, interferes with DNA replication, causes DNA damage, and induces formation of DNA adducts (Langevin, Crossan et al. 2011). *Aldh1a1* duplication can increase cellular protection against these toxins. Our finding of better toleration of ethanol by beaver cells is consistent with a previous human study, which showed low *Aldh1a1* activity might account for alcohol sensitivity in some Caucasian populations (Marchitti, Brocker et al. 2008). Stem cell exhaustion is one of hallmarks of the aging process. *Aldh1a1* is also a marker gene associated with stemness of cells and expressed higher in stem cells (Li, Condello et al. 2017). *Aldh1a1* duplication may also contribute to stem cell maintenance in beavers. This hypothesis is consistent with results from other studies, which found unsaturated fatty acids can maintain cancer cell stemness (Mukherjee, Kenny et al. 2017) and positive association between *Aldh1a1* expression and lipid unsaturation level (Li, Condello et al. 2017). Thus, *Aldh1a1* duplication may affect beavers’ aging process by increasing both resilience to oxidative stress and the stemness of beaver cells. Mice deficient in both *Aldh1a1* and *Aldh3a1* had fewer hematopoietic stem cells, more reactive oxygen species, and increased sensitivity to DNA damage (Gasparetto, Sekulovic et al. 2012), all of which are early aging phenotypes.

Several studies have shown the association of *Aldh1a1* expression with the aging process. The expression of *Aldh1a1* dramatically increases with age in mouse hematopoietic stem cells (Levi, Yilmaz et al. 2009). It is significantly lower in CD4^+^ T cells from aged mice than those from young mice, and its low expression may affect migration of T cells in aged mice (Park, Miyakawa et al. 2014). However, how and in which tissues the expression of *Aldh1a1* is significantly changed with age is not clear. To explore this, we used human gene expression data in diverse tissues from GTEx (Carithers, Ardlie et al. 2015) and studied *Aldh1a1* expression pattern during aging in the presence of several potential covariates (see **Methods**). In several tissues, the expression of *Aldh1a1* significantly changes with age (FDR < 0.01) (**Figure 7**). For example, it significantly increases with age in adipose tissue (*r* = 0.17, FDR = 0.0003), which may be associated with an increase in the percentage of body fat among the old (St-Onge and Gallagher 2010). It also significantly decreases with age in several human brain regions. This may be associated with an increase in some brain disorders among the old, as patients with Parkinson’s disease tend to have a decreased expression of *Aldh1a1* in corresponding brain regions (Galter, Buervenich et al. 2003). It is believed that ALDHs play a significant role in neuroprotection (Marchitti, Deitrich et al. 2007). For example, ALDH1A1 has been found as a marker of astrocytic differentiation during brain development (Adam, Schnell et al. 2012). Thus, *Aldh1a1* duplication may also protect beaver from age-related brain impairment.

**Figure 7.**
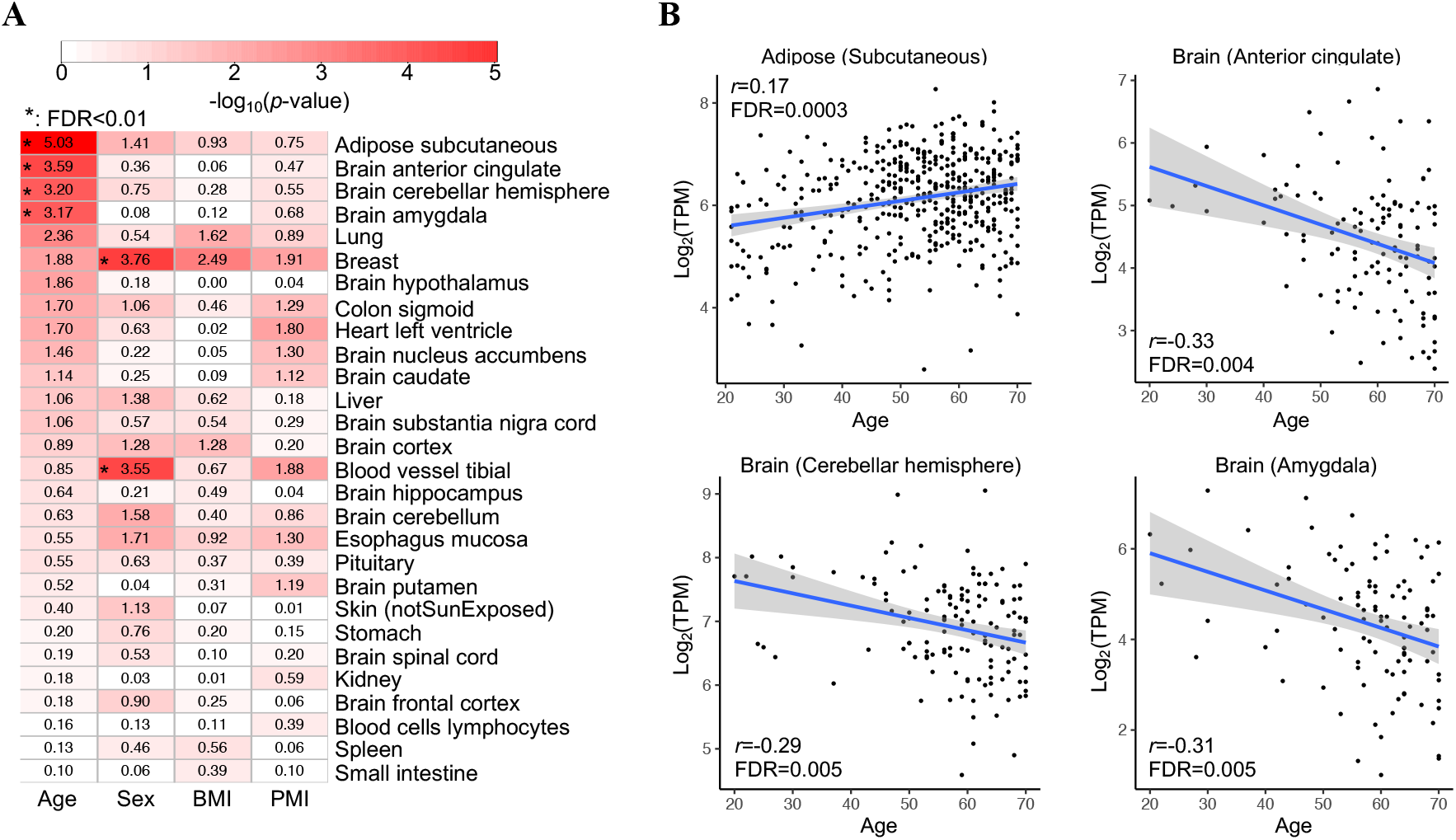
Age-associated expression of *ALDH1A1* in human. (**A**) Correlation of *ALDH1A1* expression with age, sex, body mass index (BMI), and post-mortem interval (PMI, minutes between death and sample collection). Tissues are ordered according to the *P*-values of the correlation with age. A significant correlation (FDR < 0.01) is denoted by an asterisk. (**B**) Significant correlation between *ALDH1A1* expression and age in four human tissues. The gray shaded area around the regression line indicates 95% confidence interval.

Compared to other mammals, the adipose tissue and muscles of beavers show different lipid composition, with the highest proportion of PUFAs (Martysiak-Zurowska, Zalewski et al. 2009, Zalewski, Martysiak-Zurowska et al. 2009, Domaradzki, Florek et al. 2019), which may indicate an unusual way of lipid metabolism and anti-lipoxidation in beavers. Consistent with the unusual lipid composition in beavers, we found genes differentially expressed between mice and beavers are most enriched in lipid metabolism (**Figure 6**). Besides, several genes expanded in the beaver genome, including *Aldh1a1*, *Hpgd*, *Fitm1*, *Cyp19a1*, and *Pla2g4c*, are also associated with lipid metabolism. *Aldh1a1* knockout mice show decreased accumulation of both subcutaneous and visceral fat pads (Ziouzenkova, Orasanu et al. 2007), indicating that *Aldh1a1* duplication may contribute to accumulation of protective layers of fat which beavers evolved to provide insulation in cold water. Furthermore, *Fabp3*, one of the top positively selected genes in beaver, has also been found to be associated with intramuscular fat deposition and body weight in chicken (Ye, Chen et al. 2010) and pigs (Gerbens, van Erp et al. 1999). In mice, Fabp3 is essential for fatty acid oxidation in brown adipose tissue and plays a central role in cold tolerance (Vergnes, Chin et al. 2011).

Among genes duplicated specifically in beavers, *Hpgd* is highly expressed in regulatory T cells, which prevents autoimmunity and maintains adipose tissue homeostasis (Schmidleithner, Thabet et al. 2019). *Fitm1* plays an important role in lipid droplet accumulation. *Cyp19a1* is involved in the synthesis of cholesterol, steroids, and other lipids. It also catalyzes estrogen biosynthesis, which enhances healthy aging and human longevity (Horstman, Dillon et al. 2012). A significant association between *Cyp19a1* polymorphisms and longevity was observed in humans (Corbo, Ulizzi et al. 2011). As a member of the phospholipase A2 enzyme family, *Pla2g4c* plays a role in hydrolyzing glycerol-phospholipids to produce lysophospholipids and free fatty acids. Although not enriched on the pathway level, among the top 10 putative positive selection genes, three are associated with lipid metabolism. *Erlin2* is a regulator of cytosolic lipid of cancer cells (Wang, Zhang et al. 2012)*. Fabp3* plays a role in metabolism and transport of long chain polyunsaturated fatty acid, and its role in lipid regulation may be important for mitochondrial function (Pollard, Ingram et al. 2019). Genetic variant association studies indicate *Cilp2* regulates lipid species in both mice and humans (Jha, McDevitt et al. 2018). All these genetic and phenotypic features indicate a natural selection on lipid metabolism in beavers. Genes involved in lipid composition have been found under stronger selective pressure in long-lived species (Jobson, Nabholz et al. 2010). The evolution of lipid metabolism is connected to mammalian longevity (Li and de Magalhaes 2013), and the important role of lipid metabolism in aging and lifespan regulation have been observed in different species, including humans (Johnson and Stolzing 2019). We hypothesize that altered regulation of genes associated with lipid metabolism in the beaver genome likely contributes to its longevity. Although how altered lipid metabolism affects beavers’ longevity needs further exploration, several unusual fatty acids, such as branched-chain (BCFA) and odd-chain (OCFA), which are typical for ruminants, were detected in beaver muscles. BCFA and OCFA have shown positive biological effects on anti-cancer activity (Domaradzki, Florek et al. 2019).

Like other long-lived rodent species, beavers are also cancer resistant. The malignant transformation of beaver cells needs both inactivation of Tp53, Rb1 and Pp2ca and constitutive activation of telomerase and HRas (Seluanov, Gladyshev et al. 2018). Such requirements for beaver cells are more stringent than for mouse cells and identical to those for human cells, which makes beavers a promising model to study human cancer. Several findings from this study may shed some light on the cancer resistance of beavers. *Hpgd*, a known tumor suppressor for many types of cancer, has been duplicated uniquely in beaver among rodents. The expansion of *Aldh1a1* can better protect beaver cells against lipid aldehyde, which can damage DNA and proteins, by metabolizing reactive aldehyde into harmless acetic acid. Increased expression of genes in DNA repair pathway can also protect beavers from cancer.

## Materials and Methods

### De novo genome assembly

We generated the beaver genome assembly using the third-generation sequencing technology and a new scaffolding method. A single ‘Chicago’ library (sequenced at a ~87X coverage) was generated by Dovetail Genomics. Those sequence data were mapped to our initial assembly (with scaffold N50 = 55.69kb), which was generated by using AllPath-LG with default parameters on whole genome shutgun sequences from the Illumina HiSeq platform. And then HiRise scaffolding pipeline (Putnam, O’Connell et al. 2016) was used to generate the dovetail assembly (with a scaffold N50 = 20.99 Mb). Finally, we used the published long PacBio reads to fill gaps in the dovetail assembly (Lok, Paton et al. 2017).

### BUSCO-based genome quality assessment

We used BUSCO v2(Simao, Waterhouse et al. 2015) to evaluate the completeness of the beaver genome assemblies. Briefly, BUSCO assesses the genome by searching for presence of near-universal single-copy genes from OthoDB v9(Zdobnov, Tegenfeldt et al. 2017). Absence of those conserved genes indicates incompleteness of the genome. In our analysis, we used the mammalian gene set consisting of 4,104 single-copy genes that are present in more than 90% mammalian species.

### Training for ab initio gene prediction

Augustus was trained by running BUSCO(Simao, Waterhouse et al. 2015) with the ‘--long’ parameter, which performs a full optimization of training for Augustus gene finding. SNAP was trained following the previously described pipeline(Campbell, Holt et al. 2014) with three iterations.

### Gene structure and function annotation

Maker2(Holt and Yandell 2011) was used for gene structure prediction (see **Figure 1A** for the pipeline). Repeating elements were first masked by RepeatMasker (v4.07)(Tarailo-Graovac and Chen 2009), with RepBase repeat libraries (20170127)(Bao, Kojima et al. 2015) and beaver-specific repeating elements constructed by RepeatModeler (version 1.0.10) following the instruction given by http://weatherby.genetics.utah.edu/MAKER/wiki/index.php/Repeat_Library_Construction--Basic. RepeatRunner was then used to further identified more divergent transposable protein elements provided by Maker2.

With the repeat-masked assembly, genes were predicted by ab initio gene predictors (i.e., SNAP(Korf 2004) and Augustus(Stanke and Waack 2003)) and evidence-based gene calling (i.e., using transcript assembly and protein sequences). For beaver gene transcripts, we used our 16,816 assembled transcripts (see above) and the 9,805 full length open reading frames from the published beaver genome(Lok, Paton et al. 2017). For protein evidence, about 66.7 thousands of reviewed mammalian protein sequences from Swiss-Prot(The UniProt 2017) were used for homolog-based gene prediction. These transcript and protein sequences were used to train the ab initio gene predictors (see above), polish the predicted gene models, and evaluate each predicted gene model. Finally, we predicted 26,515 beaver genes with evidence support from either transcript or protein sequences.

For each gene, Maker2 calculates an annotation edit distance (AED) score, which measures the goodness of fit of each gene to the evidence supporting it. A genome is usually considered well annotated if more than 90% genes have AED scores < 0.5 and over 50% proteins contain a recognizable domain(Campbell, Holt et al. 2014). In our beaver genome annotation, ~90.7% gene models have AED scores lower than 0.5 (**Figure 1 – Supplement Figure S1**) and 78% of predicted gene products contain known protein domains by InterProScan (v5.25)(Jones, Binns et al. 2014).

### Orthology and phylogeny analyses

To identify gene families across species, we used OrthoDB (release 9)(Zdobnov, Tegenfeldt et al. 2017), which covers more than 600 eukaryotic species with functional annotation from more than 100 sources. In our analysis we selected 14 rodent and rabbit species from OrthoDB and mapped beaver and naked mole protein sequences to OrthoDB. We chose human and chimpanzee as out-group species. We identified ~21,000 gene families, among which 5,087 gene families have single copy across all 18 species.

Single-copy gene families were used for phylogenetic analysis. Briefly, orthologs from each family were first aligned by MUSLE(Edgar 2004). Poorly aligned regions was removed by TrimAI(Capella-Gutierrez, Silla-Martinez et al. 2009). Trimmed alignments were then concatenated and used to generate the phylogenetic tree by RAxML(Stamatakis 2014). The best substitution model for the full data matrix was determined by the Akaike information criterion in MrModeltest software(Nylander 2004). The best-scoring maximum likelihood (ML) tree was inferred using a novel rapid bootstrap algorithm combined with ML searches following 1000 RAxML runs (using the ‘f -a’ option)(Stamatakis 2014). The divergence times for the species analyzed were estimated by Reltime(Tamura, Battistuzzi et al. 2012). Diverged about 46 million years ago, Ords Kangroo rat is evolutionarily closest to beaver (**Figure 1B**).

### Identification of significant gene expansion in beaver

With the phylogeny that we built and the gene counts from OrthoDB (Zdobnov, Tegenfeldt et al. 2017), we predicted gene family expansion using CAFE 3 (Han, Thomas et al. 2013). CAFE first calculated an error model for gene family size estimation as a part of genome assembly and annotation. For our gene counts, it estimated ~4.8% of the gene families had incorrect gene numbers assigned to them. It corrected this error before calculating ancestral family sizes and then estimated a more accurate gene family evolution rate. Finally, it calculated the probability of observing the sizes of each gene family of those species by Monte Carlo re-sampling procedure. Families with large variance in size, especially observed in closely related species, will tend to have a lower *P*-value. For families with low *P*-values (0.01 as the default), a *P*-value for the transition between parent and child nodes for each branch in the phylogeny was also calculated to identify where the large change of family size takes place.

We identified 84 candidate gene families showing significant expansion in the beaver branch (FDR < 0.01). To sidestep false positives due to assembly and annotation artifact, pseudogenes, etc., we first removed genes without the support of RNA-seq reads from all the tissues (**Supplement Table S1**) in our analysis. We also identified pseudogenes and removed ones with low expression levels (TMP < 5 across tissues). We kept pseudogenes with relatively high expression levels, since they may be functional. We then further filtered genes based on the percentage of identities between beaver gene copies. Briefly, we iteratively removed the gene copy with the least average identity to other beaver genes in the family, if the average identity is below 70%. After this procedure, we reduced our candidate expanded gene families to eight genes (**Figure 2 – Supplement Figure S2**).

### Beaver-specific expansion of conserved genes

To find beaver-specific expansion of conserved genes, we require the same number of copies in other species included in our study and a higher copy number only in the beaver genome. Initially, 234 gene candidates were collected. After removing copies without RNA-seq reads support, lowly expressed pseudogenes (TPM < 5), and gene copies with low identity between gene products (identity < 0.7), 83 genes remained. We further processed them to remove potential false positives by considering only genes with the following qualifications: (1) gene predictions with good synteny in their genomic neighborhood among beaver, mouse, and human; (2) genes with differences in the coding sequences of their copies; (3) genes with variable sites in their copies supported by RNA-seq reads (from the same individual beaver). 18 beaver-specific gene expansions met these stringent criteria and were further validated by qPCR, with five of them show more copy numbers comparing to reference genes.

### Pseudogene identification

GeneWise(Birney, Clamp et al. 2004) was used to identify pseudogenes (**Figure 2 – Supplement Figure S5**). For a predicted beaver gene, we extracted the genomic sequence from its locus with both upstream and downstream 5-kb regions. Using its mouse (or human) ortholog as the reference, we then scanned the gene sequence by GeneWise and checked the presence of frameshift indels that can ‘pseudogenize’ the gene. We also checked if the predicted gene can be a processed pseudogene, which can be generated through mRNA retrotransposition. If a predicted beaver gene has no introns, we checked its orthologs in other rodents to determine whether it is a processed pseudogene or a single-exon gene.

### Positive selection

Branch-site likelihood method (Zhang, Nielsen et al. 2005) implemented in PosiGene (Sahm, Bens et al. 2017) was used to identify genes under positive selection in the beaver genome. Only 9,750 genes with alignment of ortholog coding sequence from at least 10 species were considered. After the initial prediction by PosiGene (Sahm, Bens et al. 2017), manual checking was carried out for all ~150 candidate positive selection genes (*P* < 0.05). Briefly, we removed false positives, and for genes with ambiguous signals we manually checked and improved their predicted gene structures based on protein sequence alignments of orthologs and supporting RNA-seq reads using Apollo (Lee, Helt et al. 2013). We then used the improved coding sequences as the input for PosiGene and ran the pipeline three times. Finally, we identified 21 beaver genes under positive selection with FDR < 0.01 consistently in all three independent runs. The coding sequences of other species were download from NCBI as of January of 2020.

### RNA extraction and sequencing

Total RNA was isolated from frozen tissues (brain and liver with two replicates each) using the mRNA-Seq Sample Prep Kit Illumina (San Diego, CA. USA) in accordance with the manufacturer’s instructions, and the mRNA integrity was checked by the agarose gel analysis. Polyadenylated RNA was then isolated using a poly-dT bead procedure and followed by reverse transcription. Short-insert ‘paired-end’ libraries were prepared using the Illumina TruSeq Sample Preparation Kit v2, and the sequencing was performed on the Illumina HiSeq2000 platform. The raw data were processed by NGS QC Toolkit (v.2.3.3)(Patel and Jain 2012) to remove low-quality reads.

### De novo transcriptome assembly

RNA-seq data from brain and liver (each with two replicates) were assembled using Trinity(Haas, Papanicolaou et al. 2013). We collected 16,816 high quality beaver transcripts, by requiring each transcript to meet the following conditions: (1) proper start and stop codons, (2) a correct reading frame (codons in triplets), (3) a gene length similar to the mouse ortholog (± 20%), (4) a good alignment with the mouse ortholog (in peptide sequence), (5) the best candidate among all beaver Trinity assembled sequences.

### qPCR validation

Candidate primers were designed by Primer-Blast (Ye, Coulouris et al. 2012) in the exons of selected genes. Specificity of the primers was first checked by Primer-Blast against all of the coding sequences in the beaver genome. Then genome-wide specificity was checked by MFEprimer (Qu, Zhou et al. 2012). Specificity of primers was further inspected by gel electrophoresis and the melting curve analysis. Amplification efficiency was checked by a standard curve analysis for each candidate primers with different amount of DNA input to make sure that the finally used primers have equal amplification efficiency. Quantitative PCR was used to quantify gene copy number, with three replicates for each of the three different amount of DNA input. Two genes – *Hcfc1* on an autosome and *Pelo* on chromosome X – with no predicted expansion in beaver and no annotated expansion in other rodents (according to orthoDB (Zdobnov, Tegenfeldt et al. 2017)) were used as references. The copy number of each target gene relative to its corresponding reference genes was calculated by Δ*C*_*t*_. And the genomic DNA of a male beaver, different from ones whose samples were used for the genome sequencing and assembly and transcriptomics, was used for the experiment. Primers for target and reference genes are listed in **Supplement Table S2**.

### Western blot

Liver cytosolic extracts were quantitated using the BCA assay (Thermo). 30 ug of each extract was resolved through 4-20% Criterion Tris-Glycine (TGX) Stain-Free SDS-PAGE (Biorad) and transferred to nitrocellulose. Prior to transfer, total proteins were imaged using a Biorad Gel Documentation System to control for protein loading. Rabbit polyclonal anti-Aldh1a1 (Invitrogen cat# PA5-95937) was used as the primary to detect Aldh1a1 isoforms. Protein loading was also checked by performing Western blot using anti-beta Actin (Abcam, ab8227).

### Ethanol treatment of cells

The effects of ethanol on cellular functions were studied with cultured primary beaver and mouse lung fibroblasts. We used two different beaver and mouse cell lines, low PD cells stabilized in culture. Cells were seed in complete medium containing 15% FBS in 96 well plate for 24 hours, then incubated in medium containing 300 mM ethanol for seven hours. Then WST-1 assay was performed by adding WST-1 reagent (cell proliferation reagent, Roche) directly to the culture wells, incubating for 4h at 37°C and 5% CO, shaking thoroughly for 1 min on a shaker and then measuring the absorbance at 430-480 nm with TECAN spark 20M spectrophotometric reader. The stable tetrazolium salt WST-1 was cleaved to a soluble formazan by a complex cellular mechanism that occurs primarily at the cell surface. This bio-reduction is largely dependent on the glycolytic production of NAD(P)H in viable cells. Therefore, the amount of formazan dye formed directly correlates to the number of metabolically active cells in the culture.

To examine whether beaver and mouse lung fibroblast cells show significantly different rate of cell death under the ethanol treatment, relative to corresponding controls, a linear mixed effects model was used with species as the fixed effect and individuals (*n*=4 for each species with 3 replicates each individual) as random effects (lmer(cell death rate ~ Species+(1|individual))), which is implemented in the R package lme4 (Bates, Mächler et al. 2015).

### Liver aldehyde dehydrogenase activity

In addition to the cytosolic *Aldh1a1*, a mitochondrial enzyme, *Aldh2*, also plays an important role in the aldehyde metabolism. A previous study had shown that in murine liver *Aldh1a1* plays a more important role in the metabolism of endogenous aldehydes than *Aldh2* (Makia, Bojang et al. 2011). To exclude influence from *Aldh2*, we tested aldehyde dehydrogenase activity using cytosolic protein extracts from liver cells. Crude cytosolic protein extracts were prepared from wild beaver or mouse (C57/Bl6) livers using phosphate buffer similar to a previously described method (Makia, Bojang et al. 2011). Specifically, tissue was resuspended in K-Phos Buffer (50 mM potassium phosphate pH7.4, 250 mM sucrose, 1 mM EDTA) at 1g/3.0ml. Tissues were dounce homogenized in ice using a Teflon pestle followed by ten passages through a 27Ga needle to lyse the cells. Samples were centrifuged (Thermo/Sorvall Legend Micro21R) at 1000 rpm (100x g) for 10 minutes at 4°C to remove any remaining intact cells. Supernatant was transferred to a clean tube and centrifuged at 14, 000 rpm (18,800x g) for 20 minutes at 4°C. We found that the supernatants/extracts prepared this way and rapidly aliquotted and frozen with liquid nitrogen did not lose significant activity after one freeze/thaw cycle. Just prior to assaying, a 100 μl aliquot of the supernatant was thawed on ice and passaged through a 0.5 ml Zeba desalting spin column (Thermo; 7,000 mwco) equilibrated in the same K-Phos buffer to remove small molecules that might compete with added substrates. Aldehyde dehydrogenase was measured in 384-well transparent microplates in 50 μl volume consisting of 10 μl extract in the same K-Phos buffer supplemented with 1mM NAD^+^. Aldehyde substrates all-trans-retinal (RET), malonaldehyde tetrabutylammonium salt (MAD), and 4-hydroxynonenal (NHE) were added at indicated concentrations from DMSO stocks. Final DMSO in assay was 4%. Dehydrogenase activity was measured as the change in absorbance at 340 nm (reduction of NAD^+^ to NADH) using a Tecan Spark 20M plate reader pre-equillibrated at 37°C. Rates were determined after a few minutes of lag during which time the plate temperature was adjusting to 37°C. NAD+, all-trans-retinal, and malonaldehyde tetrabutylammonium salt, were purchased from Sigma. 4-hydroxynonenal was purchased from Cayman Chemical. Aldh1a1-class of specific activity was determined by subtracting any trace amount of NAD+ to NADH conversion that occurred in the absence of added aldehyde substrate.

### Gene expression comparison between beavers and mice

For our transcriptomic analysis, we used beaver samples from two young male adults and RNA-seq data from 6 weeks old young adult mice (Li, Qing et al. 2017). The RNA-Seq data of mice were generated from eight individuals (4 females and 4 males) for both liver and brain. Although our beaver samples were from two males, we included both male and female mice to increase the detection power with sex as a factor in the model for differential gene expression analysis. Principle component analysis showed good separation between beaver and mouse samples (**Figure 6 – Supplement Figure S14**). We only considered protein coding genes and used Kallisto (Bray, Pimentel et al. 2016) to estimate gene expression levels, which showed better performance for genes with paralogs.

The detection of differential expressed genes between species is much more complicated than analysis within species. Because several differences between two genomes – e.g., different gene annotation quality, different number of genes, and different lengths of same genes between species – can lead to biased results, we used the following strategies together in our analysis. (1) For gene annotation quality control, we only considered 16,303 beaver genes that show high protein sequence similarity with their corresponding orthologs from other rodent species (including mouse) and could be successfully clustered into gene families by OthorDB (Zdobnov, Tegenfeldt et al. 2017). (2) We further restricted our analysis to genes (12,089) that share one-to-one orthology between beaver and mouse. (3) We also controlled gene length. We used tximport (Soneson, Love et al. 2015) to obtain family-level gene expression and weighted gene lengths for each gene family. Together with estimated read counts, the weighted gene lengths were also provided to DESeq2 (Love, Huber et al. 2014), and differences in gene lengths among samples were considered for differential expression analysis.

### ALDH1A1 expression during human aging

Using data from the GTEx Project (phs000424.GTEx.v7.p2.c1) (Carithers, Ardlie et al. 2015), we studied gene expression changes of ALDH1A1 during aging. In the GTEx Project, many samples were taken from post-mortem individuals. Studies have demonstrated that expression of some genes may change in certain tissues after death and show significant association with post-mortem interval (PMI, in minutes between death and sample collection) (Ferreira, Munoz-Aguirre et al. 2018). To reduce artifacts from PMIs, we ignored tissues where the expression of ALDH1A1 shows significant association with PMI (P < 0.01). For the remaining tissues, we used liner regression to assess the association between gene expression and age with sex, body mass index, and PMI as covariates.

## Data Analyses and Availability

Genome DNA sequence data were submitted to BioProject database (https://www.ncbi.nlm.nih.gov/bioproject/) with accession numbers PRJNA505050. Genome assembly was submitted to the NCBI Assembly database (https://www.ncbi.nlm.nih.gov/assembly/) with accession number RPDE00000000. RNA-Seq data of liver and brain tissue of two beaver individuals were submitted to NCBI with accession number PRJNA627298, which will be released once the manuscript is accepted.

## Acknowledgments

This work was supported by NIH grant P01AG047200 to V.G., A.S., V.N.G., J.V., and Z.D.Z.

## Additional files

*Supplementary file 1*. Coding sequences and protein sequences alignments with selection signals of beaver genes under positive selection (FDR<0.01).

---

**Figure S1. Related to Figure 1.**
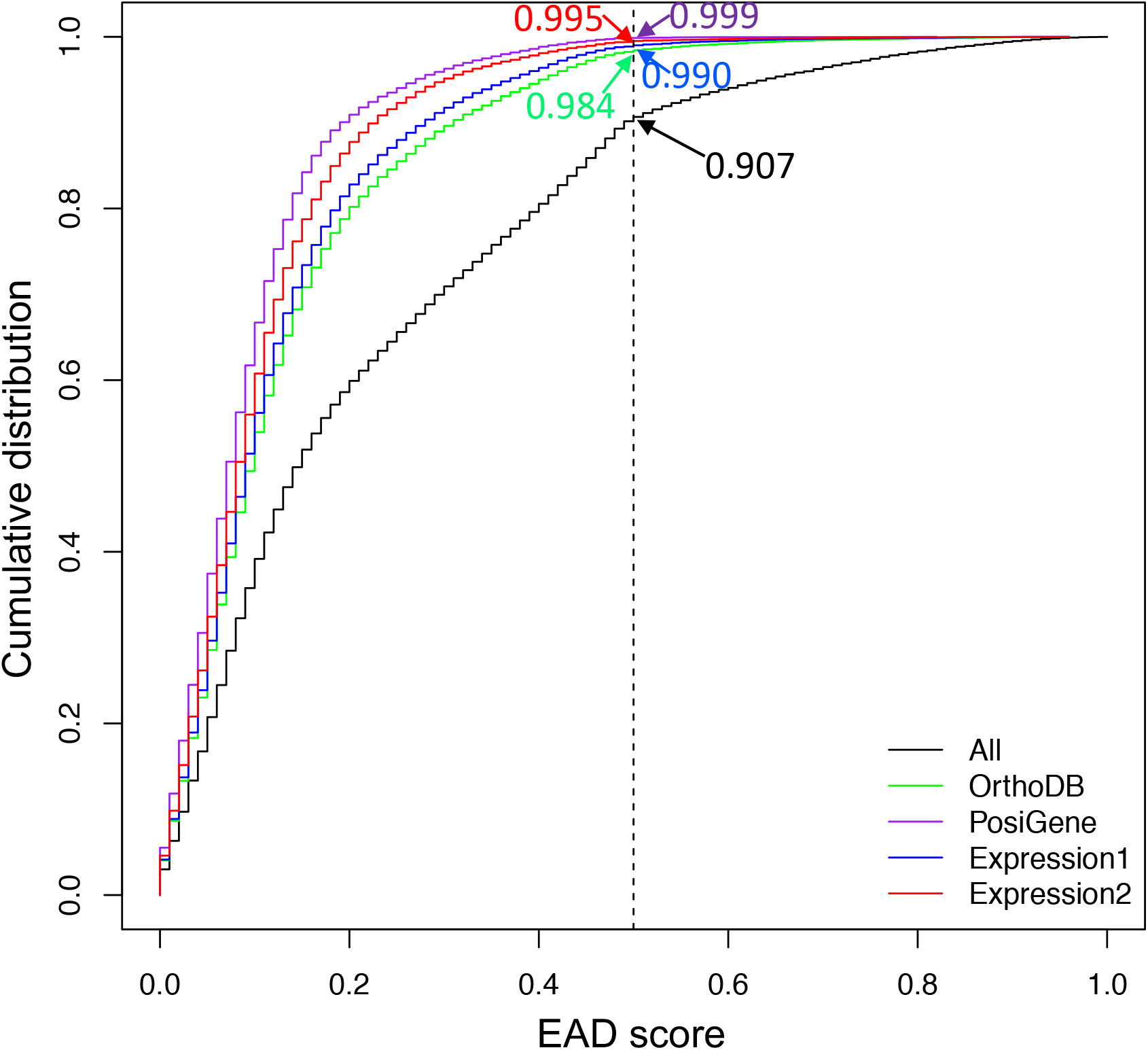
Cumulative Distribution of AED score. All: 26,515 predicted genes; OrthoDB: 17,661 genes grouped into gene families by mapping beaver proteins to OrthoDB(Zdobnov, Tegenfeldt et al. 2017); PosiGene: 9,750 genes with coding sequences of at least 10 species in the alignment for identification of genes under positive selection; Expression1: 16,303 beaver genes with mouse orthologs; Expression2: 12,089 beaver genes with one-to-one mouse orthologs. An AED (Annotation Edit Distance) score measures the goodness of fit of each gene to the evidence supporting it. The dashed line shows the EAD score = 0.5. A genome is considered well annotated if more than 90% genes have AED scores < 0.5(Campbell, Holt et al. 2014).

**Figure S2. Related to Figure 2.**
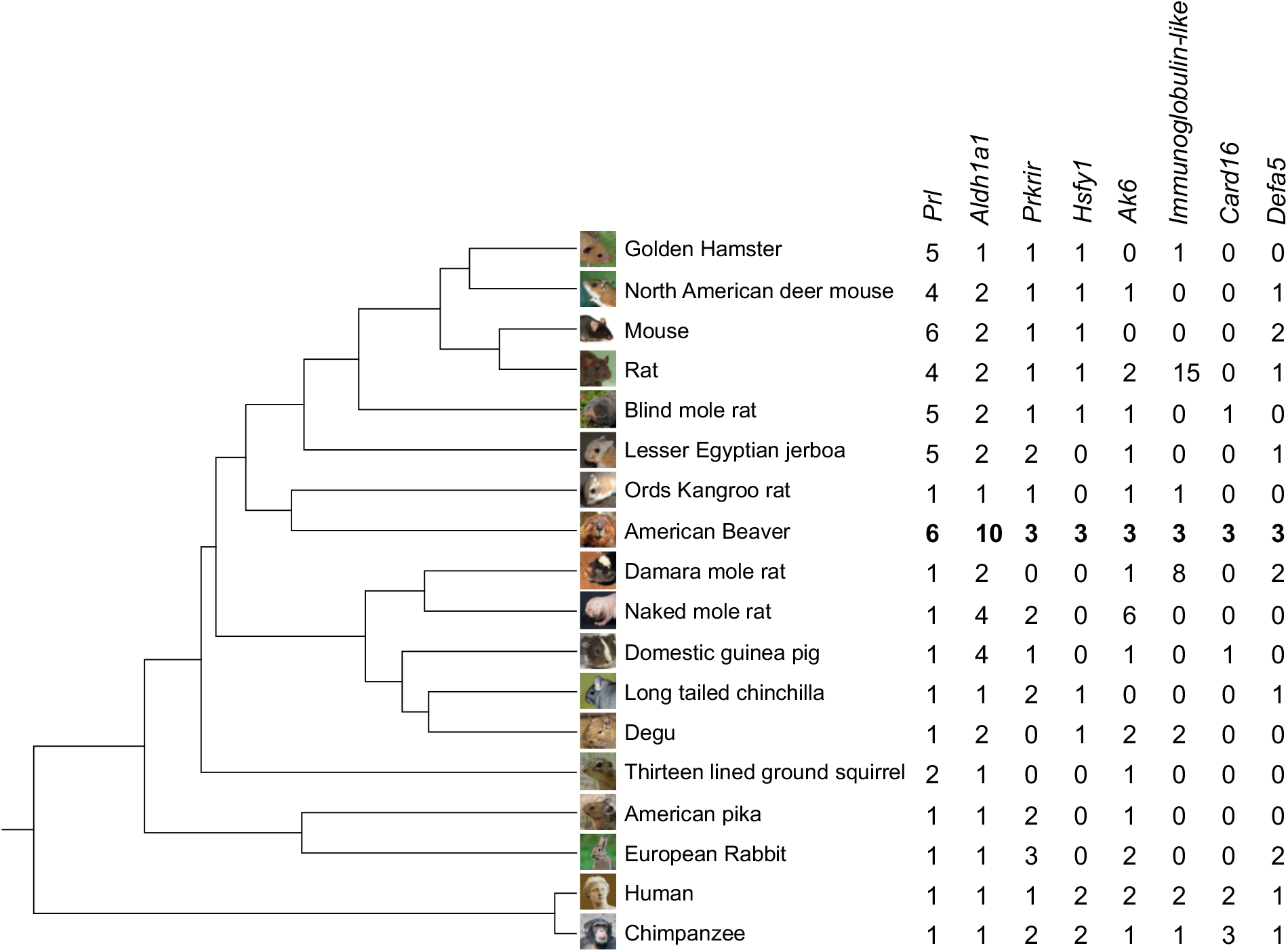
Gene families with significant expansion in beavers. *Prl*: prolactin; *Aldh1a1*: aldehyde dehydrogenase 1 family member A1; *Prkrir*: 52-KDa repressor of the inhibitor of the protein kinase; *Hsfy1*: heat shock transcription factor Y-linked 1; *Ak6*: adenylate kinase 6; *Immunoglobulin-like*: no gene name but with Immunoglobulin-like domain; *Card16*: caspase recruitment domain family member 16; *Defa5*: defensin alpha 5.

**Figure S3. Related to Figure 2.**
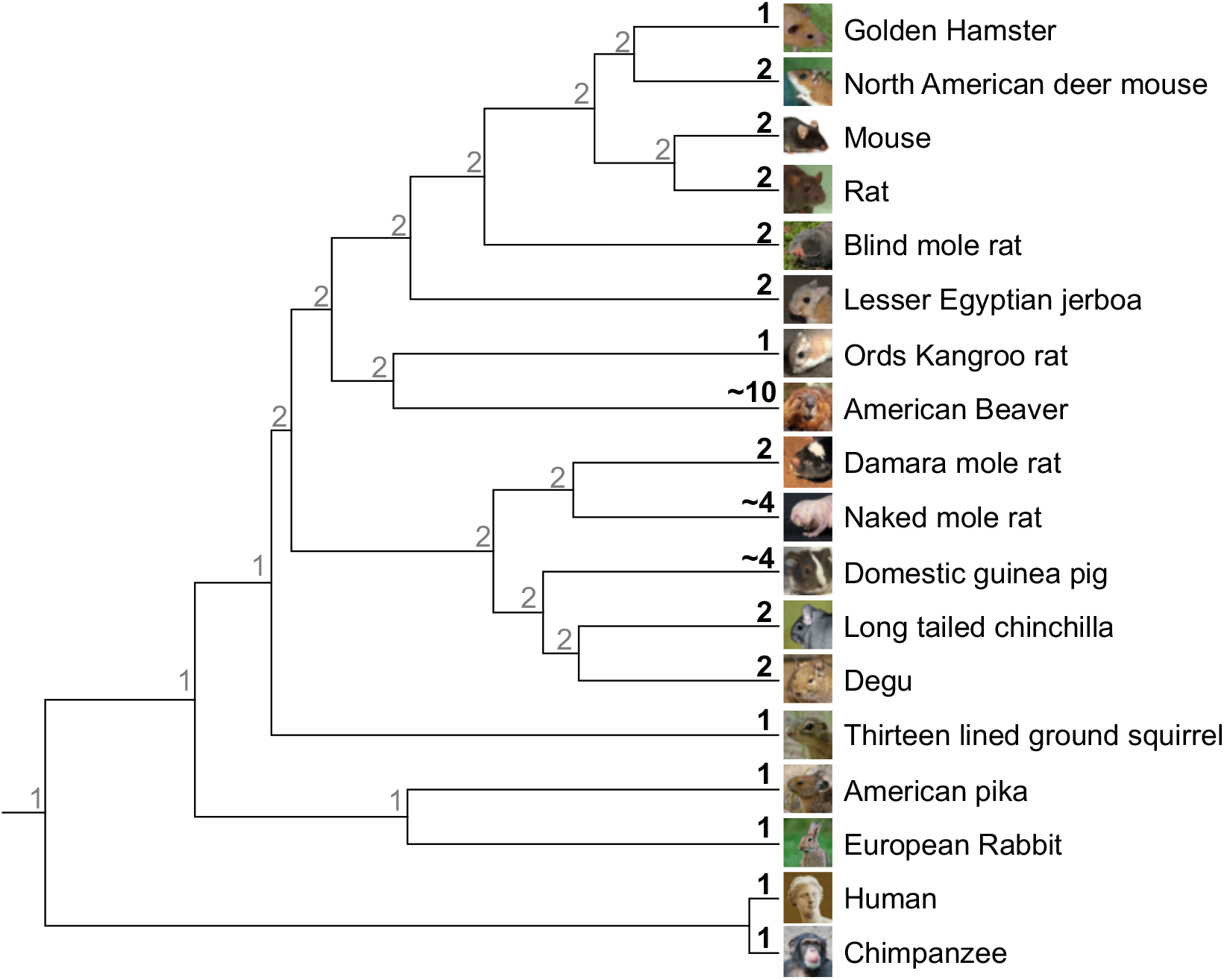
Phylogenetic tree with *Aldh1a1* copy numbers. Shown on the leaf and the ancestral nodes are the copy numbers (in black) of *Aldh1a1* in species annotated by this study and OrthoDB(Zdobnov, Tegenfeldt et al. 2017) and ones (in gray) estimated by CAFE3(Han, Thomas et al. 2013), respectively.

**Figure S4. Related to Figure 2.**
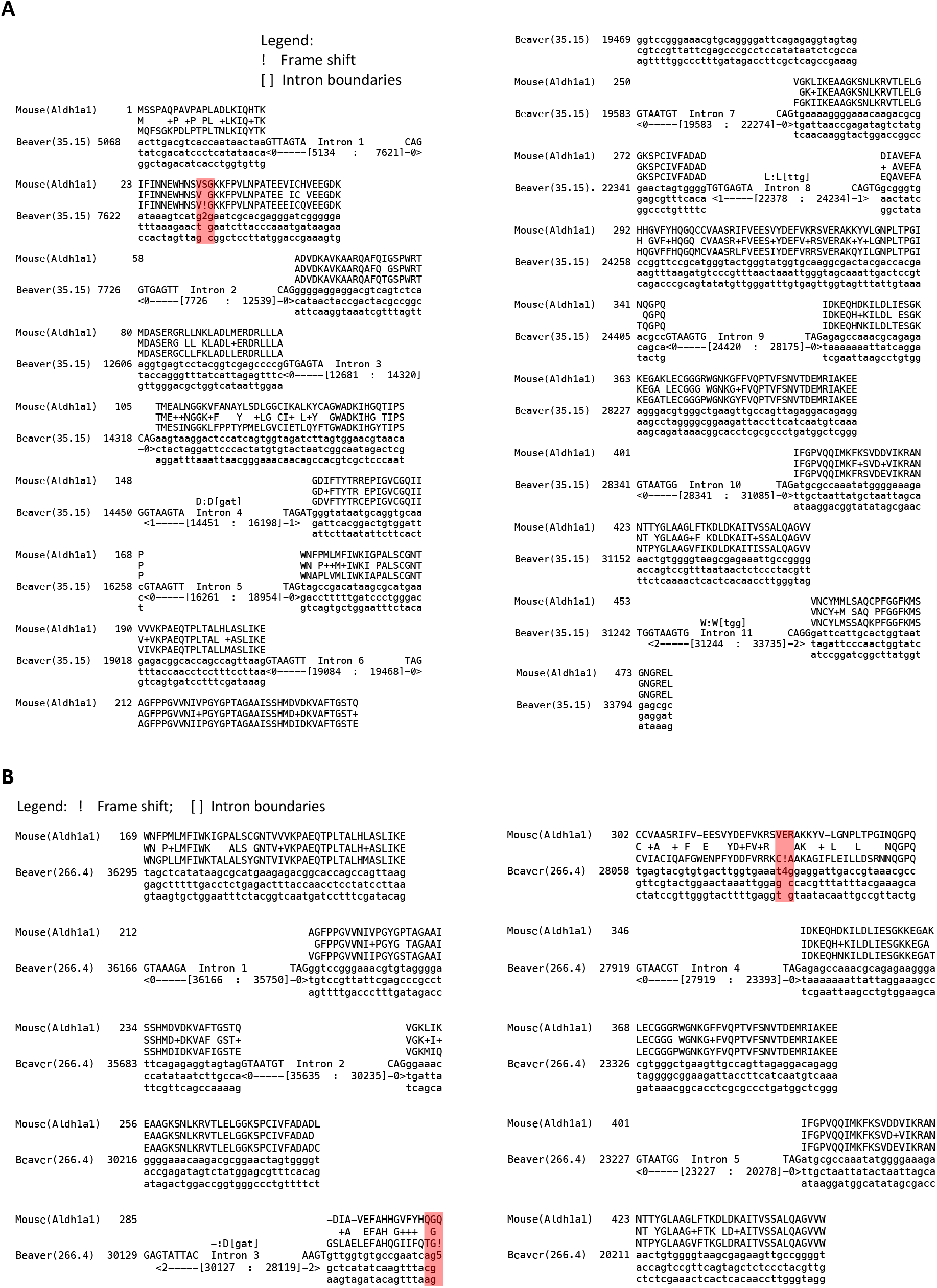
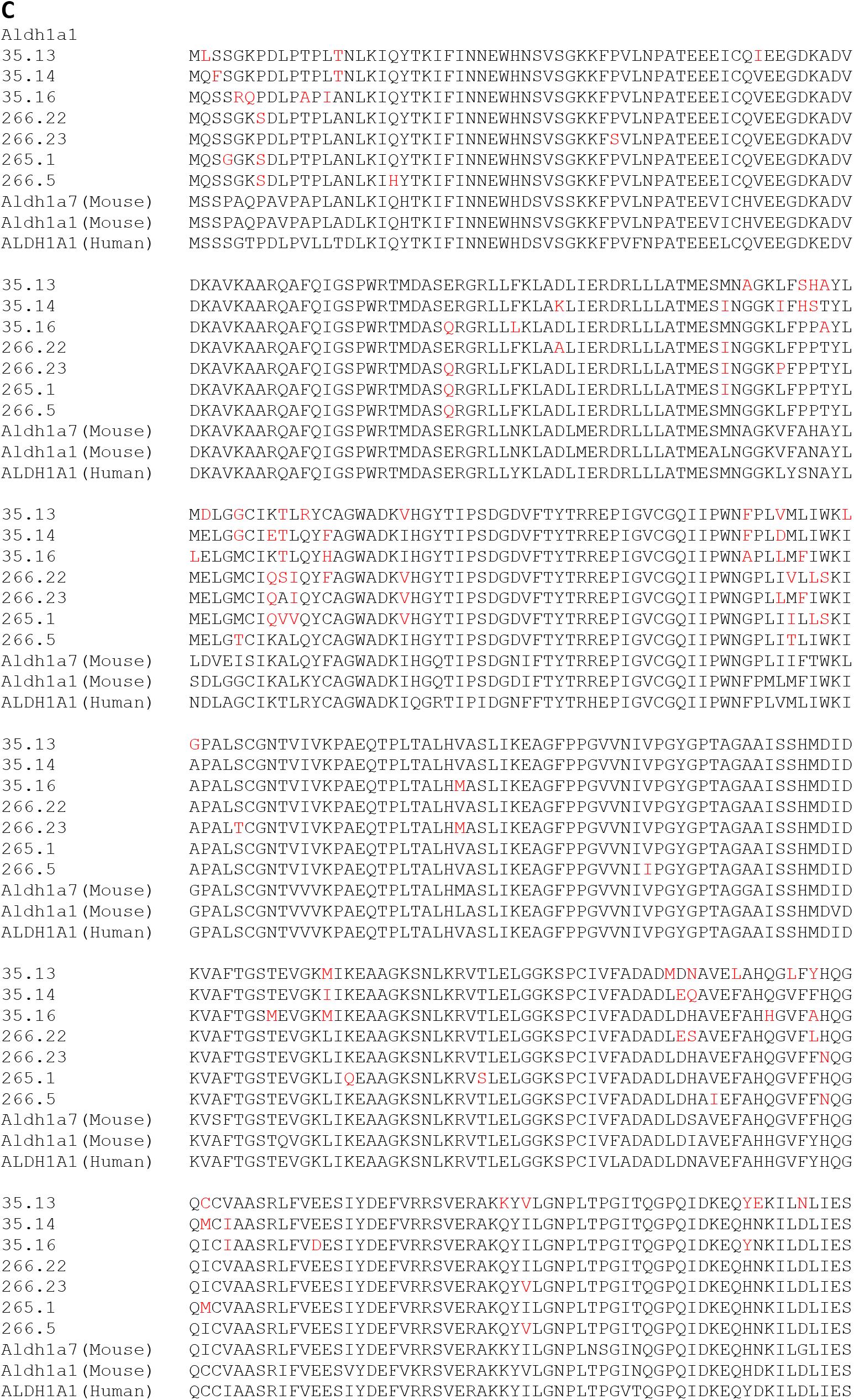
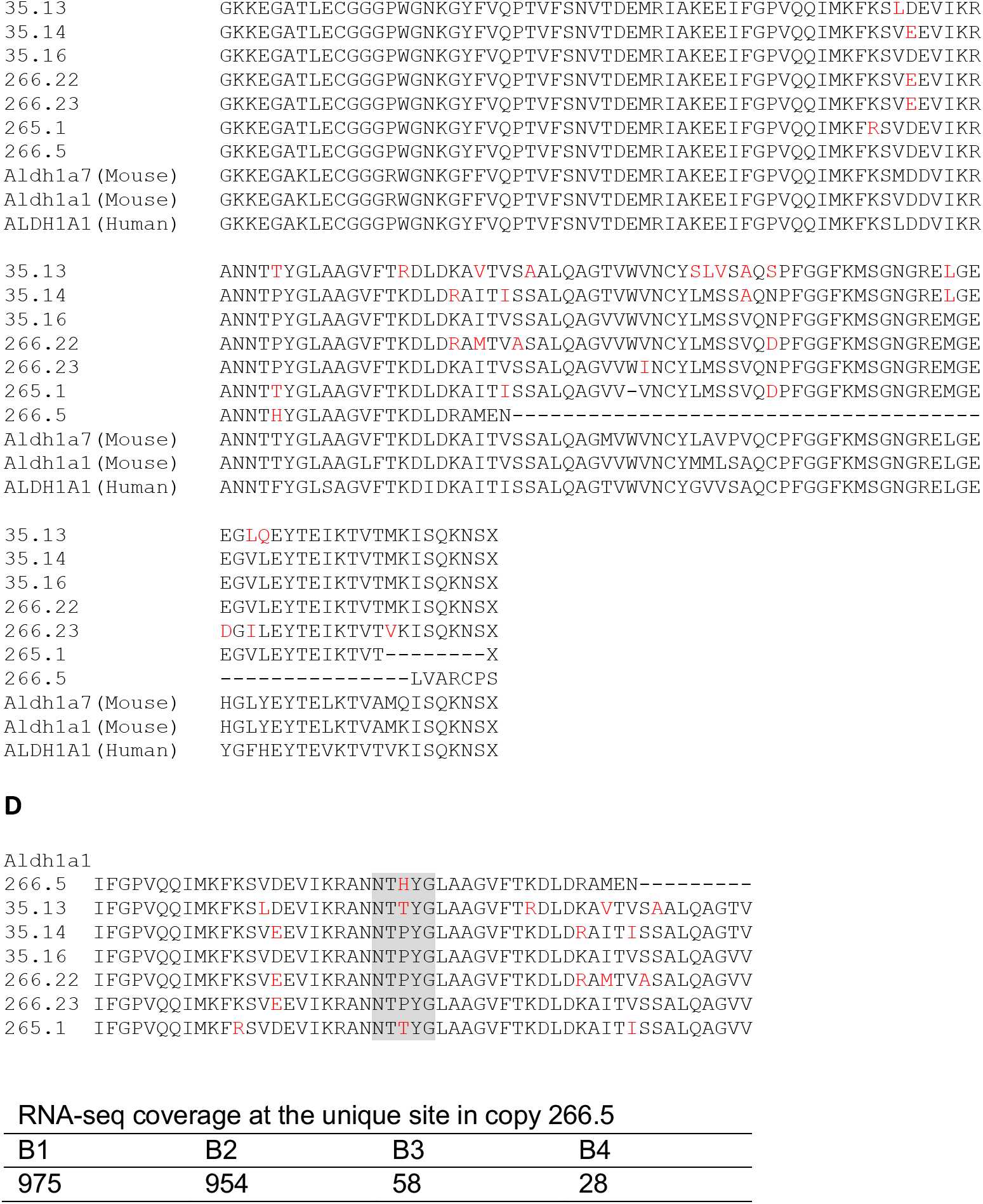
Predicted beaver *Aldh1a1* copies. Two copies are likely pseudogenes. Gene structures of (**A**) copy 35.15 with a frame-shift indel (highlighted in red) and (**B**) copy 266.4 with two frame-shift indels. (**C**) Multiple sequence alignment of beaver Aldh1a1 with of mouse and human orthologs. Amino acid residues in red show divergent evolution among different copies in the beaver genome. (**D**) RNA-seq coverage at the unique amino acid residue in beaver *Aldh1a1* 266.5. The alignment of peptide sequences encoded by exon 11 is shown. Variable sites were highlighted in red, and the gray box indicates the selected sites where we checked the coverage of RNA-seq reads. The amino acid residue in copy 266.5 is different from the ones in all other copies. The table below shows the RNA-seq read coverage at genome region corresponding to this unique amino acid residue based on RNA-seq data from each of four beaver individuals (see **Supplement Table S1**).

**Figure S5. Related to Figure 2.**
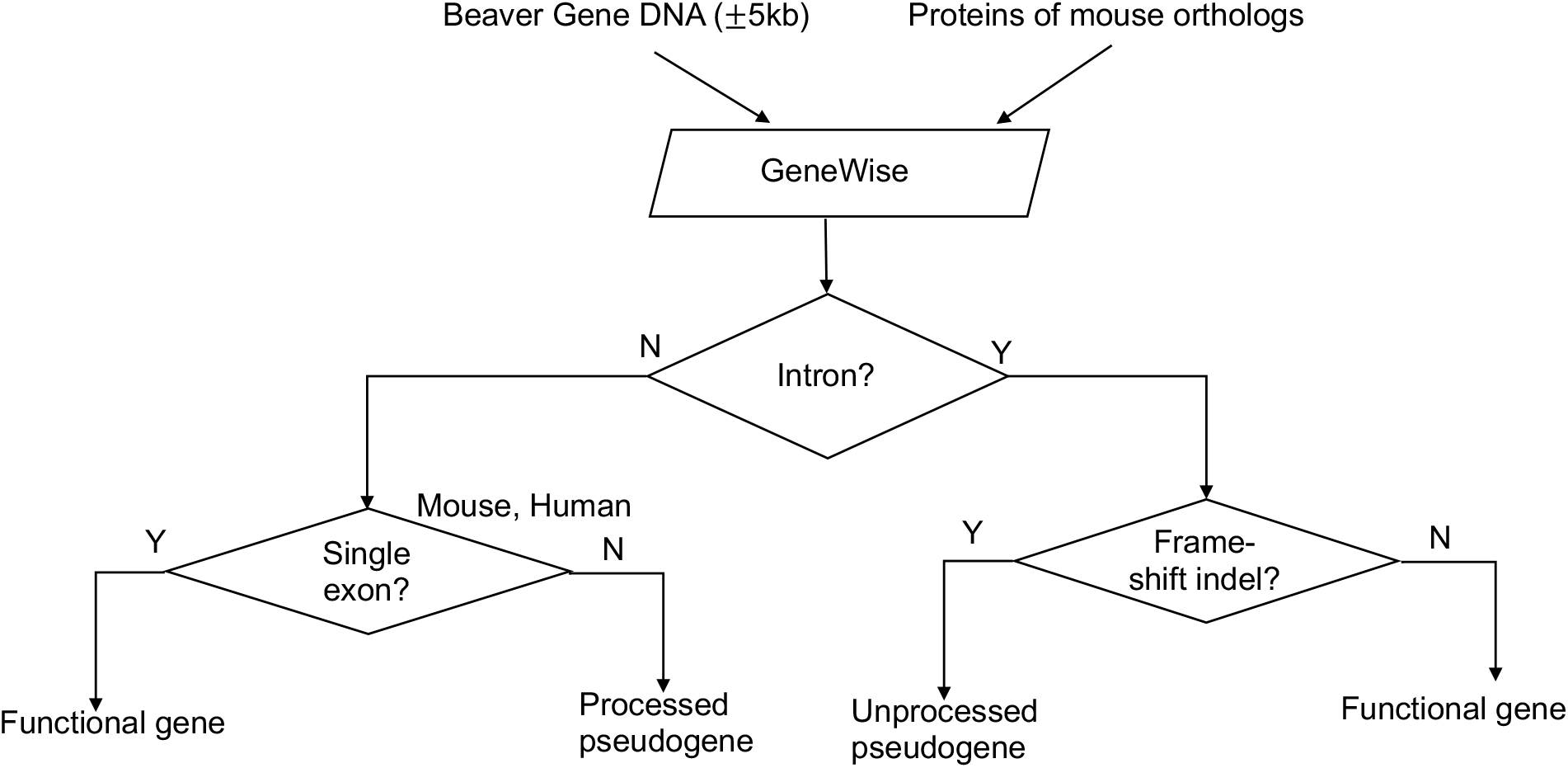
Pseudogene identification pipeline.

**Figure S6. Related to Figure 2.**
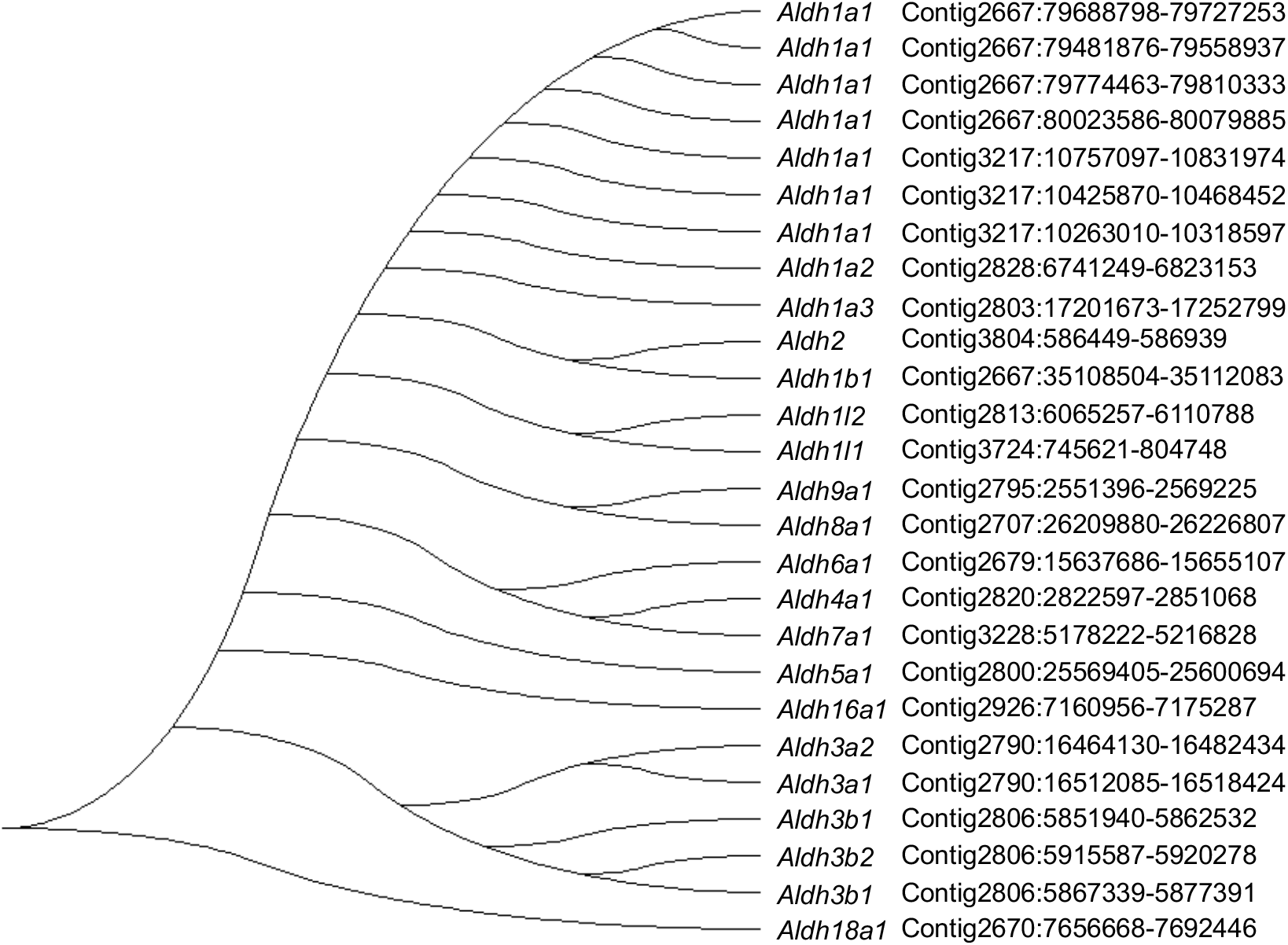
Phylogenetic tree of beaver *Aldh*. The genome coordinates of each gene are given after the gene names.

**Figure S7. Related to Figure 2.**
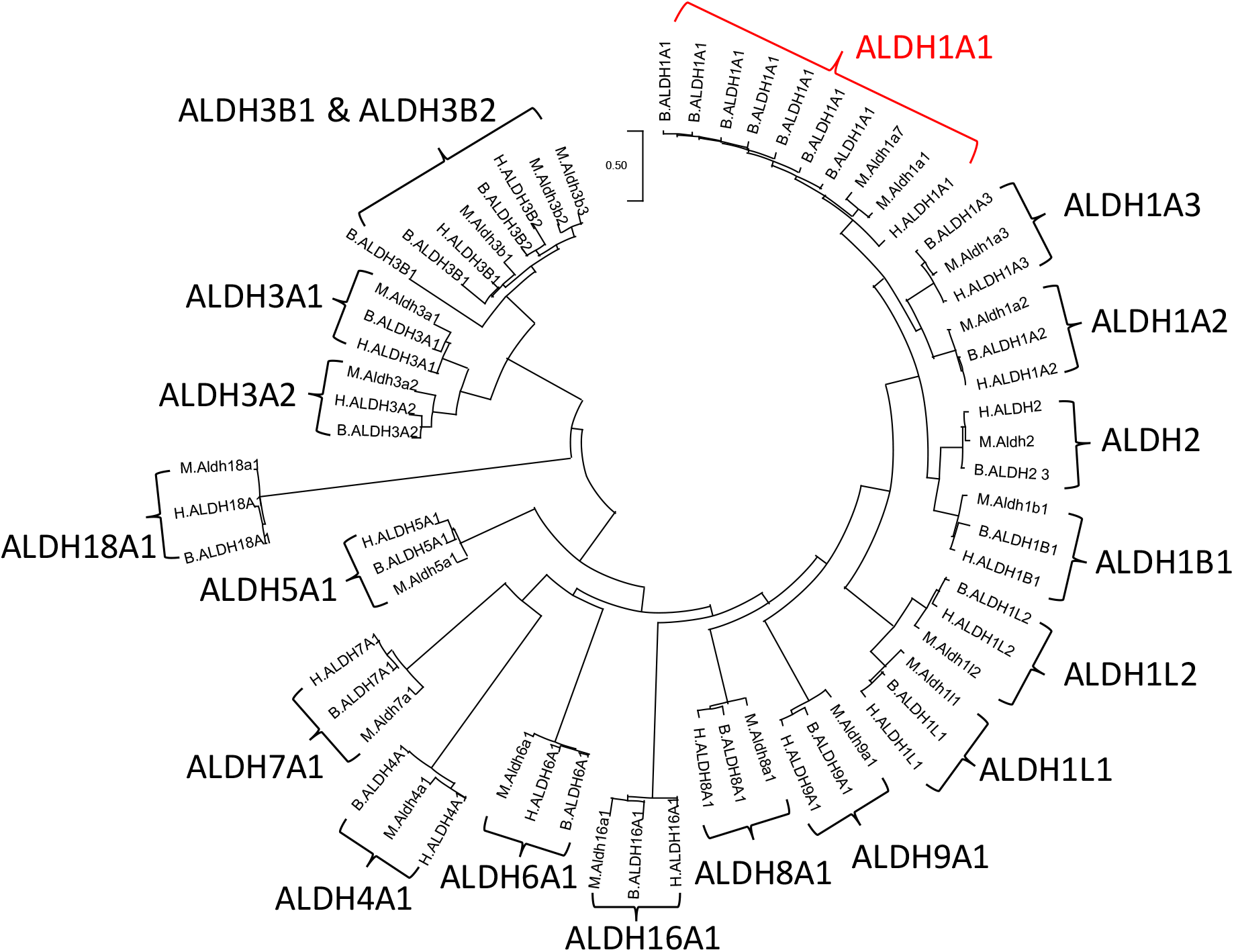
Phylogeny of beaver, mouse, and human *ALDH*. The letter prefix of the gene name denotes the species: B for beaver, M for mouse, and H for human. Gene annotation of each beaver *Aldh* gene was manually checked, and manually editing was done by Apollo(Lee, Helt et al. 2013), according to RNA-Seq reads alignment, when it was necessary.

**Figure S8. Related to Figure 4.**
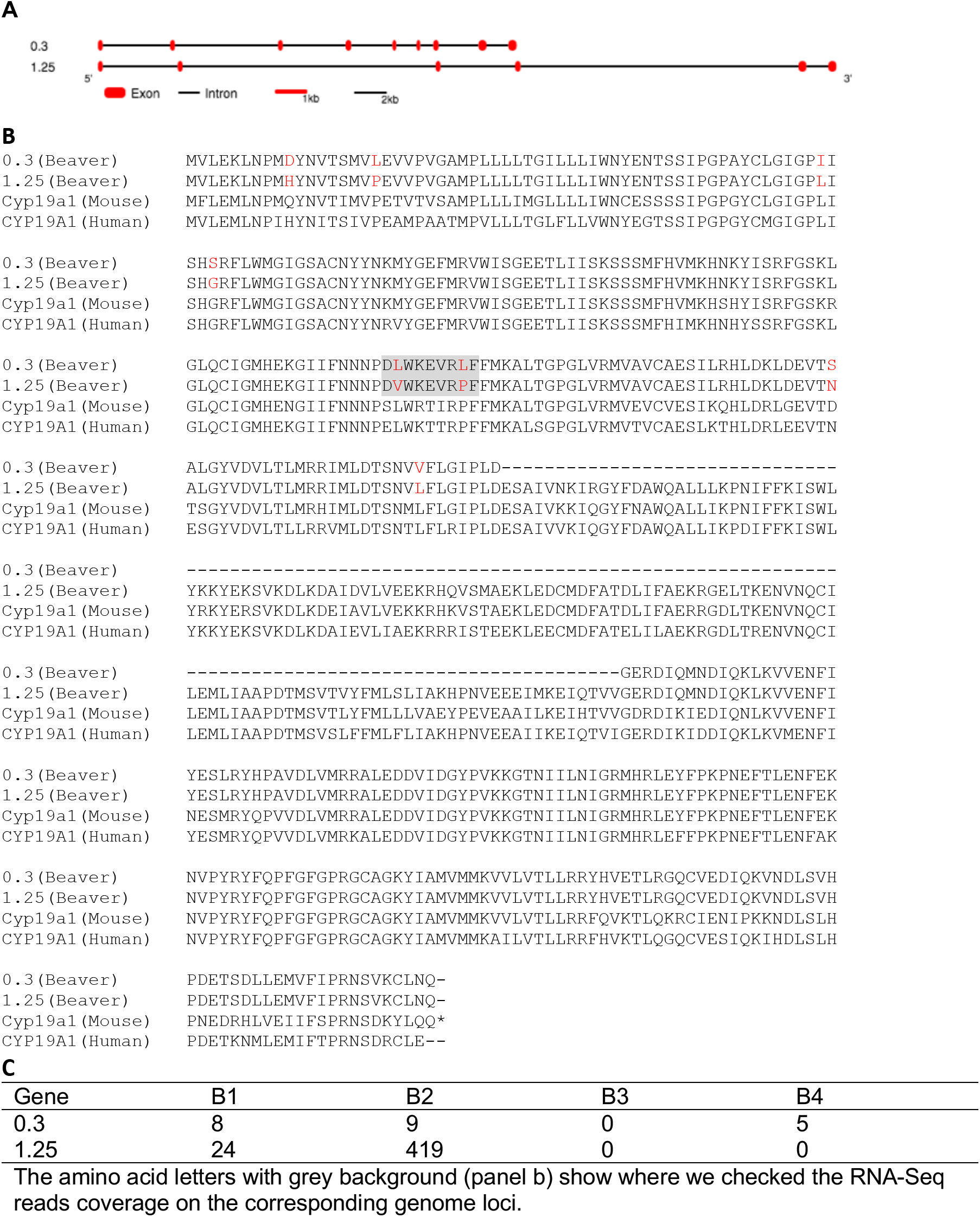
Duplication of Cyp19a1. **A**) Gene structure of beaver Cyp19a1. **B**) Protein sequence alignment. Sequences with grey background show where we check the RNA-Seq reads coverage. **C**) RNA-Seq reads coverage at unique sites.

**Figure S9. Related to Figure 4.**
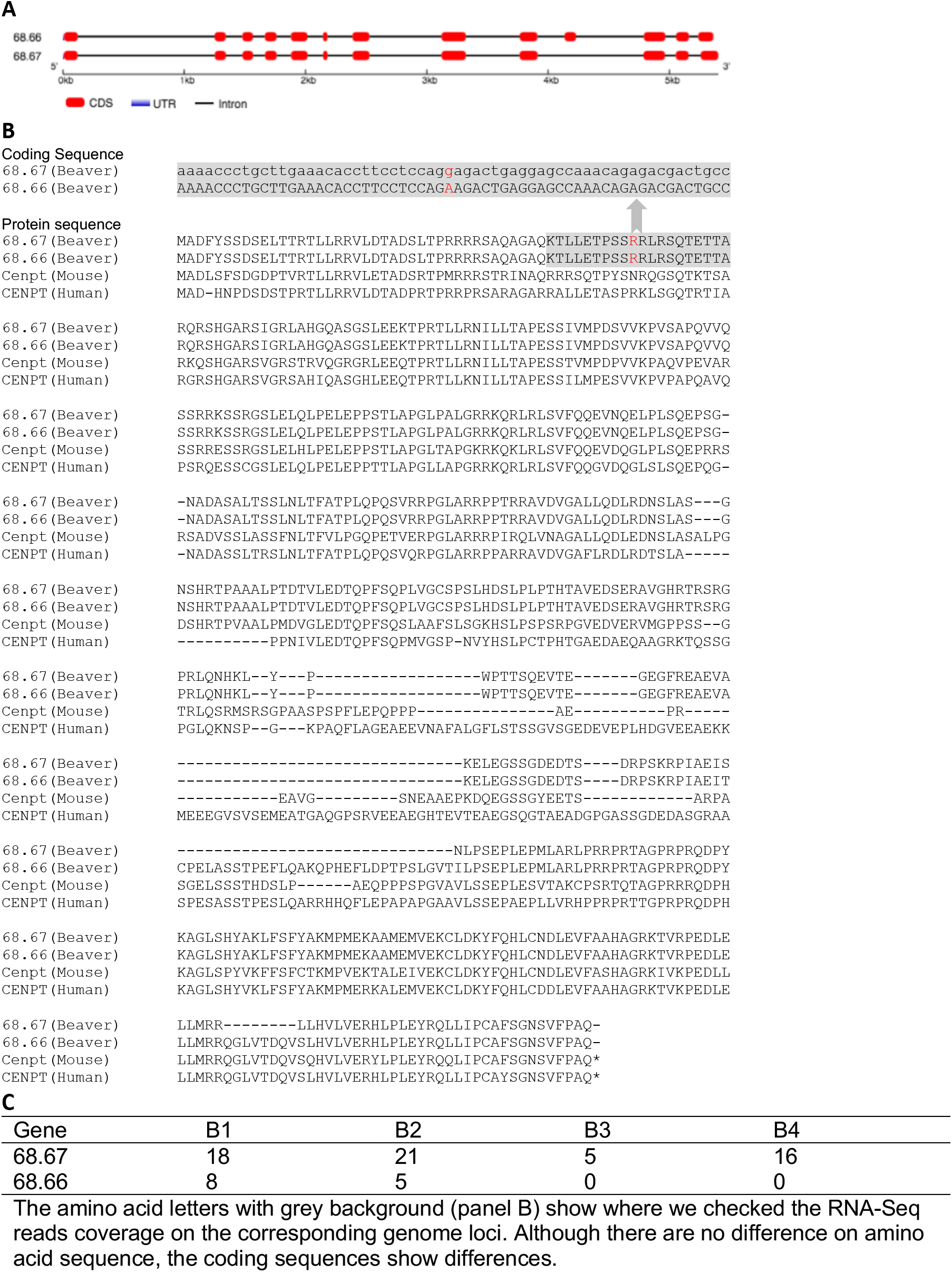
Duplication of *Cenpt*. **A**) Gene structure of beaver *Cenpt*. **B**) Protein sequence alignment. Sequences with grey background show where we check the RNA-Seq reads coverage. **C**) RNA-Seq reads coverage at unique sites.

**Figure S10. Related to Figure 4.**
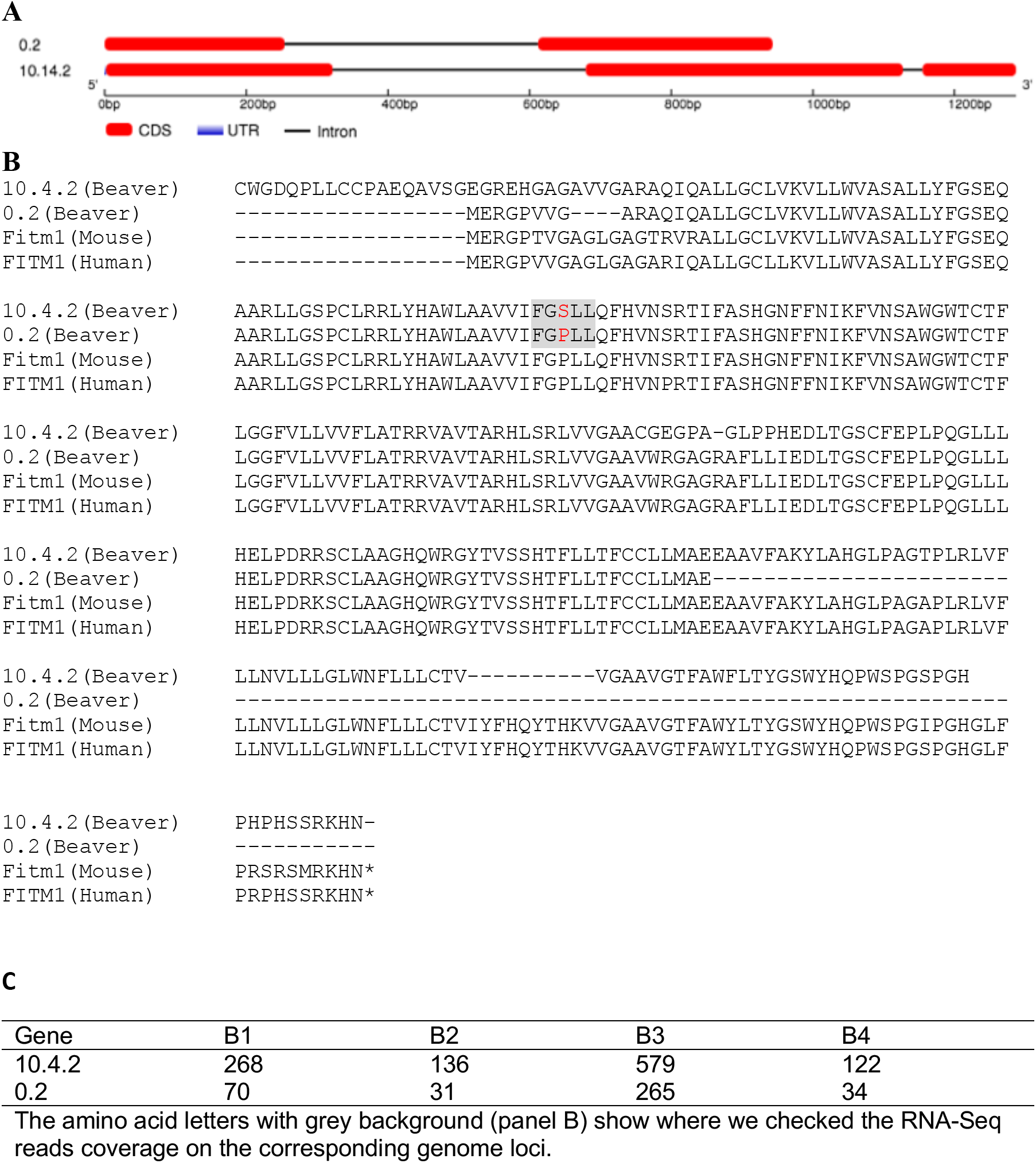
Duplication of *Fitm1*. **A**) Gene structure of beaver *Fitm1*. **B**) Protein sequence alignment. Sequences with grey background show where we check the RNA-Seq reads coverage. **C**) RNA-Seq reads coverage at unique sites.

**Figure S11. Related to Figure 4.**
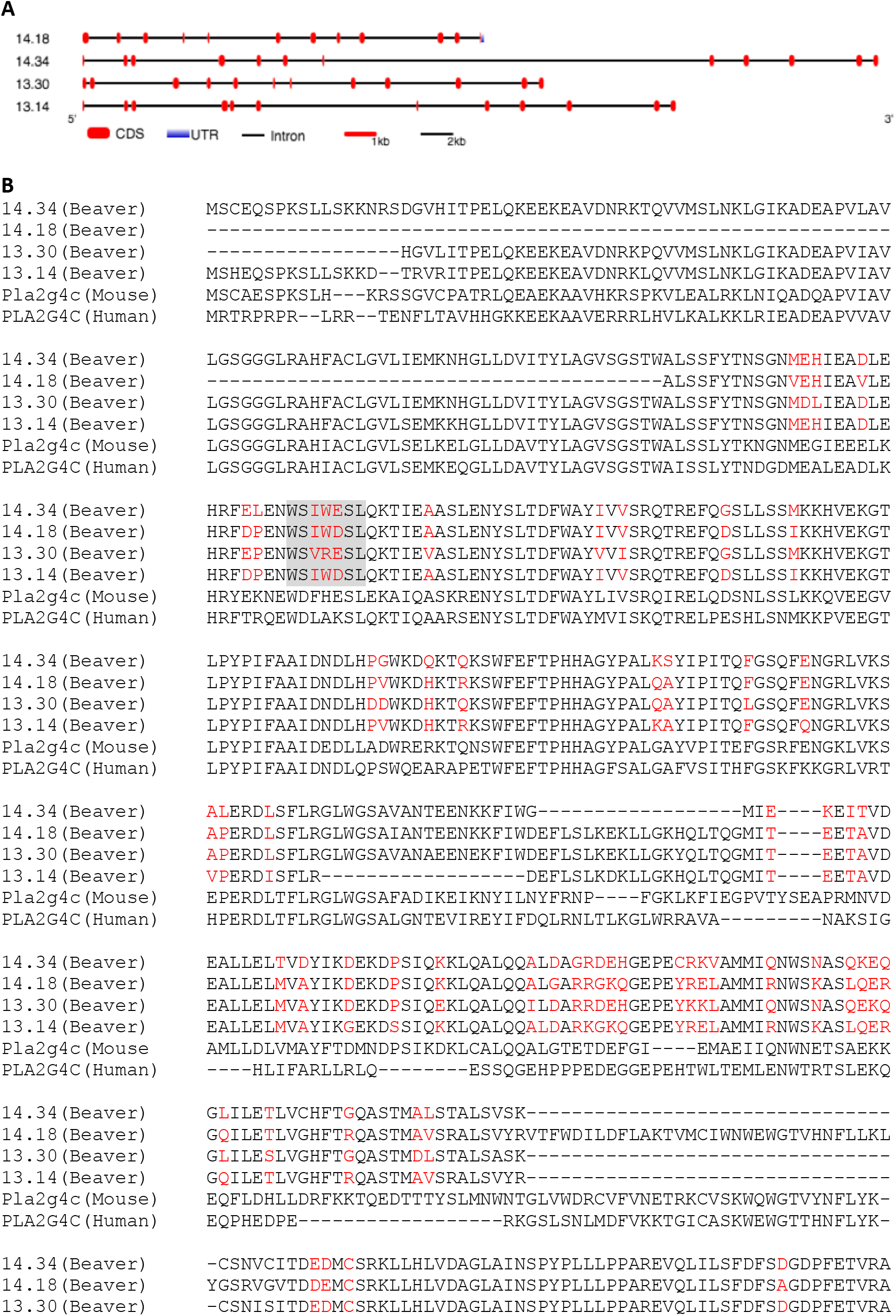
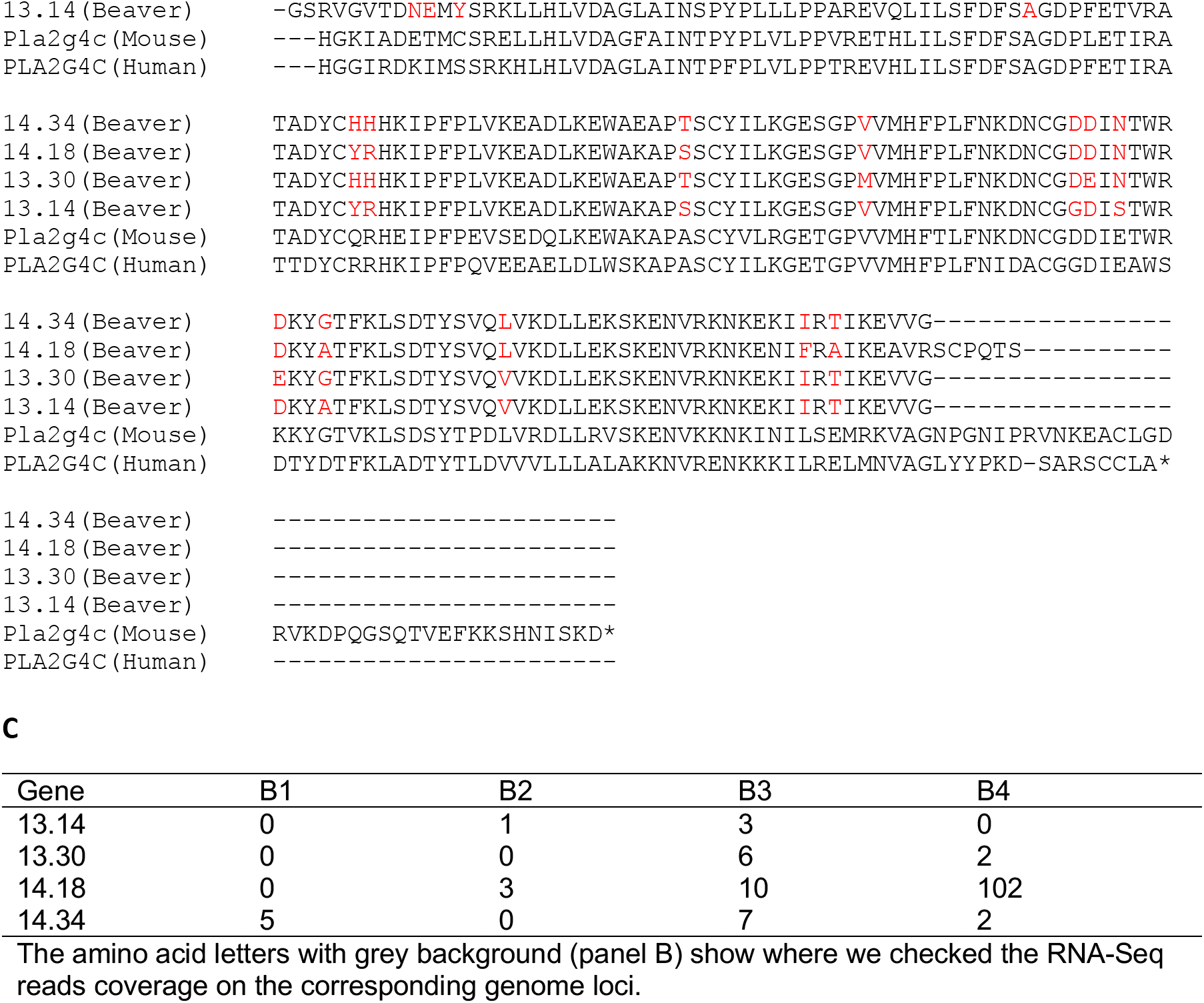
Duplication of *Pla2gc*. **A**) Gene structure of beaver *Pla2gc*. **B**) Protein sequence alignment. Sequences with grey background show where we check the RNA-Seq reads coverage. **C**) RNA-Seq reads coverage at unique sites. Consistent with genome sequencing result and qPCR also showed there are duplication of *Pla2gc* in beaver genome (**Figure 4A**). However, we did not observe 4 times signals by qPCR. It may be because the primer we designed do not perfectly match sequences from each of those copies. The exact copy number of *Pla2gc* in beaver genome need further exploration.

**Figure S12. Related to Figure 5.**
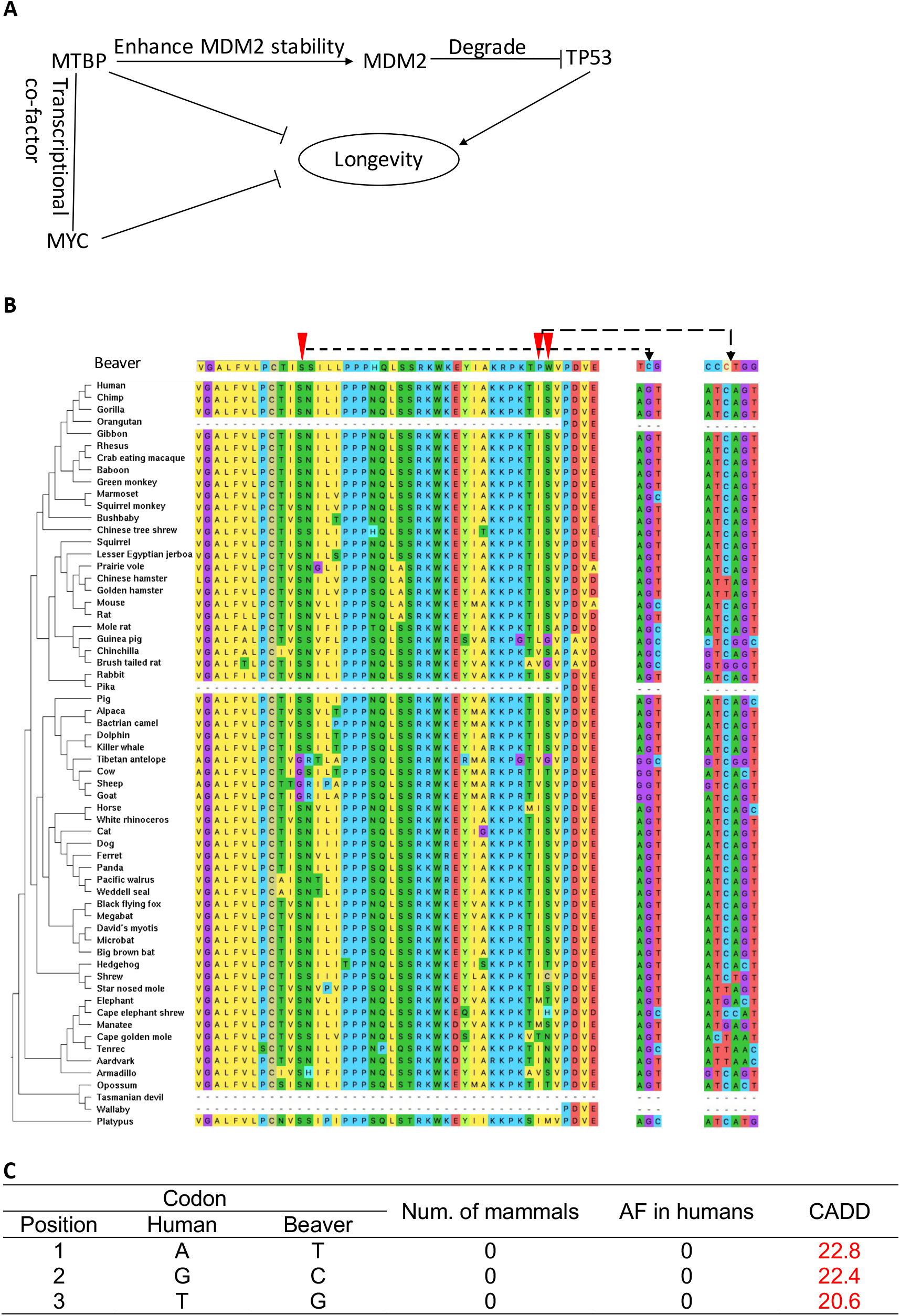
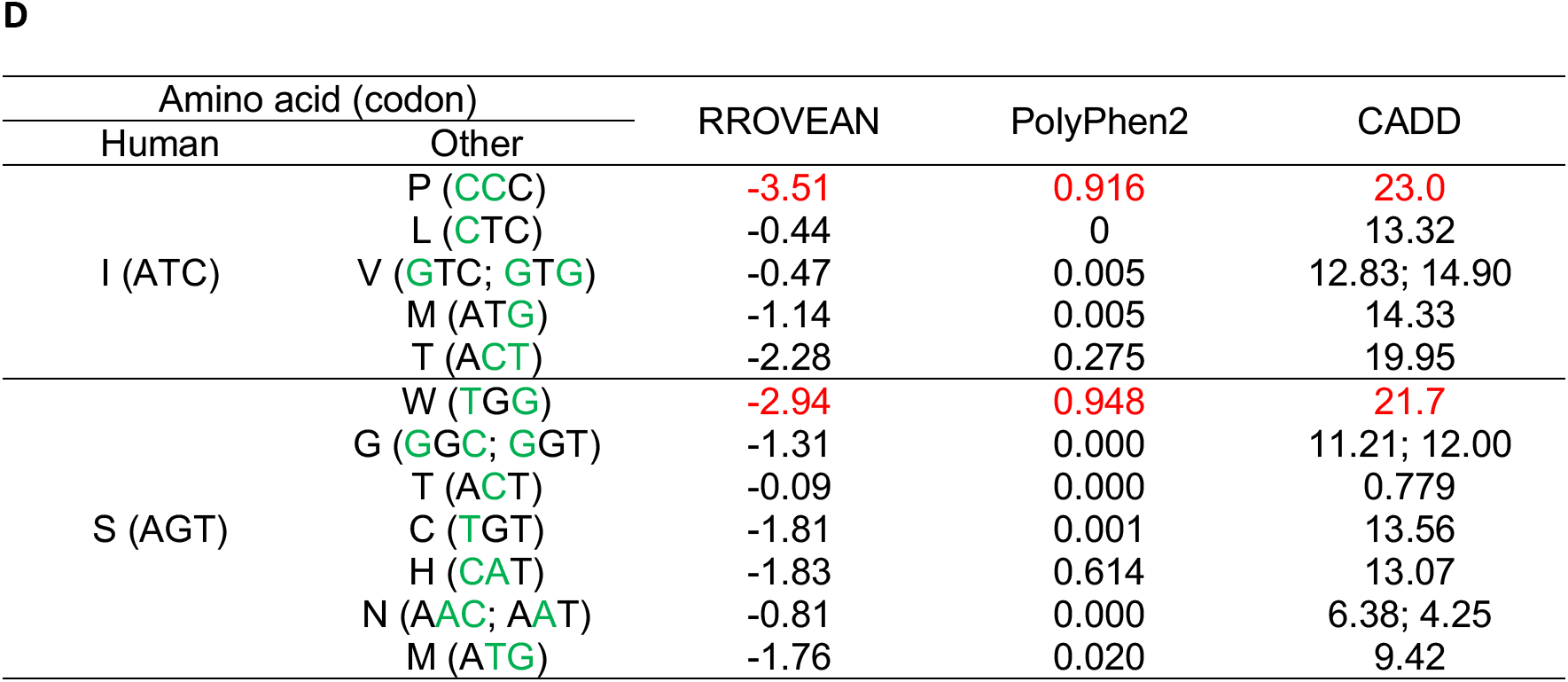
MTBP function and positive selection. (**A**) Functional interaction among MTBP and other genes related to longevity or cancer. MTBP enhances MDM2 stability by inhibiting auto-ubiquitination of MDM2, which in turn promotes TP53 degradation through MDM2-mediated ubiquitination of TP53. MTBP and MYC are transcriptional co-factors. Decreased expression of either MTBP and MYC show increased longevity and enhanced health-span(Hofmann, Zhao et al. 2015, Grieb, Boyd et al. 2016). (**B**) Alignment of 62 mammalian species at the three sites likely under positive selection in beavers. These three sites have selection probabilities higher than 88%. The alignment were extracted from 100-way vertebrates’ alignments. (**C**) One codon (S:TCG) under positive selection in beavers. It is a synonymous change (S/S: TCG/AGT) between human and beaver. ‘Num. of mammals’ is the number of mammalian species (among the 62 in the phylogeny) with the same nucleotide as beavers. ‘AF in humans’ is the frequency of the allele in humans identical to the nucleotide in beavers from gnomAD(Karczewski, Francioli et al. 2019). The CADD score quantifies the deleteriousness of the sequence change between human and beaver at each nucleotide position in the codon. A CADD score greater than 20 indicates a very likely deleterious change. (**D**) Other two codons (P:CCC and W:TGG) under positive selection in beavers. They are both non-synonymous change (P/I: CCC/ATC and W/S: TGG/AGT) between human and beaver. We predicted the functional effects of the non-synonymous changes between human and other mammalian species including beaver. Green letters are nucleotide changes from the human codons. PROVEAN scores lower than −2.5 indicate deleterious changes, which Polyphen2 scores between 0.85 to 1.0 indicate damaging changes.

**Figure S13. Related to Figure 6.**
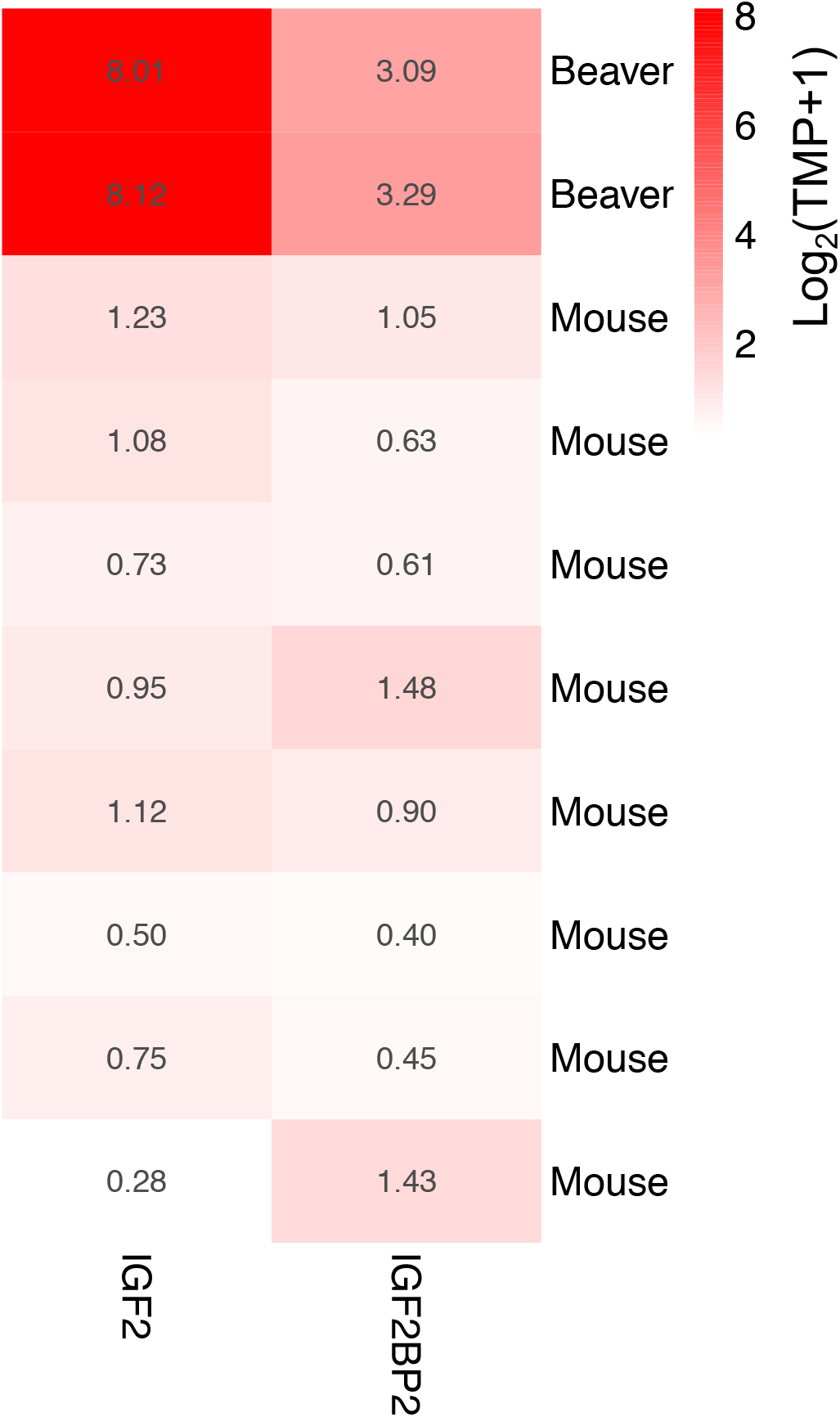
Comparison of *Igf2* and *Igf2bp2* expression between beavers and mice.

**Figure S14. Related to Figure 6.**
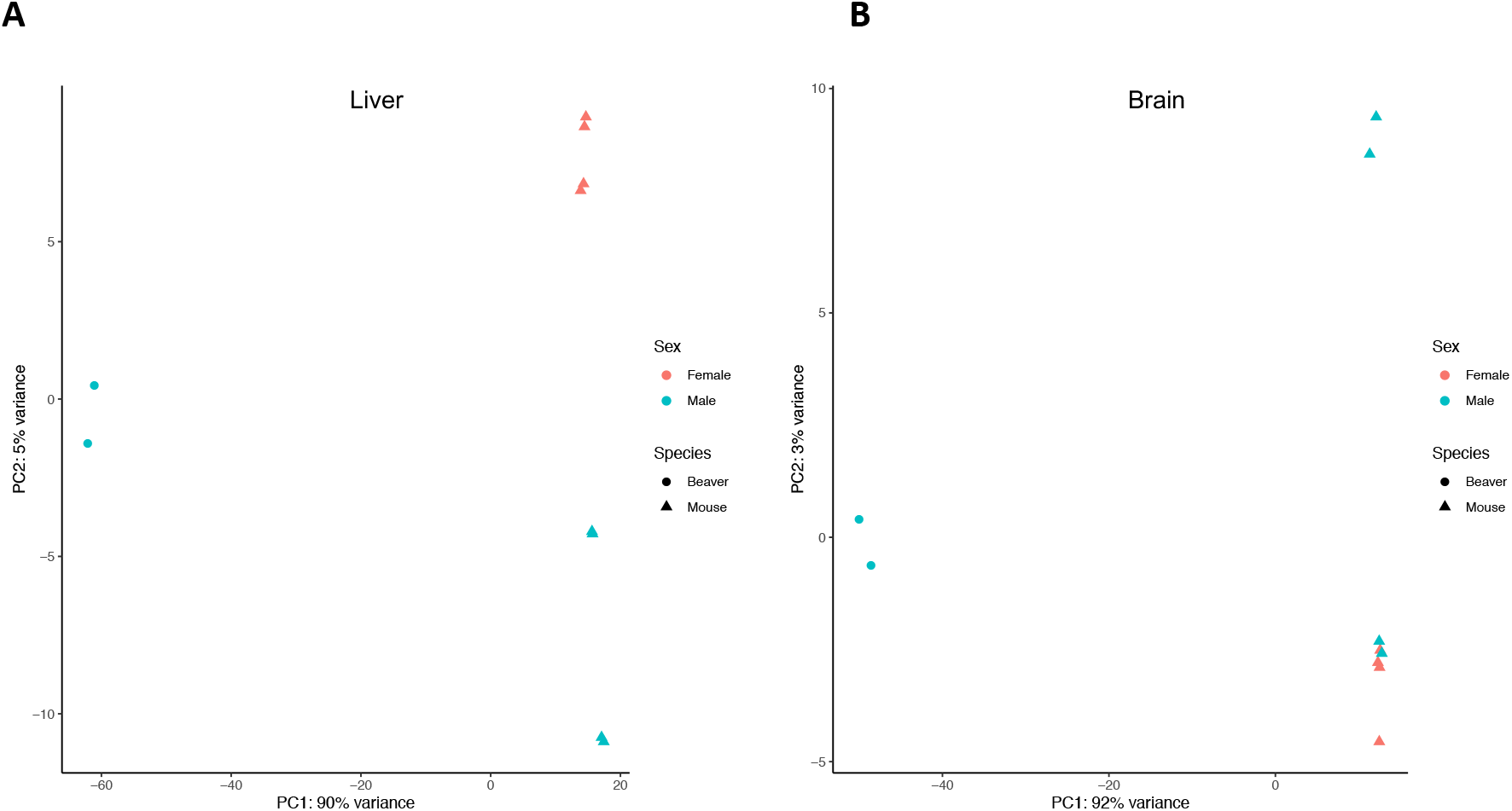
Principle component analysis of RNA-Seq data from beavers and mice. (**A**) Liver. (**B**) Brain.

**Table S1.**
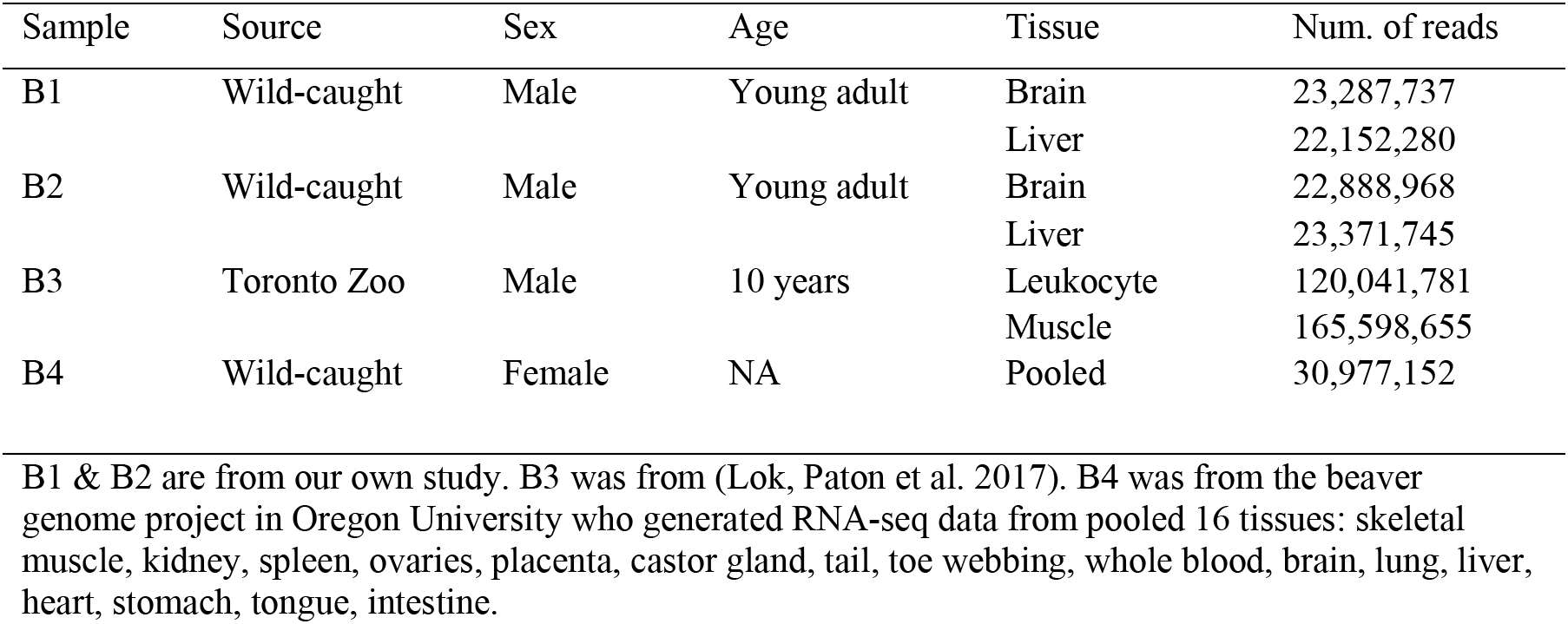
RNA-Seq data from four beaver samples.

| Sample | Source | Sex | Age | Tissue | Num. of reads |
| --- | --- | --- | --- | --- | --- |
| B1 | Wild-caught | Male | Young adult | Brain | 23,287,737 |
|  |  |  |  | Liver | 22,152,280 |
| B2 | Wild-caught | Male | Young adult | Brain | 22,888,968 |
|  |  |  |  | Liver | 23,371,745 |
| B3 | Toronto Zoo | Male | 10 years | Leukocyte | 120,041,781 |
|  |  |  |  | Muscle | 165,598,655 |
| B4 | Wild-caught | Female | NA | Pooled | 30,977,152 |

B1 & B2 are from our own study. B3 was from (Lok, Paton et al. 2017). B4 was from the beaver genome project in Oregon University who generated RNA-seq data from pooled 16 tissues: skeletal muscle, kidney, spleen, ovaries, placenta, castor gland, tail, toe webbing, whole blood, brain, lung, liver, heart, stomach, tongue, intestine.

**Table S2.**
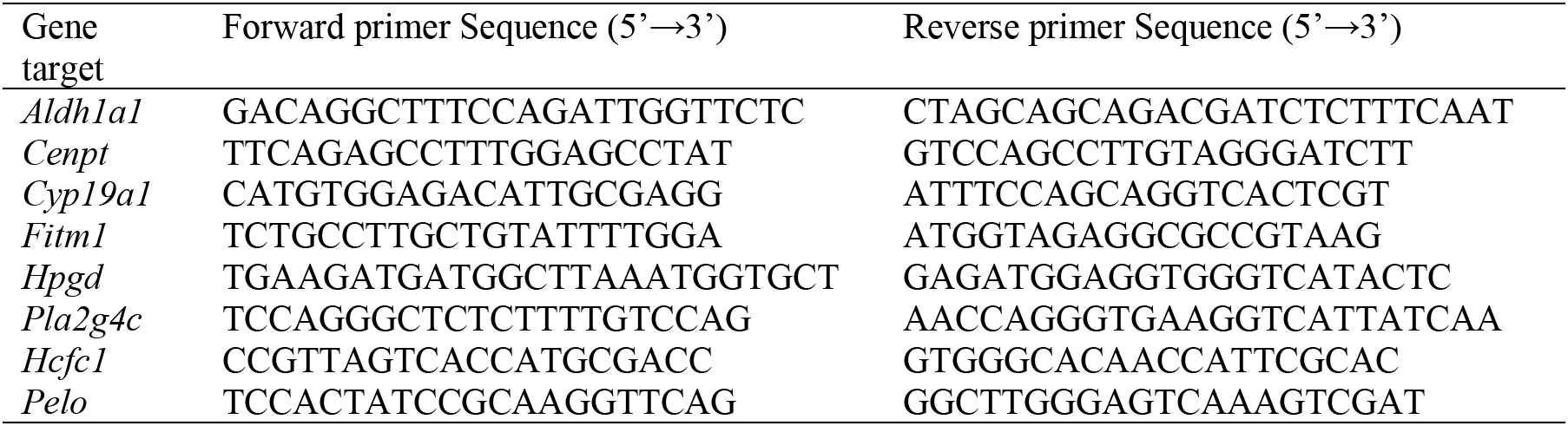
qPCR primers.

## References

Abegglen, L. M., A. F. Caulin, A. Chan, K. Lee, R. Robinson, M. S. Campbell, W. K. Kiso, D. L. Schmitt, P. J. Waddell, S. Bhaskara, S. T. Jensen, C. C. Maley and J. D. Schiffman (2015). “Potential Mechanisms for Cancer Resistance in Elephants and Comparative Cellular Response to DNA Damage in Humans.” JAMA 314(17): 1850–1860.

Adam, S. A., O. Schnell, J. Poschl, S. Eigenbrod, H. A. Kretzschmar, J. C. Tonn and U. Schuller (2012). “ALDH1A1 is a marker of astrocytic differentiation during brain development and correlates with better survival in glioblastoma patients.” Brain Pathol 22(6): 788–797.

Adzhubei, I., D. M. Jordan and S. R. Sunyaev (2013). “Predicting functional effect of human missense mutations using PolyPhen-2.” Curr Protoc Hum Genet Chapter 7: Unit7 20.

Ashur-Fabian, O., A. Avivi, L. Trakhtenbrot, K. Adamsky, M. Cohen, G. Kajakaro, A. Joel, N. Amariglio, E. Nevo and G. Rechavi (2004). “Evolution of p53 in hypoxia-stressed Spalax mimics human tumor mutation.” Proc Natl Acad Sci U S A 101(33): 12236–12241.

Ayala, A., M. F. Munoz and S. Arguelles (2014). “Lipid peroxidation: production, metabolism, and signaling mechanisms of malondialdehyde and 4-hydroxy-2-nonenal.” Oxid Med Cell Longev 2014: 360438.

Bao, W., K. K. Kojima and O. Kohany (2015). “Repbase Update, a database of repetitive elements in eukaryotic genomes.” Mob DNA 6: 11.

Bates, D., M. Mächler, B. Bolker and S. Walker (2015). “Fitting Linear Mixed-Effects Models Using lme4.” 2015 67(1): 48.

Belenguer-Varea, A., F. J. Tarazona-Santabalbina, J. A. Avellana-Zaragoza, M. Martinez-Reig, C. Mas-Bargues and M. Ingles (2019). “Oxidative stress and exceptional human longevity: Systematic review.” Free Radic Biol Med.

Birney, E., M. Clamp and R. Durbin (2004). “GeneWise and Genomewise.” Genome Res 14(5): 988–995.

Bray, N. L., H. Pimentel, P. Melsted and L. Pachter (2016). “Near-optimal probabilistic RNA-seq quantification.” Nat Biotechnol 34(5): 525–527.

Campbell, M. S., C. Holt, B. Moore and M. Yandell (2014). “Genome Annotation and Curation Using MAKER and MAKER-P.” Curr Protoc Bioinformatics 48: 4 11 11–39.

Capella-Gutierrez, S., J. M. Silla-Martinez and T. Gabaldon (2009). “trimAl: a tool for automated alignment trimming in large-scale phylogenetic analyses.” Bioinformatics 25(15): 1972–1973.

Carithers, L. J., K. Ardlie, M. Barcus, P. A. Branton, A. Britton, S. A. Buia, C. C. Compton, D. S. DeLuca, J. Peter-Demchok, E. T. Gelfand, P. Guan, G. E. Korzeniewski, N. C. Lockhart, C. A. Rabiner, A. K. Rao, K. L. Robinson, N. V. Roche, S. J. Sawyer, A. V. Segre, C. E. Shive, A. M. Smith, L. H. Sobin, A. H. Undale, K. M. Valentino, J. Vaught, T. R. Young, H. M. Moore and G. T. Consortium (2015). “A Novel Approach to High-Quality Postmortem Tissue Procurement: The GTEx Project.” Biopreserv Biobank 13(5): 311–319.

Carpenter, J. and C. J. Marion (2013). Exotic Animal Formulary. St. Louis, Missouri, USA, Elsevier Saunders.

Choi, Y. and A. P. Chan (2015). “PROVEAN web server: a tool to predict the functional effect of amino acid substitutions and indels.” Bioinformatics 31(16): 2745–2747.

Corbo, R. M., L. Ulizzi, L. Positano and R. Scacchi (2011). “Association of CYP19 and ESR1 pleiotropic genes with human longevity.” J Gerontol A Biol Sci Med Sci 66(1): 51–55.

Deepashree, S., S. Niveditha, T. Shivanandappa and S. R. Ramesh (2019). “Oxidative stress resistance as a factor in aging: evidence from an extended longevity phenotype of Drosophila melanogaster.” Biogerontology 20(4): 497–513.

Ding, Y., M. Tong, S. Liu, J. A. Moscow and H. H. Tai (2005). “NAD+-linked 15-hydroxyprostaglandin dehydrogenase (15-PGDH) behaves as a tumor suppressor in lung cancer.” Carcinogenesis 26(1): 65–72.

Domaradzki, P., M. Florek, P. Skalecki, A. Litwhiczuk, M. Kedzierska-Matysek, A. Wolanciuk and K. Tajchman (2019). “Fatty acid composition, cholesterol content and lipid oxidation indices of intramuscular fat from skeletal muscles of beaver (Castor fiber L.).” Meat Science 150: 131–140.

Doni, A., M. Stravalaci, A. Inforzato, E. Magrini, A. Mantovani, C. Garlanda and B. Bottazzi (2019). “The Long Pentraxin PTX3 as a Link Between Innate Immunity, Tissue Remodeling, and Cancer.” Front Immunol 10: 712.

Edgar, R. C. (2004). “MUSCLE: multiple sequence alignment with high accuracy and high throughput.” Nucleic Acids Res 32(5): 1792–1797.

Fang, X., I. Seim, Z. Huang, M. V. Gerashchenko, Z. Xiong, A. A. Turanov, Y. Zhu, A. V. Lobanov, D. Fan, S. H. Yim, X. Yao, S. Ma, L. Yang, S. G. Lee, E. B. Kim, R. T. Bronson, R. Sumbera, R. Buffenstein, X. Zhou, A. Krogh, T. J. Park, G. Zhang, J. Wang and V. N. Gladyshev (2014). “Adaptations to a subterranean environment and longevity revealed by the analysis of mole rat genomes.” Cell Rep 8(5): 1354–1364.

Ferreira, P. G., M. Munoz-Aguirre, F. Reverter, C. P. Sa Godinho, A. Sousa, A. Amadoz, R. Sodaei, M. R. Hidalgo, D. Pervouchine, J. Carbonell-Caballero, R. Nurtdinov, A. Breschi, R. Amador, P. Oliveira, C. Cubuk, J. Curado, F. Aguet, C. Oliveira, J. Dopazo, M. Sammeth, K. G. Ardlie and R. Guigo (2018). “The effects of death and post-mortem cold ischemia on human tissue transcriptomes.” Nat Commun 9(1): 490.

Galter, D., S. Buervenich, A. Carmine, M. Anvret and L. Olson (2003). “ALDH1 mRNA: presence in human dopamine neurons and decreases in substantia nigra in Parkinson’s disease and in the ventral tegmental area in schizophrenia.” Neurobiol Dis 14(3): 637–647.

Garaycoechea, J. I., G. P. Crossan, F. Langevin, L. Mulderrig, S. Louzada, F. Yang, G. Guilbaud, N. Park, S. Roerink, S. Nik-Zainal, M. R. Stratton and K. J. Patel (2018). “Alcohol and endogenous aldehydes damage chromosomes and mutate stem cells.” Nature 553(7687): 171–177.

Gasparetto, M., S. Sekulovic, C. Brocker, P. Tang, A. Zakaryan, P. Xiang, F. Kuchenbauer, M. Wen, K. Kasaian, M. F. Witty, P. Rosten, Y. Chen, S. Imren, G. Duester, D. C. Thompson, R. K. Humphries, V. Vasiliou and C. Smith (2012). “Aldehyde dehydrogenases are regulators of hematopoietic stem cell numbers and B-cell development.” Exp Hematol 40(4): 318–329 e312.

Gerbens, F., A. J. van Erp, F. L. Harders, F. J. Verburg, T. H. Meuwissen, J. H. Veerkamp and M. F. te Pas (1999). “Effect of genetic variants of the heart fatty acid-binding protein gene on intramuscular fat and performance traits in pigs.” J Anim Sci 77(4): 846–852.

Grieb, B. C., K. Boyd, R. Mitra and C. M. Eischen (2016). “Haploinsufficiency of the Myc regulator Mtbp extends survival and delays tumor development in aging mice.” Aging (Albany NY) 8(10): 2590–2602.

Haas, B. J., A. Papanicolaou, M. Yassour, M. Grabherr, P. D. Blood, J. Bowden, M. B. Couger, D. Eccles, B. Li, M. Lieber, M. D. Macmanes, M. Ott, J. Orvis, N. Pochet, F. Strozzi, N. Weeks, R. Westerman, T. William, C. N. Dewey, R. Henschel, R. D. Leduc, N. Friedman and A. Regev (2013). “De novo transcript sequence reconstruction from RNA-seq using the Trinity platform for reference generation and analysis.” Nat Protoc 8(8): 1494–1512.

Han, M. V., G. W. C. Thomas, J. Lugo-Martinez and M. W. Hahn (2013). “Estimating Gene Gain and Loss Rates in the Presence of Error in Genome Assembly and Annotation Using CAFE 3.” Molecular Biology and Evolution 30(8): 1987–1997.

Hofmann, J. W., X. Zhao, M. De Cecco, A. L. Peterson, L. Pagliaroli, J. Manivannan, G. B. Hubbard, Y. Ikeno, Y. Zhang, B. Feng, X. Li, T. Serre, W. Qi, H. Van Remmen, R. A. Miller, K. G. Bath, R. de Cabo, H. Xu, N. Neretti and J. M. Sedivy (2015). “Reduced expression of MYC increases longevity and enhances healthspan.” Cell 160(3): 477–488.

Holt, C. and M. Yandell (2011). “MAKER2: an annotation pipeline and genome-database management tool for second-generation genome projects.” BMC Bioinformatics 12: 491.

Horstman, A. M., E. L. Dillon, R. J. Urban and M. Sheffield-Moore (2012). “The role of androgens and estrogens on healthy aging and longevity.” J Gerontol A Biol Sci Med Sci 67(11): 1140–1152.

Houtkooper, R. H., L. Mouchiroud, D. Ryu, N. Moullan, E. Katsyuba, G. Knott, R. W. Williams and J. Auwerx (2013). “Mitonuclear protein imbalance as a conserved longevity mechanism.” Nature 497(7450): 451–457.

Hulbert, A. J., M. A. Kelly and S. K. Abbott (2014). “Polyunsaturated fats, membrane lipids and animal longevity.” J Comp Physiol B 184(2): 149–166.

Jha, P., M. T. McDevitt, R. Gupta, P. M. Quiros, E. G. Williams, K. Gariani, M. B. Sleiman, L. Diserens, A. Jochem, A. Ulbrich, J. J. Coon, J. Auwerx and D. J. Pagliarini (2018). “Systems Analyses Reveal Physiological Roles and Genetic Regulators of Liver Lipid Species.” Cell Syst 6(6): 722–733 e726.

Jobson, R. W., B. Nabholz and N. Galtier (2010). “An evolutionary genome scan for longevity-related natural selection in mammals.” Mol Biol Evol 27(4): 840–847.

Johnson, A. A. and A. Stolzing (2019). “The role of lipid metabolism in aging, lifespan regulation, and age-related disease.” Aging Cell 18(6): e13048.

Jones, P., D. Binns, H. Y. Chang, M. Fraser, W. Li, C. McAnulla, H. McWilliam, J. Maslen, A. Mitchell, G. Nuka, S. Pesseat, A. F. Quinn, A. Sangrador-Vegas, M. Scheremetjew, S. Y. Yong, R. Lopez and S. Hunter (2014). “InterProScan 5: genome-scale protein function classification.” Bioinformatics 30(9): 1236–1240.

Karczewski, K. J., L. C. Francioli, G. Tiao, B. B. Cummings, J. Alföldi, Q. Wang, R. L. Collins, K. M. Laricchia, A. Ganna, D. P. Birnbaum, L. D. Gauthier, H. Brand, M. Solomonson, N. A. Watts, D. Rhodes, M. Singer-Berk, E. M. England, E. G. Seaby, J. A. Kosmicki, R. K. Walters, K. Tashman, Y. Farjoun, E. Banks, T. Poterba, A. Wang, C. Seed, N. Whiffin, J. X. Chong, K. E. Samocha, E. Pierce-Hoffman, Z. Zappala, A. H. O’Donnell-Luria, E. V. Minikel, B. Weisburd, M. Lek, J. S. Ware, C. Vittal, I. M. Armean, L. Bergelson, K. Cibulskis, K. M. Connolly, M. Covarrubias, S. Donnelly, S. Ferriera, S. Gabriel, J. Gentry, N. Gupta, T. Jeandet, D. Kaplan, C. Llanwarne, R. Munshi, S. Novod, N. Petrillo, D. Roazen, V. Ruano-Rubio, A. Saltzman, M. Schleicher, J. Soto, K. Tibbetts, C. Tolonen, G. Wade, M. E. Talkowski, B. M. Neale, M. J. Daly and D. G. MacArthur (2019). “Variation across 141,456 human exomes and genomes reveals the spectrum of loss-of-function intolerance across human protein-coding genes.” bioRxiv: 531210.

Keane, M., J. Semeiks, A. E. Webb, Y. I. Li, V. Quesada, T. Craig, L. B. Madsen, S. van Dam, D. Brawand, P. I. Marques, P. Michalak, L. Kang, J. Bhak, H. S. Yim, N. V. Grishin, N. H. Nielsen, M. P. Heide-Jorgensen, E. M. Oziolor, C. W. Matson, G. M. Church, G. W. Stuart, J. C. Patton, J. C. George, R. Suydam, K. Larsen, C. Lopez-Otin, M. J. O’Connell, J. W. Bickham, B. Thomsen and J. P. de Magalhaes (2015). “Insights into the evolution of longevity from the bowhead whale genome.” Cell Rep 10(1): 112–122.

Kim, E. B., X. Fang, A. A. Fushan, Z. Huang, A. V. Lobanov, L. Han, S. M. Marino, X. Sun, A. A. Turanov, P. Yang, S. H. Yim, X. Zhao, M. V. Kasaikina, N. Stoletzki, C. Peng, P. Polak, Z. Xiong, A. Kiezun, Y. Zhu, Y. Chen, G. V. Kryukov, Q. Zhang, L. Peshkin, L. Yang, R. T. Bronson, R. Buffenstein, B. Wang, C. Han, Q. Li, L. Chen, W. Zhao, S. R. Sunyaev, T. J. Park, G. Zhang, J. Wang and V. N. Gladyshev (2011). “Genome sequencing reveals insights into physiology and longevity of the naked mole rat.” Nature 479(7372): 223–227.

Korf, I. (2004). “Gene finding in novel genomes.” BMC Bioinformatics 5: 59.

Langevin, F., G. P. Crossan, I. V. Rosado, M. J. Arends and K. J. Patel (2011). “Fancd2 counteracts the toxic effects of naturally produced aldehydes in mice.” Nature 475(7354): 53–58.

Lee, E., G. A. Helt, J. T. Reese, M. C. Munoz-Torres, C. P. Childers, R. M. Buels, L. Stein, I. H. Holmes, G. Elsik and S. E. Lewis (2013). “Web Apollo: a web-based genomic annotation editing platform.” Genome Biol 14(8): R93.

Levi, B. P., O. H. Yilmaz, G. Duester and S. J. Morrison (2009). “Aldehyde dehydrogenase 1a1 is dispensable for stem cell function in the mouse hematopoietic and nervous systems.” Blood 113(8): 1670–1680.

Li, B., T. Qing, J. Zhu, Z. Wen, Y. Yu, R. Fukumura, Y. Zheng, Y. Gondo and L. Shi (2017). “A Comprehensive Mouse Transcriptomic BodyMap across 17 Tissues by RNA-seq.” Sci Rep 7(1): 4200.

Li, J., S. Condello, J. Thomes-Pepin, X. Ma, Y. Xia, T. D. Hurley, D. Matei and J. X. Cheng (2017). “Lipid Desaturation Is a Metabolic Marker and Therapeutic Target of Ovarian Cancer Stem Cells.” Cell Stem Cell 20(3): 303–314 e305.

Li, Y. and J. P. de Magalhaes (2013). “Accelerated protein evolution analysis reveals genes and pathways associated with the evolution of mammalian longevity.” Age (Dordr) 35(2): 301–314.

Liberzon, A., C. Birger, H. Thorvaldsdottir, M. Ghandi, J. P. Mesirov and P. Tamayo (2015). “The Molecular Signatures Database (MSigDB) hallmark gene set collection.” Cell Syst 1(6): 417–425.

Liguori, I., G. Russo, F. Curcio, G. Bulli, L. Aran, D. Della-Morte, G. Gargiulo, G. Testa, F. Cacciatore, D. Bonaduce and P. Abete (2018). “Oxidative stress, aging, and diseases.” Clin Interv Aging 13: 757–772.

Lok, S., T. A. Paton, Z. Wang, G. Kaur, S. Walker, R. K. Yuen, W. W. Sung, J. Whitney, J. A. Buchanan, B. Trost, N. Singh, B. Apresto, N. Chen, M. Coole, T. J. Dawson, K. Ho, Z. Hu, S. Pullenayegum, K. Samler, A. Shipstone, F. Tsoi, T. Wang, S. L. Pereira, P. Rostami, C. A. Ryan, A. H. Tong, K. Ng, Y. Sundaravadanam, J. T. Simpson, B. K. Lim, M. D. Engstrom, C. J. Dutton, K. C. Kerr, M. Franke, W. Rapley, R. F. Wintle and S. W. Scherer (2017). “De Novo Genome and Transcriptome Assembly of the Canadian Beaver (Castor canadensis).” G3 (Bethesda) 7(2): 755–773.

Love, M. I., W. Huber and S. Anders (2014). “Moderated estimation of fold change and dispersion for RNA-seq data with DESeq2.” Genome Biol 15(12): 550.

Lu, D., C. Han and T. Wu (2014). “15-PGDH inhibits hepatocellular carcinoma growth through 15-keto-PGE2/PPARgamma-mediated activation of p21WAF1/Cip1.” Oncogene 33(9): 1101–1112.

Ma, S. and V. N. Gladyshev (2017). “Molecular signatures of longevity: Insights from cross-species comparative studies.” Semin Cell Dev Biol.

MacRae, S. L., M. M. Croken, R. B. Calder, A. Aliper, B. Milholland, R. R. White, A. Zhavoronkov, V. N. Gladyshev, A. Seluanov, V. Gorbunova, Z. D. Zhang and J. Vijg (2015). “DNA repair in species with extreme lifespan differences.” Aging (Albany NY) 7(12): 1171–1184.

Makia, N. L., P. Bojang, K. C. Falkner, D. J. Conklin and R. A. Prough (2011). “Murine hepatic aldehyde dehydrogenase 1a1 is a major contributor to oxidation of aldehydes formed by lipid peroxidation.” Chem Biol Interact 191(1-3): 278–287.

Marchitti, S. A., C. Brocker, D. Stagos and V. Vasiliou (2008). “Non-P450 aldehyde oxidizing enzymes: the aldehyde dehydrogenase superfamily.” Expert Opin Drug Metab Toxicol 4(6): 697–720.

Marchitti, S. A., R. A. Deitrich and V. Vasiliou (2007). “Neurotoxicity and metabolism of the catecholamine-derived 3,4-dihydroxyphenylacetaldehyde and 3,4-dihydroxyphenylglycolaldehyde: the role of aldehyde dehydrogenase.” Pharmacol Rev 59(2): 125–150.

Martysiak-Zurowska, D., K. Zalewski and R. Kamieniarz (2009). “Unusual odd-chain and trans-octadecenoic fatty acids in tissues of feral European beaver (Castorfiber), Eurasian badger (Melesmeles) and raccoon dog (Nyctereutesprocyonoides).” Comp Biochem Physiol B Biochem Mol Biol 153(2): 145–148.

Mitchell, T. W., R. Buffenstein and A. J. Hulbert (2007). “Membrane phospholipid composition may contribute to exceptional longevity of the naked mole-rat (Heterocephalus glaber): a comparative study using shotgun lipidomics.” Exp Gerontol 42(11): 1053–1062.

Mukherjee, A., H. A. Kenny and E. Lengyel (2017). “Unsaturated Fatty Acids Maintain Cancer Cell Stemness.” Cell Stem Cell 20(3): 291–292.

Muller-Schwarze, D. (2003). “The Beaver: Natural History of a Wetlands Engineer. Ithaca, New York : Cornell University Press.”.

Myung, S. J., R. M. Rerko, M. Yan, P. Platzer, K. Guda, A. Dotson, E. Lawrence, A. J. Dannenberg, A. K. Lovgren, G. Luo, T. P. Pretlow, R. A. Newman, J. Willis, D. Dawson and S. D. Markowitz (2006). “15-Hydroxyprostaglandin dehydrogenase is an in vivo suppressor of colon tumorigenesis.” Proc Natl Acad Sci U S A 103(32): 12098–12102.

Nylander, J. A. A. (2004). “MrModeltest v2. Program distributed by the author.” Evolutionary Biology Centre, Uppsala University.

Park, J., T. Miyakawa, A. Shiokawa, H. Nakajima-Adachi, M. Tanokura and S. Hachimura (2014). “Attenuation of migration properties of CD4+ T cells from aged mice correlates with decrease in chemokine receptor expression, response to retinoic acid, and RALDH expression compared to young mice.” Biosci Biotechnol Biochem 78(6): 976–980.

Patel, R. K. and M. Jain (2012). “NGS QC Toolkit: a toolkit for quality control of next generation sequencing data.” PLoS One 7(2): e30619.

Pollard, A. K., T. L. Ingram, C. A. Ortori, F. Shephard, M. Brown, S. Liddell, D. A. Barrett and L. Chakrabarti (2019). “A comparison of the mitochondrial proteome and lipidome in the mouse and long-lived Pipistrelle bats.” Aging (Albany NY) 11(6): 1664–1685.

Putnam, N. H., B. L. O’Connell, J. C. Stites, B. J. Rice, M. Blanchette, R. Calef, C. J. Troll, A. Fields, P. D. Hartley, C. W. Sugnet, D. Haussler, D. S. Rokhsar and R. E. Green (2016). “Chromosome-scale shotgun assembly using an in vitro method for long-range linkage.” Genome Res 26(3): 342–350.

Qu, W., Y. Zhou, Y. Zhang, Y. Lu, X. Wang, D. Zhao, Y. Yang and C. Zhang (2012). “MFEprimer-2.0: a fast thermodynamics-based program for checking PCR primer specificity.” Nucleic Acids Res 40(Web Server issue): W205–208.

Rentzsch, P., D. Witten, G. M. Cooper, J. Shendure and M. Kircher (2019). “CADD: predicting the deleteriousness of variants throughout the human genome.” Nucleic Acids Res 47(D1): D886–D894.

Sahm, A., M. Bens, M. Platzer and K. Szafranski (2017). “PosiGene: automated and easy-to-use pipeline for genome-wide detection of positively selected genes.” Nucleic Acids Res.

Scarabino, D., R. Scacchi, A. Pinto and R. M. Corbo (2015). “Genetic Basis of the Relationship Between Reproduction and Longevity: A Study on Common Variants of Three Genes in Steroid Hormone Metabolism--CYP17, HSD17B1, and COMT.” Rejuvenation Res 18(5): 464–472.

Schmidleithner, L., Y. Thabet, E. Schonfeld, M. Kohne, D. Sommer, Z. Abdullah, T. Sadlon, C. Osei-Sarpong, K. Subbaramaiah, F. Copperi, K. Haendler, T. Varga, O. Schanz, S. Bourry, K. Bassler, W. Krebs, A. E. Peters, A. K. Baumgart, M. Schneeweiss, K. Klee, S. V. Schmidt, S. Nussing, J. Sander, N. Ohkura, A. Waha, T. Sparwasser, F. T. Wunderlich, I. Forster, T. Ulas, H. Weighardt, S. Sakaguchi, A. Pfeifer, M. Bluher, A. J. Dannenberg, N. Ferreiros, L. J. Muglia, C. Wickenhauser, S. C. Barry, J. L. Schultze and M. Beyer (2019). “Enzymatic Activity of HPGD in Treg Cells Suppresses Tconv Cells to Maintain Adipose Tissue Homeostasis and Prevent Metabolic Dysfunction.” Immunity 50(5): 1232–1248 e1214.

Seluanov, A., V. N. Gladyshev, J. Vijg and V. Gorbunova (2018). “Mechanisms of cancer resistance in long-lived mammals.” Nat Rev Cancer 18(7): 433–441.

Simao, F. A., R. M. Waterhouse, P. Ioannidis, E. V. Kriventseva and E. M. Zdobnov (2015). “BUSCO: assessing genome assembly and annotation completeness with single-copy orthologs.” Bioinformatics 31(19): 3210–3212.

Singh, S., C. Brocker, V. Koppaka, Y. Chen, B. C. Jackson, A. Matsumoto, D. C. Thompson and V. Vasiliou (2013). “Aldehyde dehydrogenases in cellular responses to oxidative/electrophilic stress.” Free Radic Biol Med 56: 89–101.

Soneson, C., M. I. Love and M. D. Robinson (2015). “Differential analyses for RNA-seq: transcript-level estimates improve gene-level inferences.” F1000Res 4: 1521.

Song, G. X., Y. H. Shen, Y. Q. Liu, W. Sun, L. P. Miao, L. J. Zhou, H. L. Liu, R. Yang, X. Q. Kong, K. J. Cao, L. M. Qian and Y. H. Sheng (2012). “Overexpression of FABP3 promotes apoptosis through inducing mitochondrial impairment in embryonic cancer cells.” J Cell Biochem 113(12): 3701–3708.

St-Onge, M. P. and D. Gallagher (2010). “Body composition changes with aging: the cause or the result of alterations in metabolic rate and macronutrient oxidation?” Nutrition 26(2): 152–155.

Stamatakis, A. (2014). “RAxML version 8: a tool for phylogenetic analysis and post-analysis of large phylogenies.” Bioinformatics 30(9): 1312–1313.

Stanke, M. and S. Waack (2003). “Gene prediction with a hidden Markov model and a new intron submodel.” Bioinformatics 19 Suppl 2: ii215–225.

Subramanian, A., P. Tamayo, V. K. Mootha, S. Mukherjee, B. L. Ebert, M. A. Gillette, A. Paulovich, S. L. Pomeroy, T. R. Golub, E. S. Lander and J. P. Mesirov (2005). “Gene set enrichment analysis: a knowledge-based approach for interpreting genome-wide expression profiles.” Proc Natl Acad Sci U S A 102(43): 15545–15550.

Tamura, K., F. U. Battistuzzi, P. Billing-Ross, O. Murillo, A. Filipski and S. Kumar (2012). “Estimating divergence times in large molecular phylogenies.” Proc Natl Acad Sci U S A 109(47): 19333–19338.

Tarailo-Graovac, M. and N. Chen (2009). “Using RepeatMasker to identify repetitive elements in genomic sequences.” Curr Protoc Bioinformatics Chapter 4: Unit 4 10.

The UniProt, C. (2017). “UniProt: the universal protein knowledgebase.” Nucleic Acids Res 45(D1): D158–D169.

Trumble, S. J. and S. B. Kanatous (2012). “Fatty Acid use in Diving Mammals: More than Merely Fuel.” Front Physiol 3: 184.

Vassalli, G. (2019). “Aldehyde Dehydrogenases: Not Just Markers, but Functional Regulators of Stem Cells.” Stem Cells Int 2019: 3904645.

Vergnes, L., R. Chin, S. G. Young and K. Reue (2011). “Heart-type fatty acid-binding protein is essential for efficient brown adipose tissue fatty acid oxidation and cold tolerance.” J Biol Chem 286(1): 380–390.

Wang, G., X. Zhang, J. S. Lee, X. Wang, Z. Q. Yang and K. Zhang (2012). “Endoplasmic reticulum factor ERLIN2 regulates cytosolic lipid content in cancer cells.” Biochem J 446(3): 415–425.

Wolfson, M., A. Budovsky, R. Tacutu and V. Fraifeld (2009). “The signaling hubs at the crossroad of longevity and age-related disease networks.” Int J Biochem Cell Biol 41(3): 516–520.

Wu, R., T. Liu, P. Yang, X. Liu, F. Liu, Y. Wang, H. Xiong, S. Yu, X. Huang and L. Zhuang (2017). “Association of 15-hydroxyprostaglandin dehydrogenate and poor prognosis of obese breast cancer patients.” Oncotarget 8(14): 22842–22853.

Ye, J., G. Coulouris, I. Zaretskaya, I. Cutcutache, S. Rozen and T. L. Madden (2012). “Primer-BLAST: a tool to design target-specific primers for polymerase chain reaction.” BMC Bioinformatics 13: 134.

Ye, M. H., J. L. Chen, G. P. Zhao, M. Q. Zheng and J. Wen (2010). “Associations of A-FABP and H-FABP markers with the content of intramuscular fat in Beijing-You chicken.” Anim Biotechnol 21(1): 14–24.

Zalewski, K., D. Martysiak-Zurowska, M. Chylinska-Ptak and B. Nitkiewicz (2009). “Characterization of Fatty Acid Composition in the European Beaver (Castor fiber L.).” Polish Journal of Environmental Studies 18(3): 493–499.

Zdobnov, E. M., F. Tegenfeldt, D. Kuznetsov, R. M. Waterhouse, F. A. Simao, P. Ioannidis, M. Seppey, A. Loetscher and E. V. Kriventseva (2017). “OrthoDB v9.1: cataloging evolutionary and functional annotations for animal, fungal, plant, archaeal, bacterial and viral orthologs.” Nucleic Acids Res 45(D1): D744–D749.

Zhang, J., R. Nielsen and Z. Yang (2005). “Evaluation of an improved branch-site likelihood method for detecting positive selection at the molecular level.” Mol Biol Evol 22(12): 2472–2479.

Zhou, X., Q. H. Dou, F. G.Y., Z. Q.W., M. Sanderford, A. Kaya, J. Johnson, E. Karlsson, X. Tian, M. A., S. Kuma, A. Seluanov, Z. D. Zhang, V. Gorbunova, L. X. and V. N. Gladyshev (2020). “Beaver and naked mole rat genomes reveal common approaches to longevity.” Cell Rep(Accepted).

Ziouzenkova, O., G. Orasanu, M. Sharlach, T. E. Akiyama, J. P. Berger, J. Viereck, J. A. Hamilton, G. Tang, G. G. Dolnikowski, S. Vogel, G. Duester and J. Plutzky (2007). “Retinaldehyde represses adipogenesis and diet-induced obesity.” Nat Med 13(6): 695–702.

