## Supplemental File 1 for "The genome of North American beaver provides insights into the mechanisms of its longevity and cancer resistance"

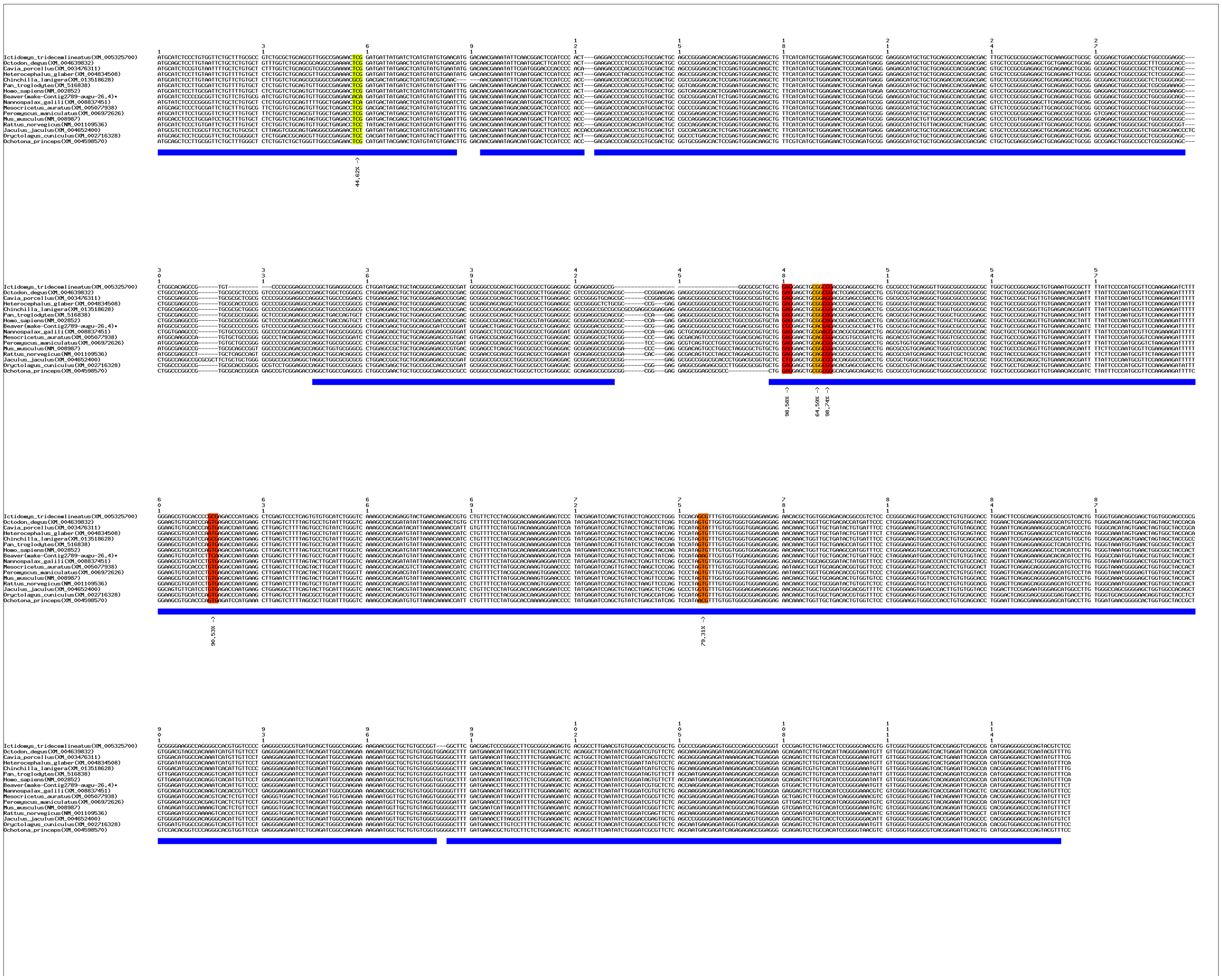

**(a)** Positive selection signals in *Ptx3* shown by multiple sequence alignment of coding sequences.

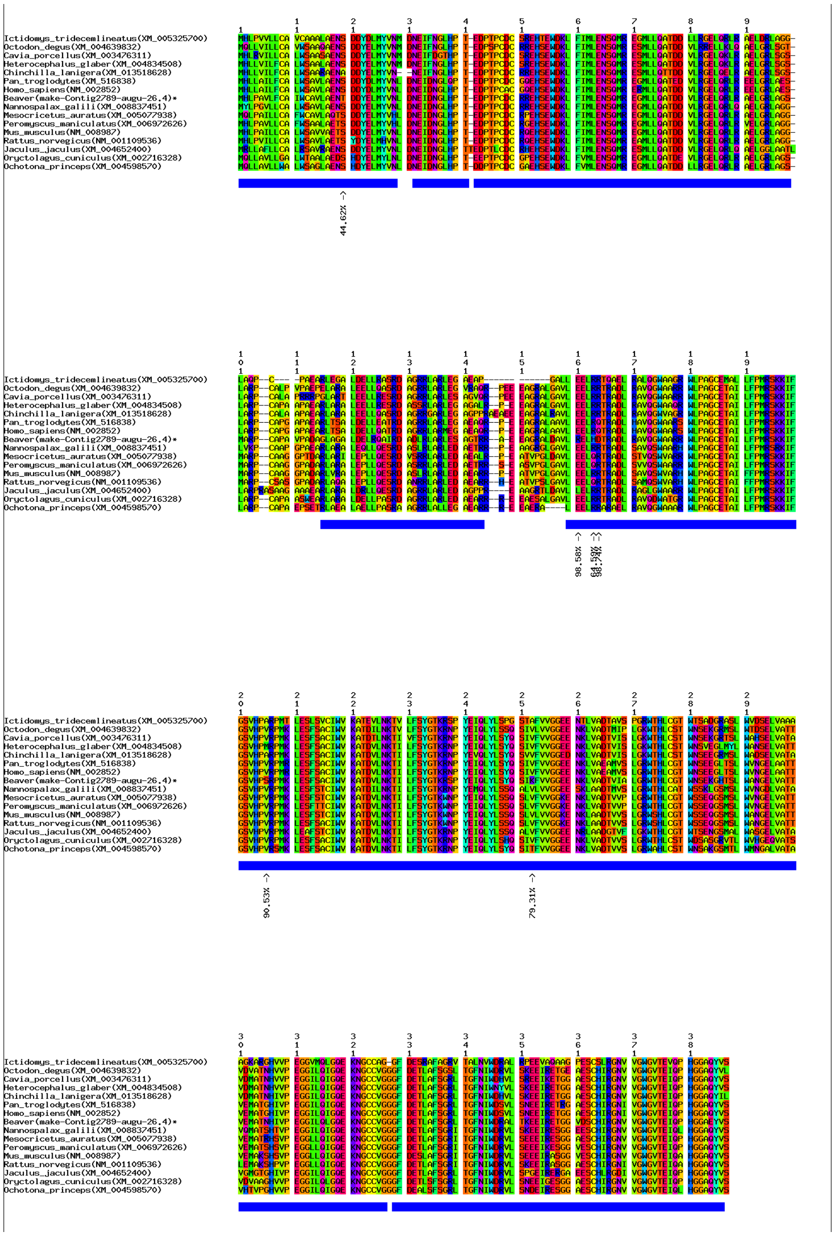

**(b)** Positive selection signals in *Ptx3* shown by multiple sequence alignment of protein sequences.

**Supplementary Figure 15. Positive selection signals in gene *Ptx3.***

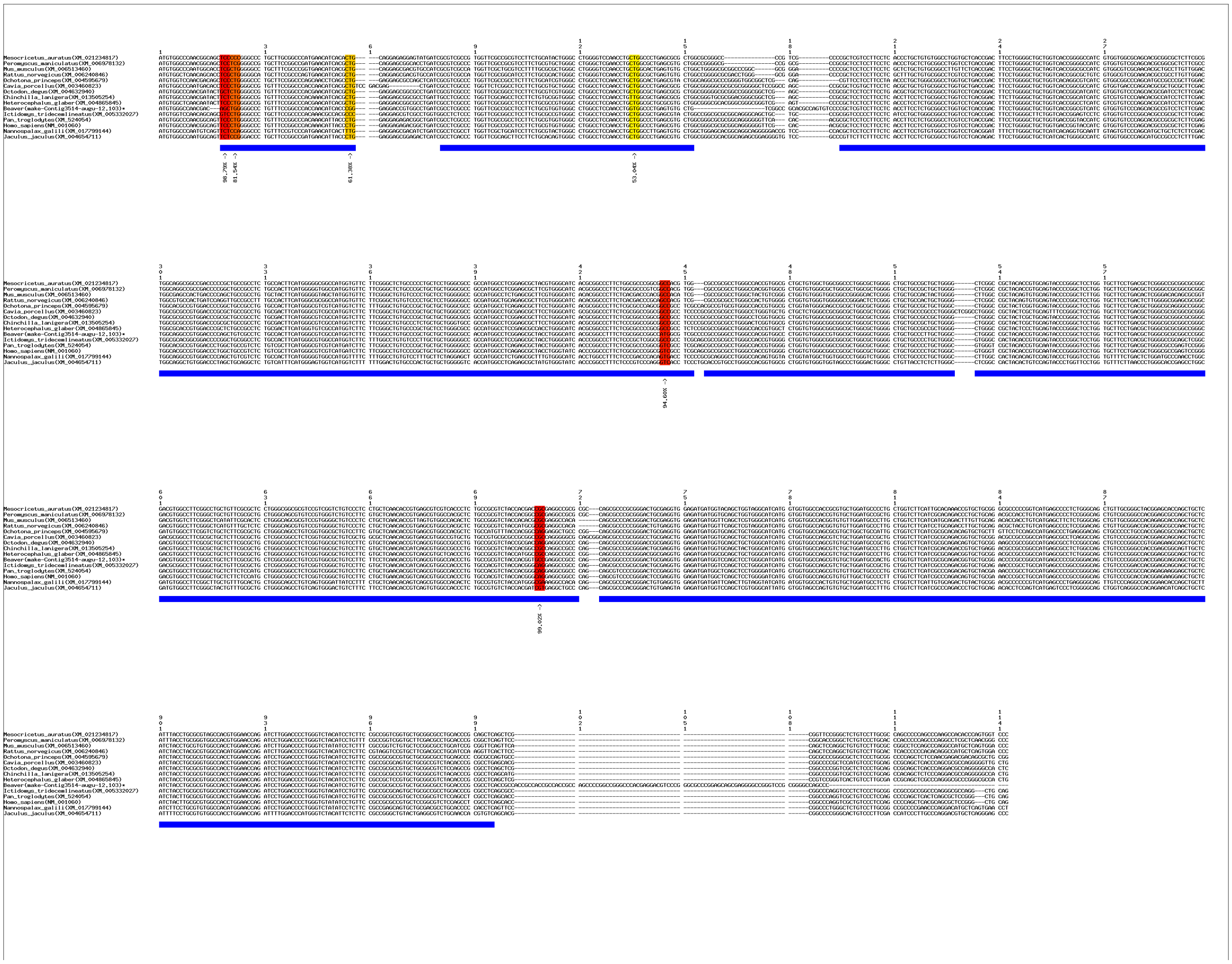

**(a)** Positive selection signals in *Tbxa2r* shown by multiple sequence alignment of coding sequences.

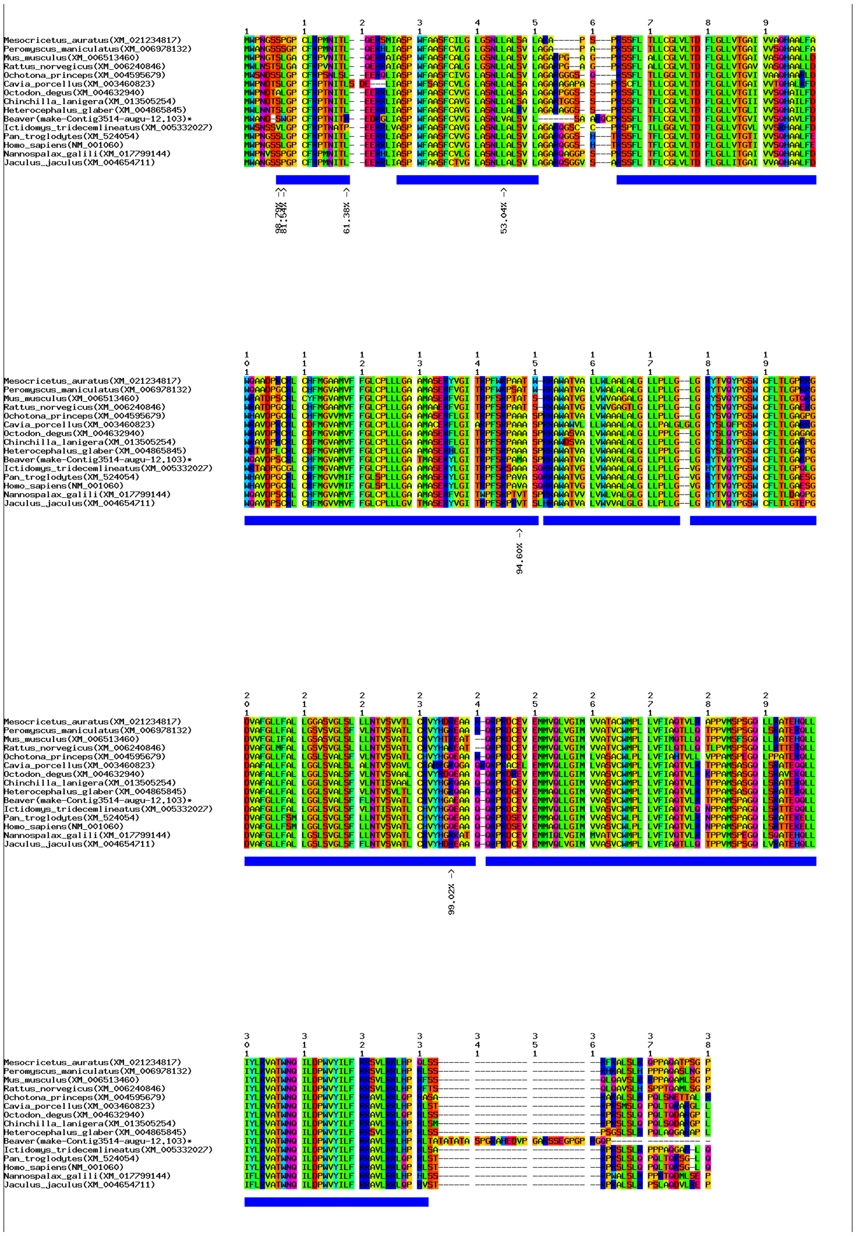

**(b)** Positive selection signals in *Tbxa2r* shown by multiple sequence alignment of protein sequences.
**Supplementary Figure 16. Positive selection signals in gene *Tbxa2r*.**

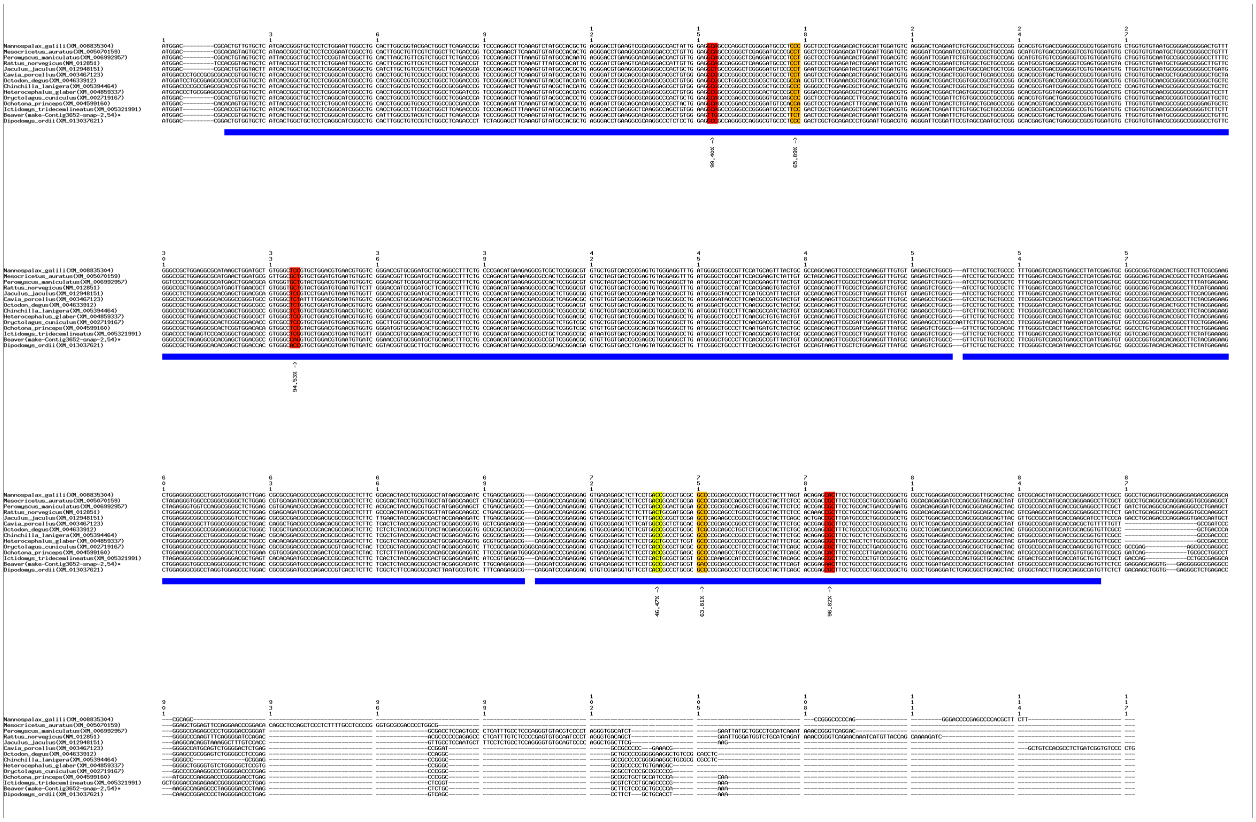

**(a)** Positive selection signals in *Hsd17b1* shown by multiple sequence alignment of coding sequences.

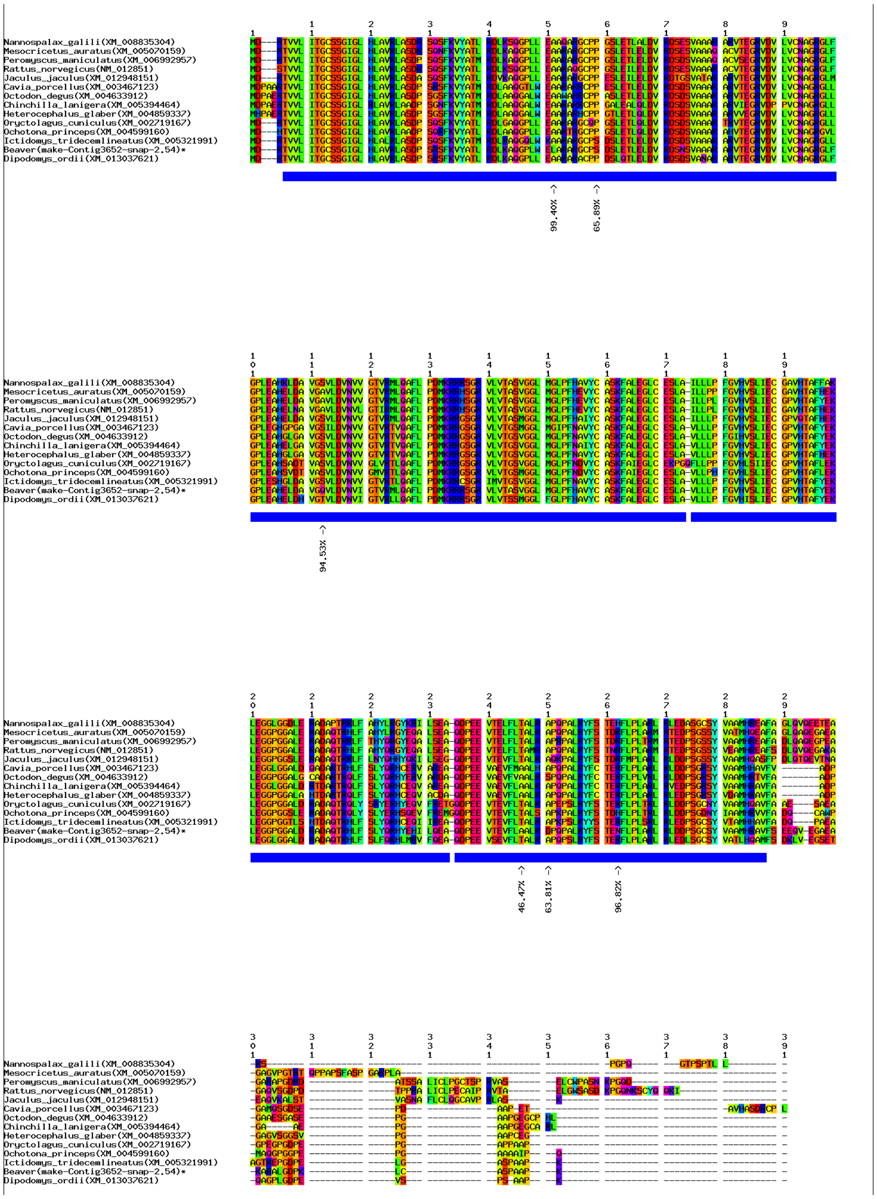

**(b)** Positive selection signals in *Hsd17b1* shown by multiple sequence alignment of protein sequences.

**Supplementary Figure 17. Positive selection signals in gene *Hsd17b1.***

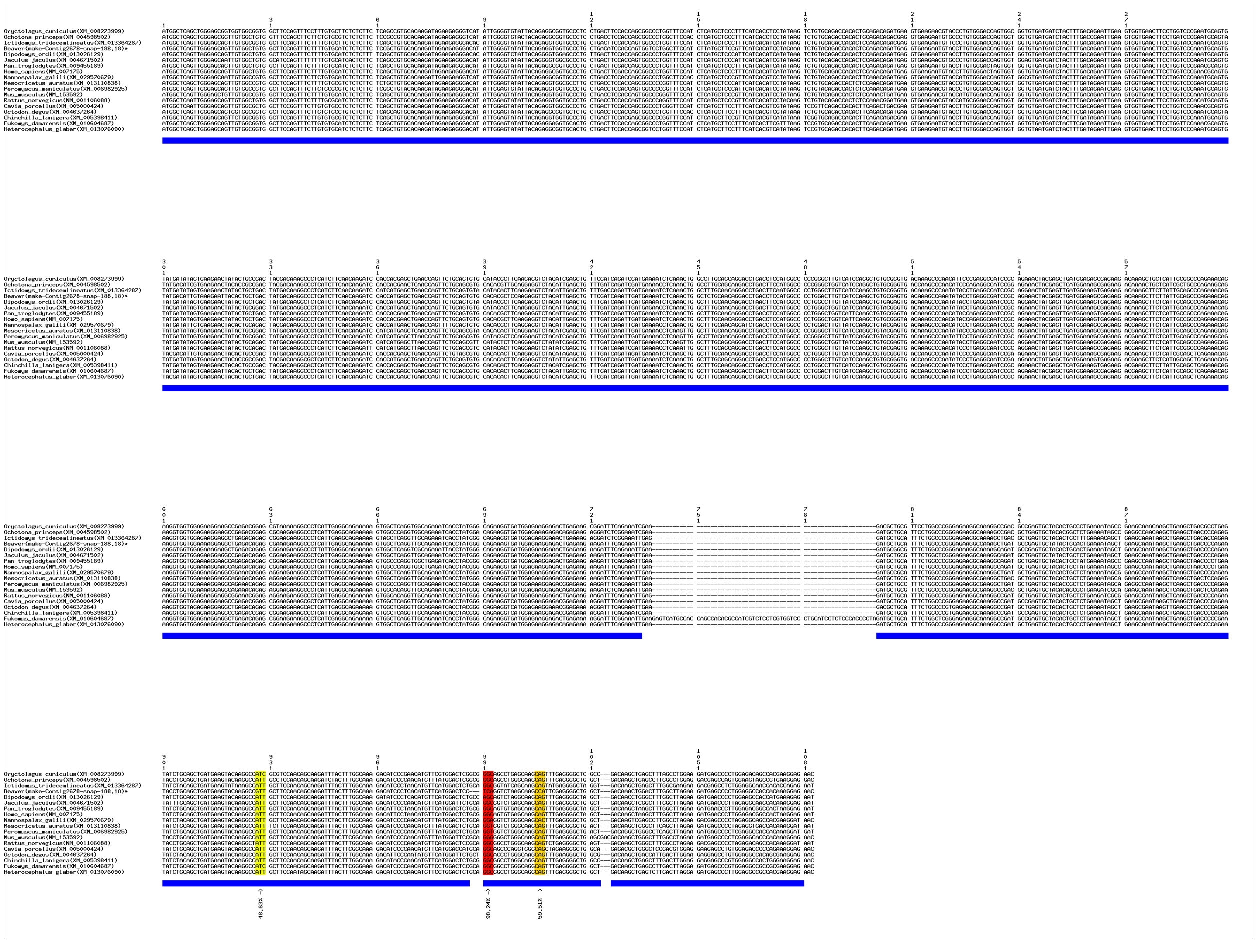

**(a)** Positive selection signals in *Erlin2* shown by multiple sequence alignment of coding sequences.

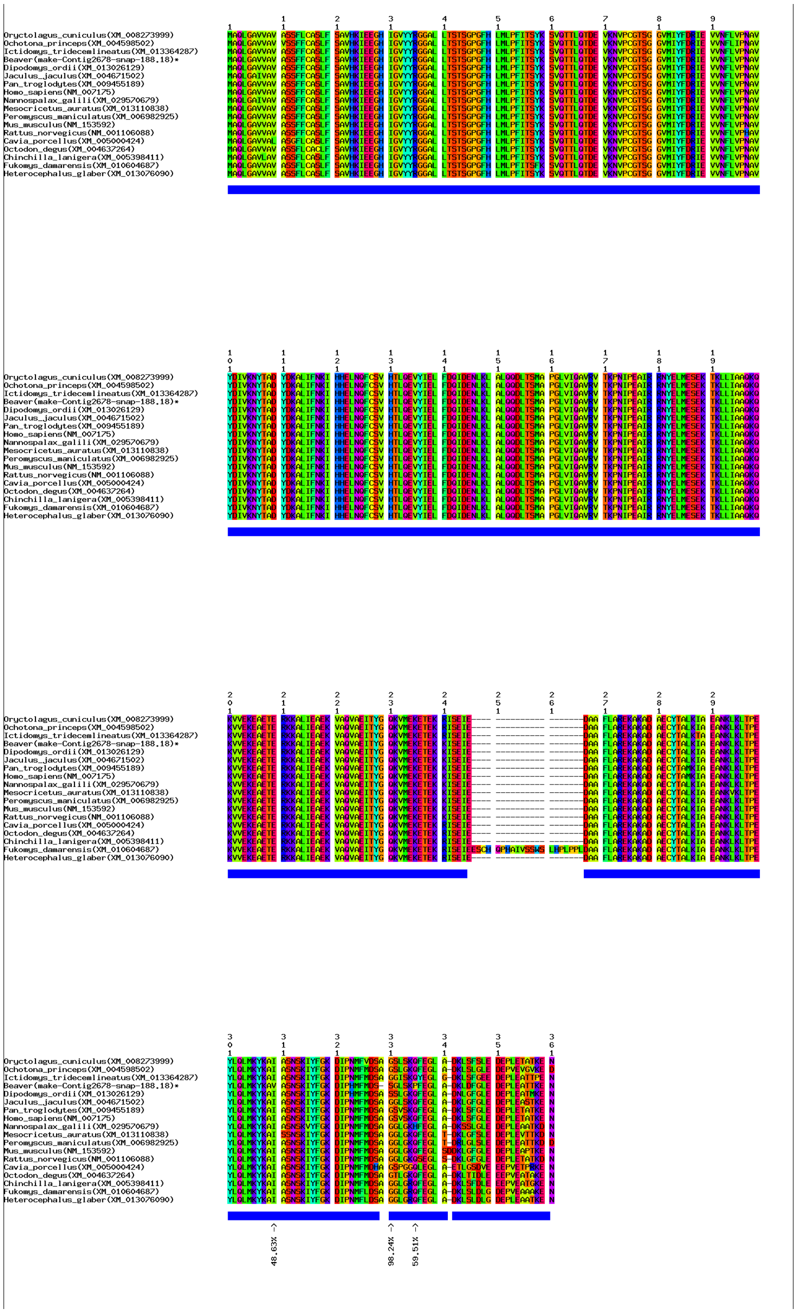

**(b)** Positive selection signals in *Erlin2* shown by multiple sequence alignment of protein sequences.
**Supplementary Figure 18. Positive selection signals in gene *Erlin2*.**

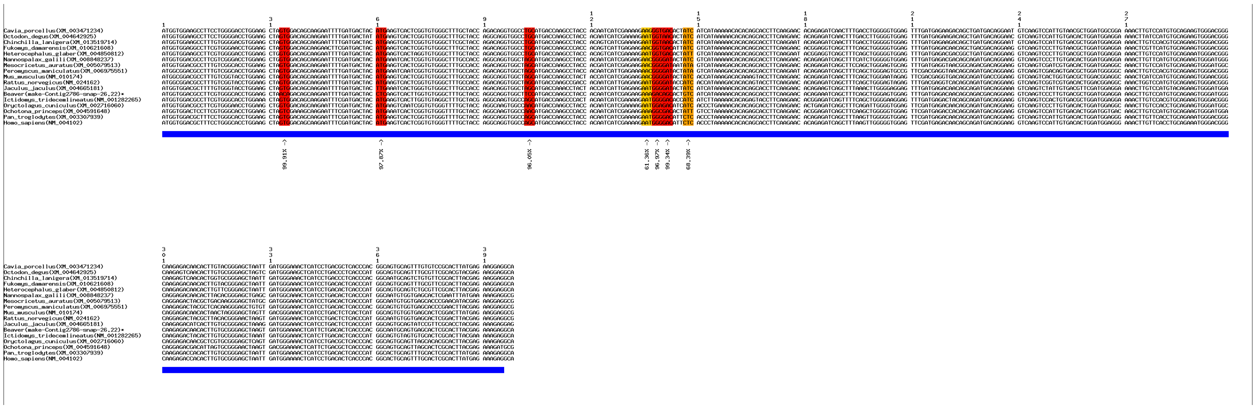

**(a)** Positive selection signals in *Fabp3* shown by multiple sequence alignment of coding sequences.

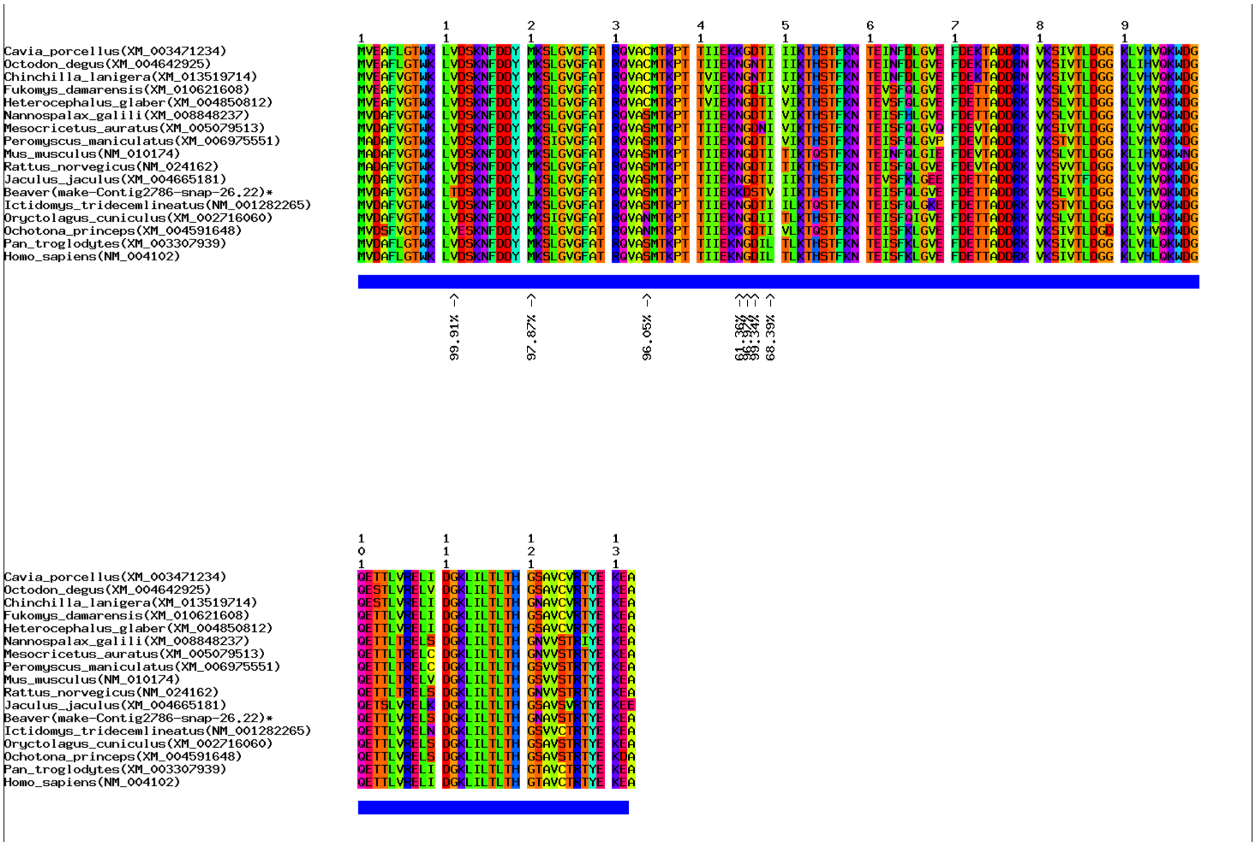


**(b)** Positive selection signals in *Fabp3* shown by multiple sequence alignment of protein sequences.

**Supplementary Figure 19. Positive selection signals in gene *Fabp3*.**

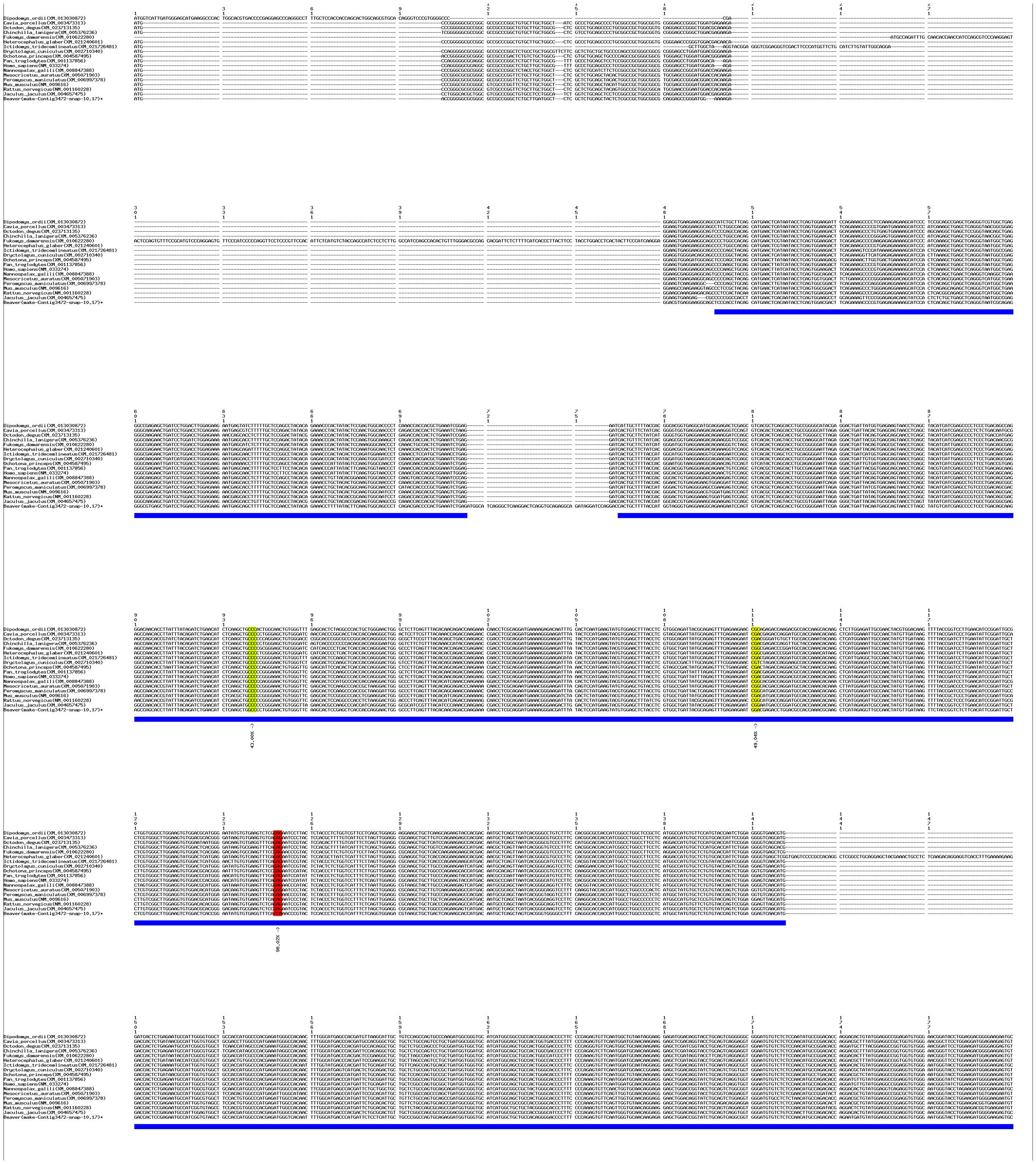

**(a)** Positive selection signals in *Adam19* shown by multiple sequence alignment of coding sequences.

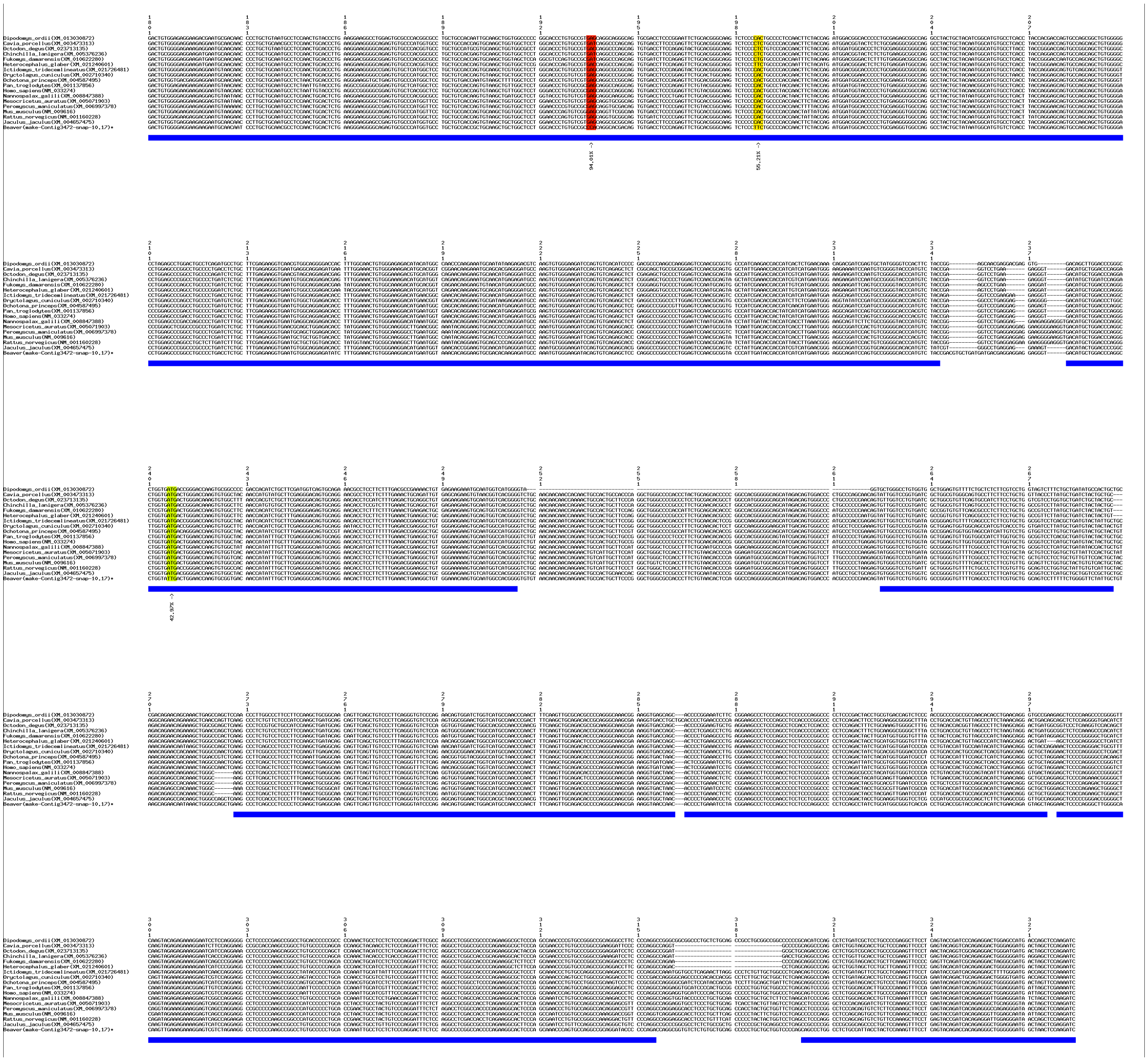

**(a)** Positive selection signals in *Adam19* shown by multiple sequence alignment of coding sequences (continue).

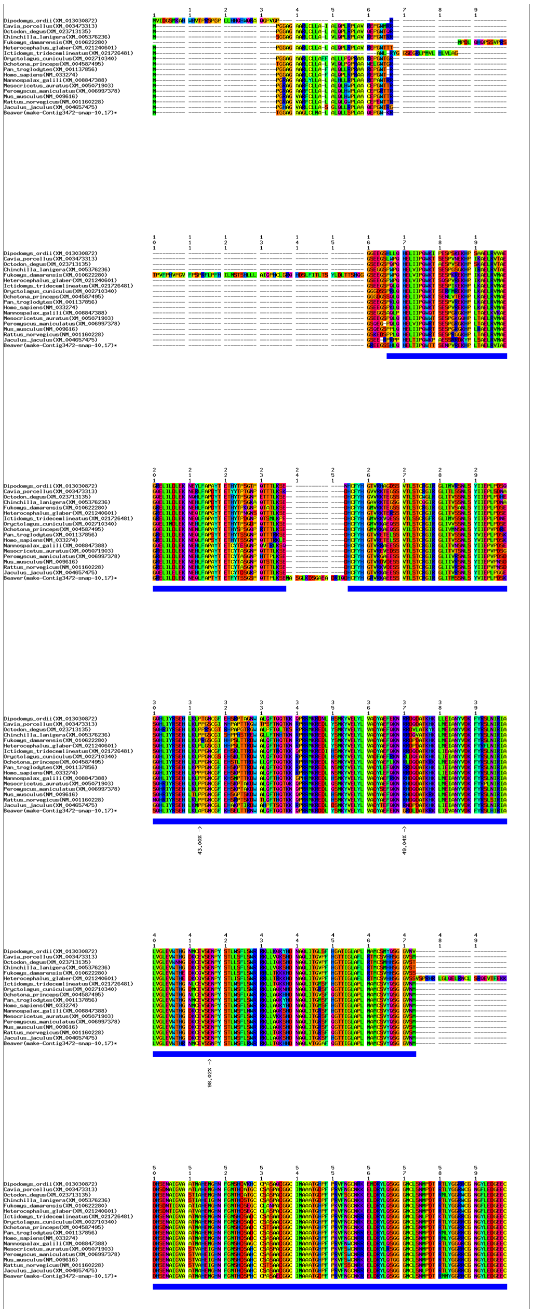

**(b)** Positive selection signals in *Adam19* shown by multiple sequence alignment of protein sequences.

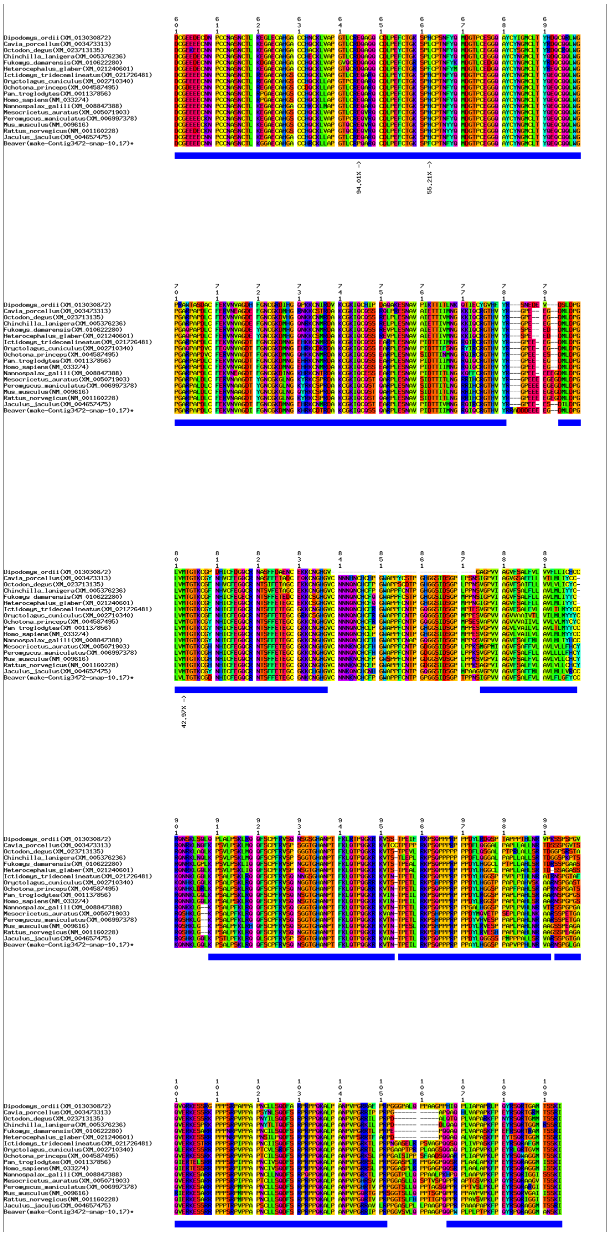

**(b)** Positive selection signals in *Adam19* shown by multiple sequence alignment of protein sequences (continue).

**Supplementary Figure 20. Positive selection signals in gene *Adam19.***

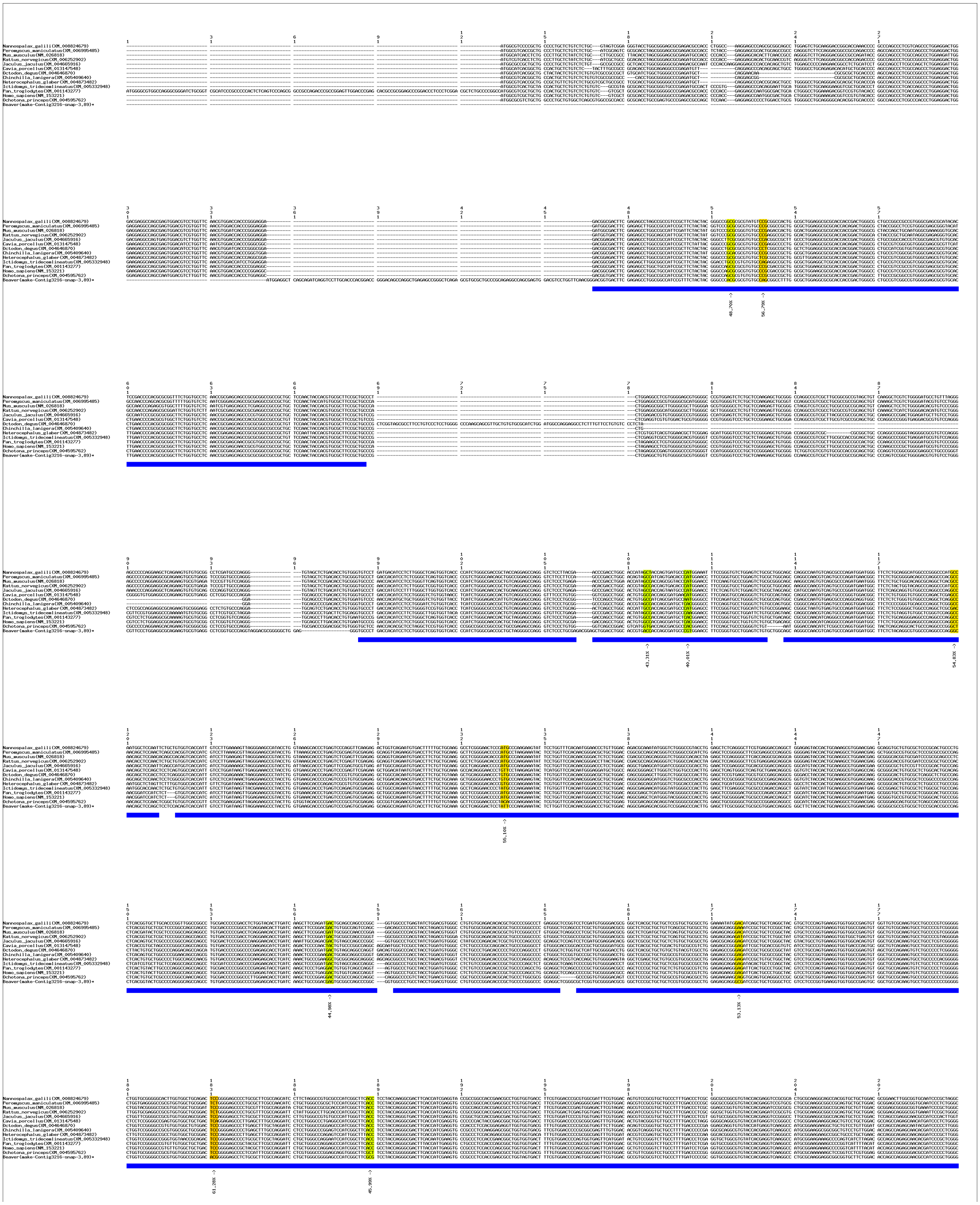

**(a)** Positive selection signals in *Clip2* shown by multiple sequence alignment of coding sequences.

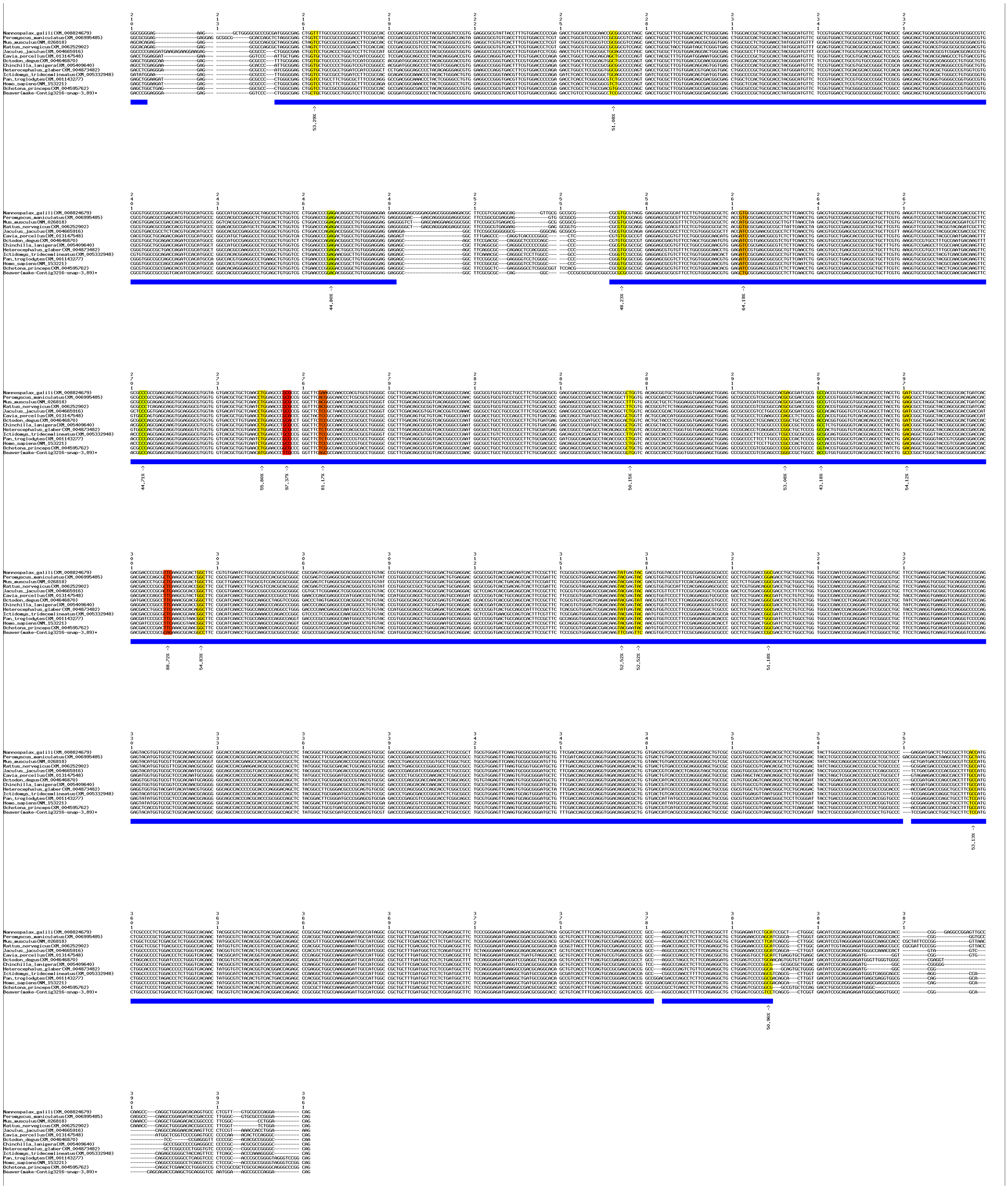

**(a)** Positive selection signals in *Clip2* shown by multiple sequence alignment of coding sequences (continue).

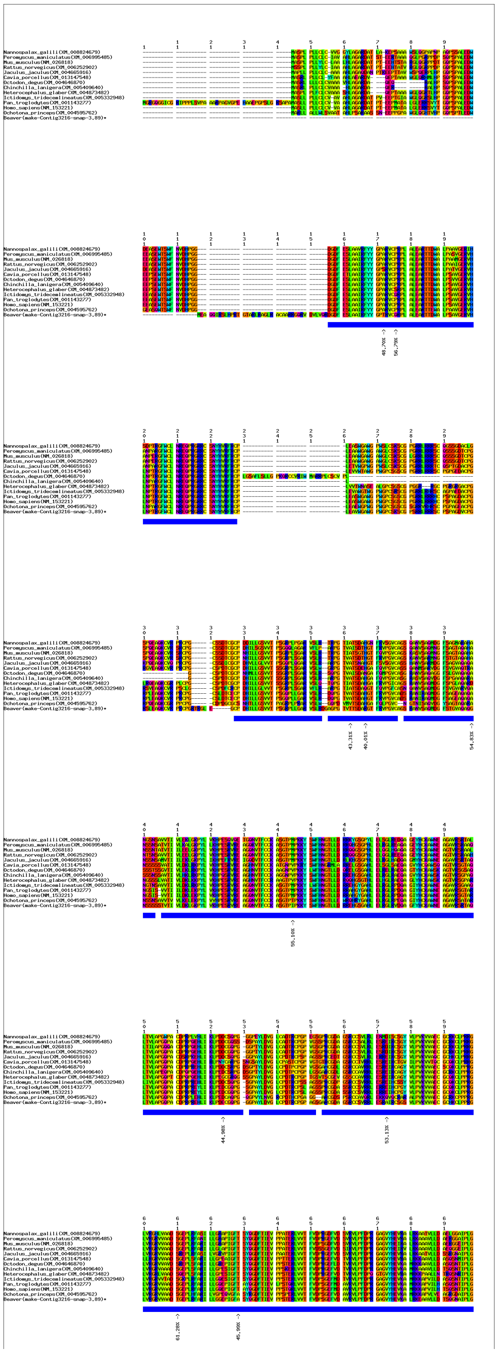

**(b)** Positive selection signals in *Clip2* shown by multiple sequence alignment of protein sequences.

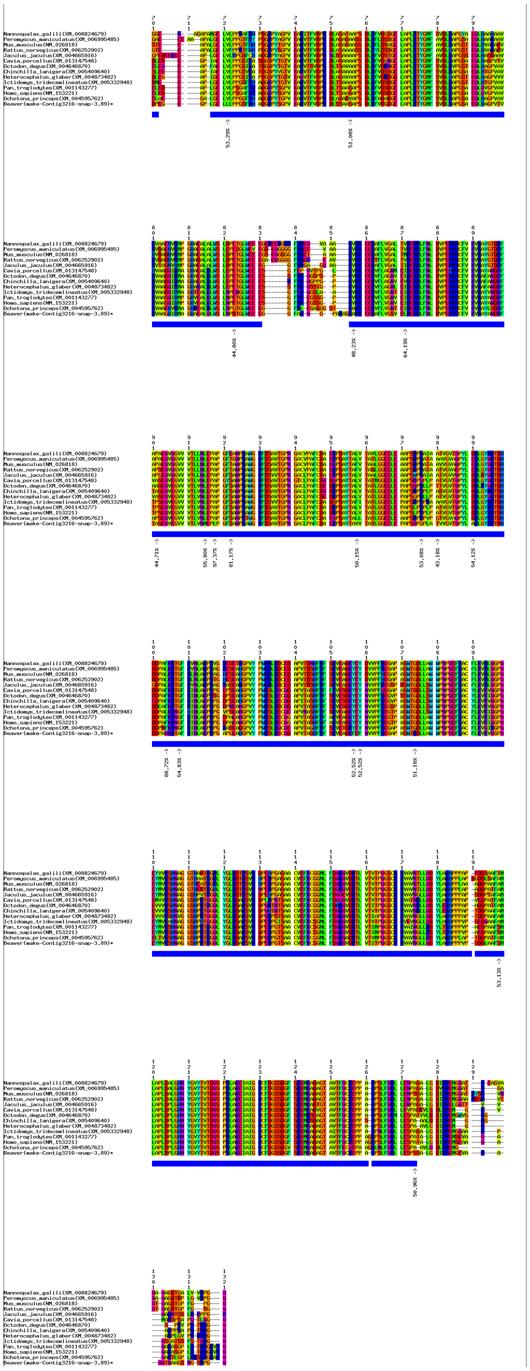

**(b)** Positive selection signals in *Clip2* shown by multiple sequence alignment of protein sequences (continue).

**Supplementary Figure 21. Positive selection signals in gene *Clip2*.**

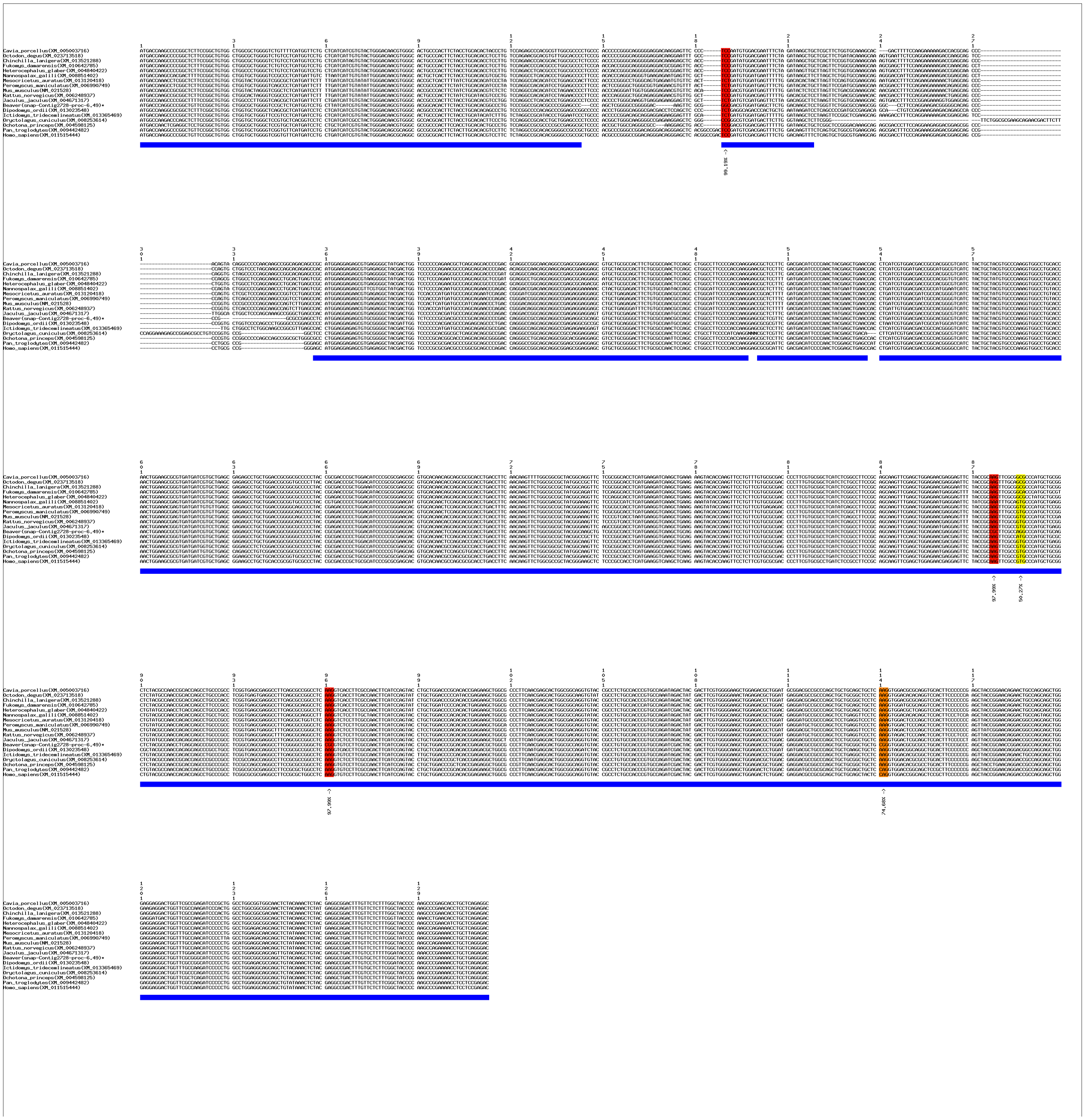

**(a)** Positive selection signals in in *Chst12* shown by multiple sequence alignment of coding sequences.

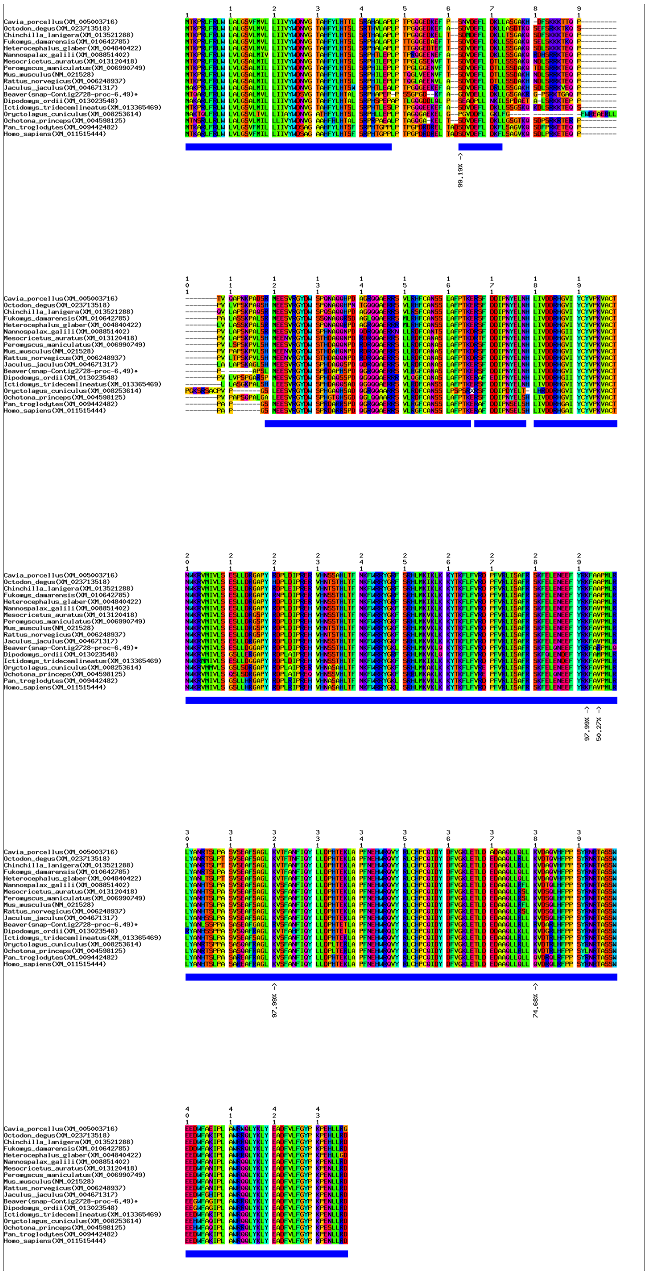

**(b)** Positive selection signals in *Chst12* shown by multiple sequence alignment of protein sequences.

**Supplementary Figure 22. Positive selection signals in gene *Chst12*.**

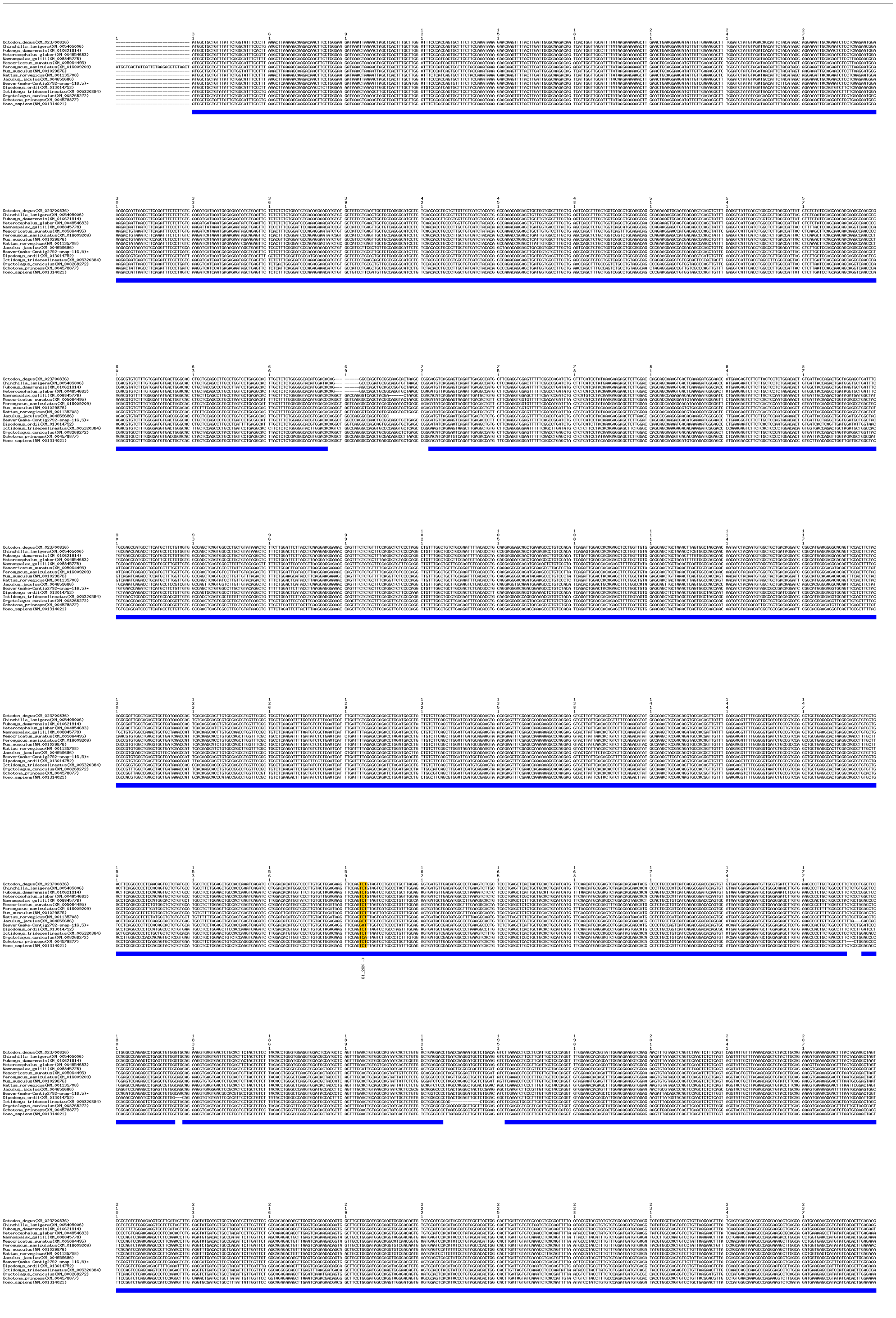

**(a)** Positive selection signals in *Urb2* shown by multiple sequence alignment of coding sequences.

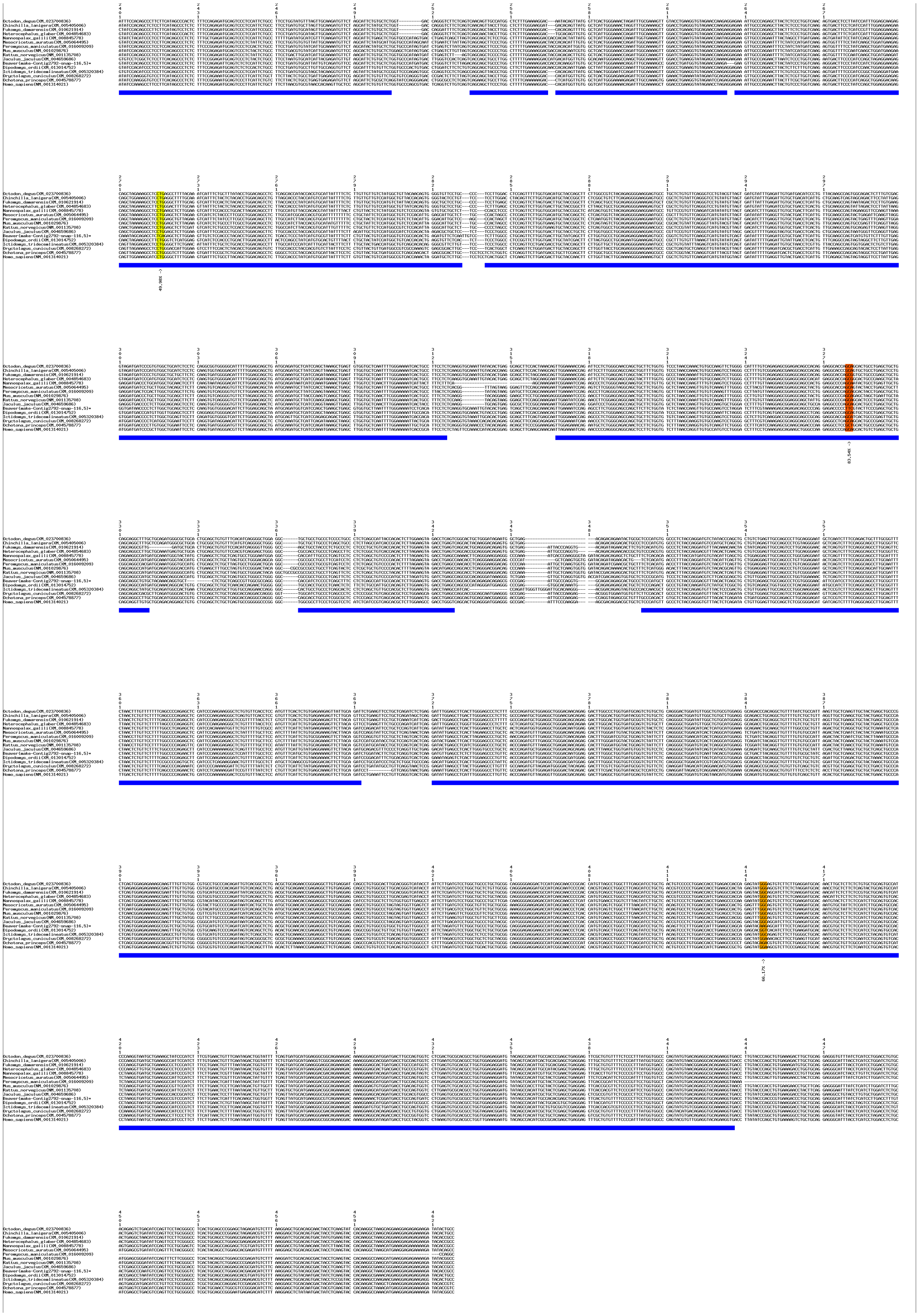

**(a)** Positive selection signals in *Urb2* shown by multiple sequence alignment of coding sequences (continue).

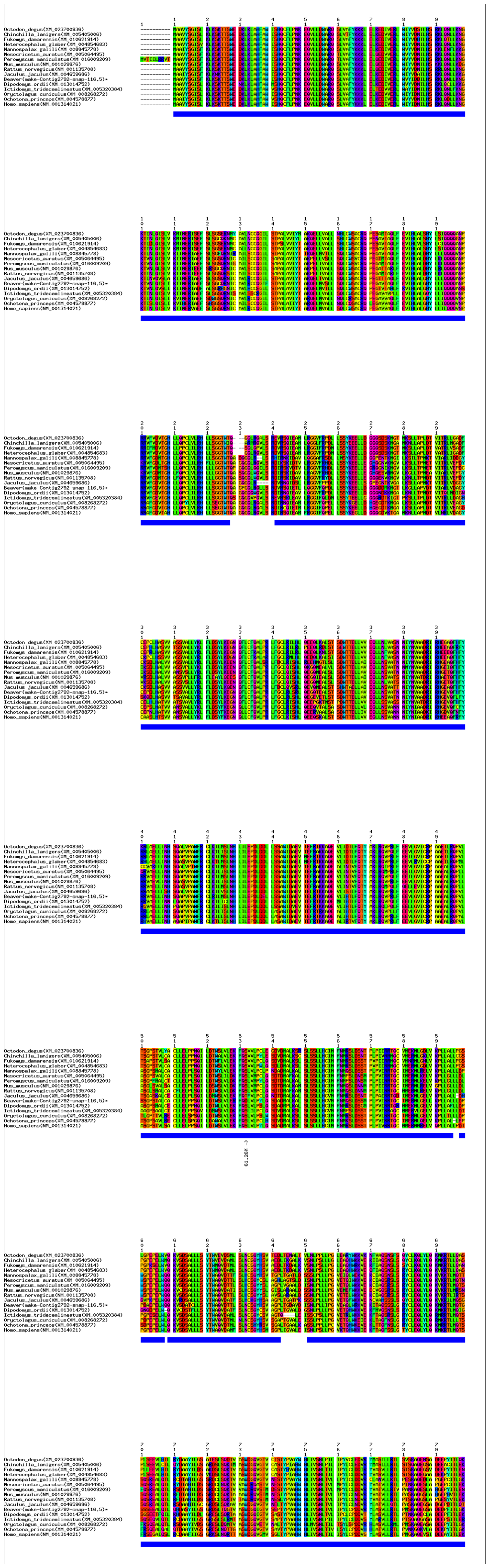

**(b)** Positive selection signals in *Urb2* shown by multiple sequence alignment of protein sequences.

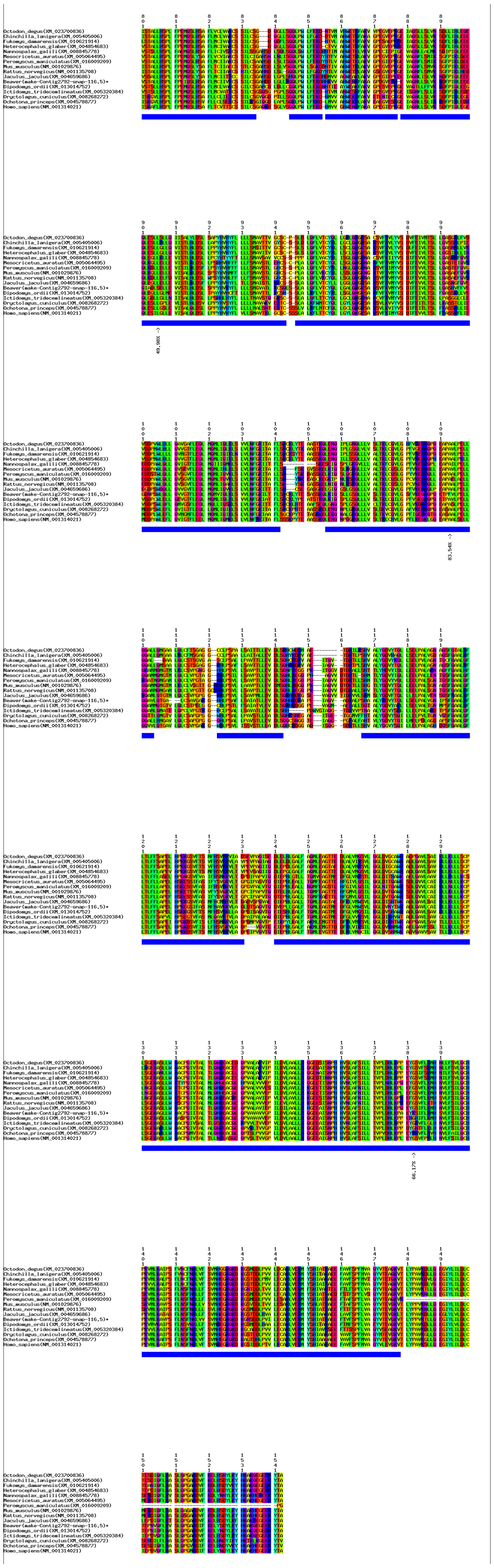

**(b)** Positive selection signals in *Urb2* shown by multiple sequence alignment of protein sequences.
**Supplementary Figure 23. Positive selection signals in gene *Urb2*.**

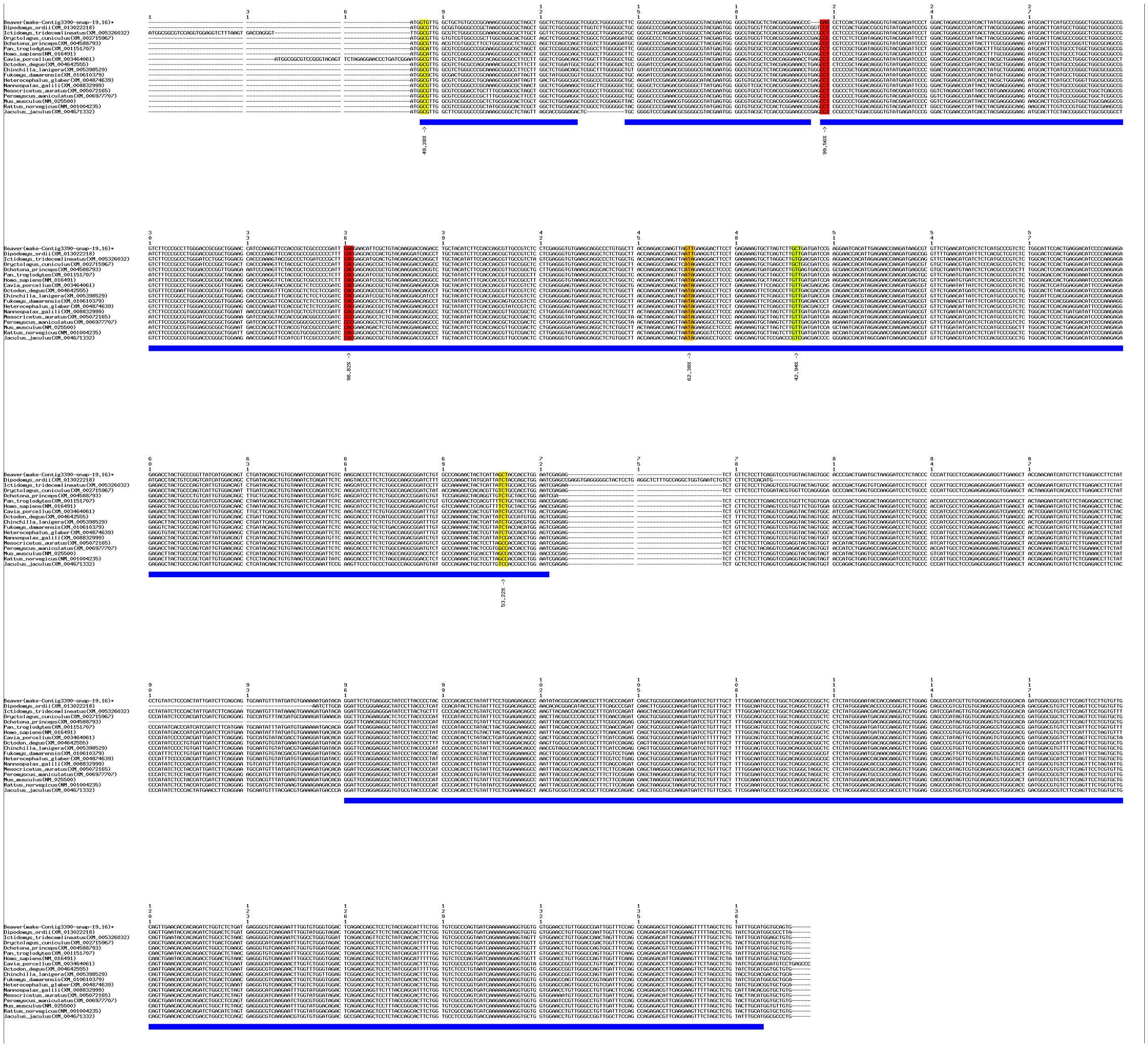

**(a)** Positive selection signals in *Mrpl37* shown by multiple sequence alignment of coding sequences.

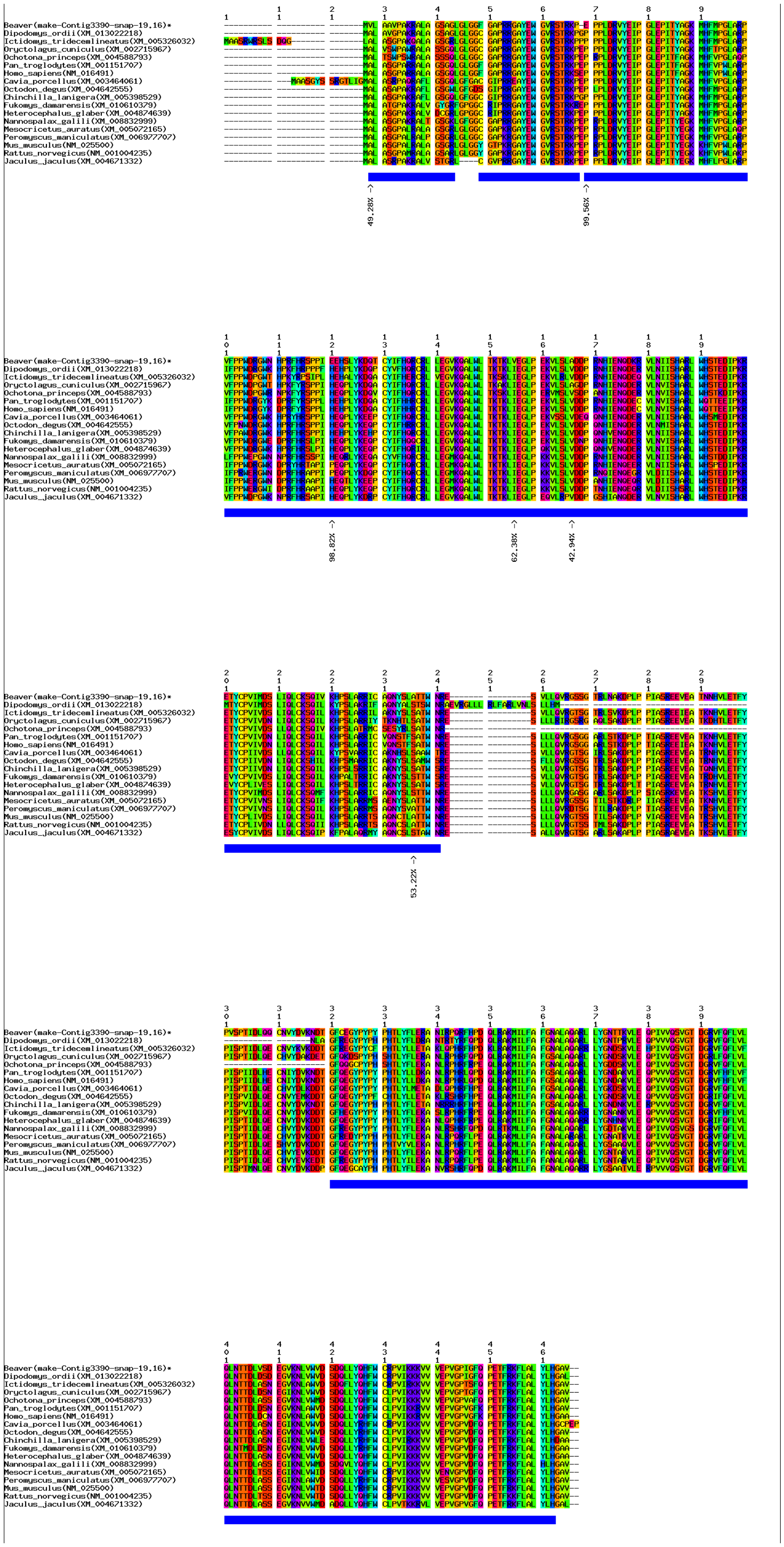

**(b)** Positive selection signals in *Mrpl37* shown by multiple sequence alignment of protein sequences.
**Supplementary Figure 24. Positive selection signals in gene *Mrpl37.***

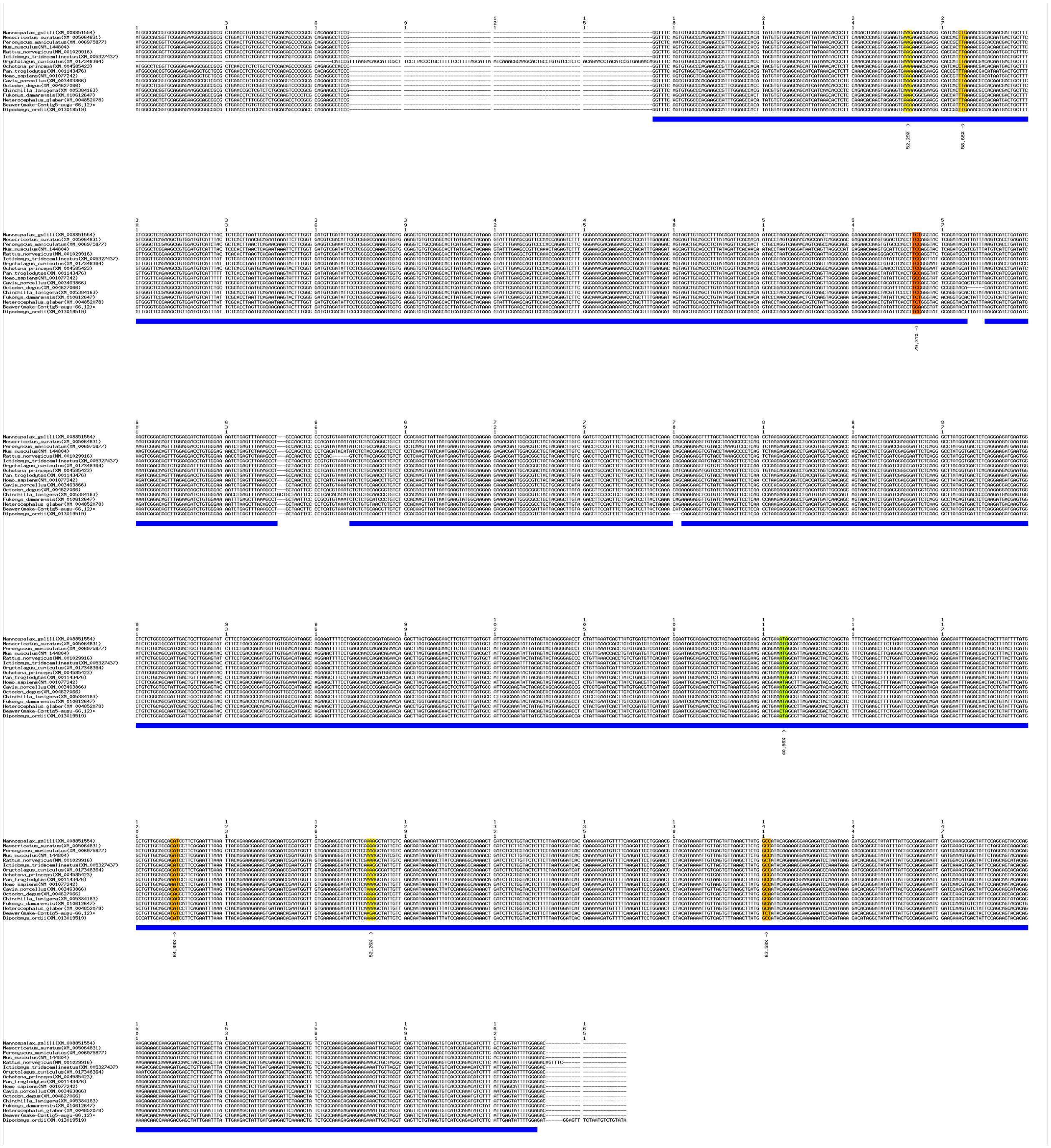

**(a)** Positive selection signals in *Depdc7* shown by multiple sequence alignment of coding sequences.

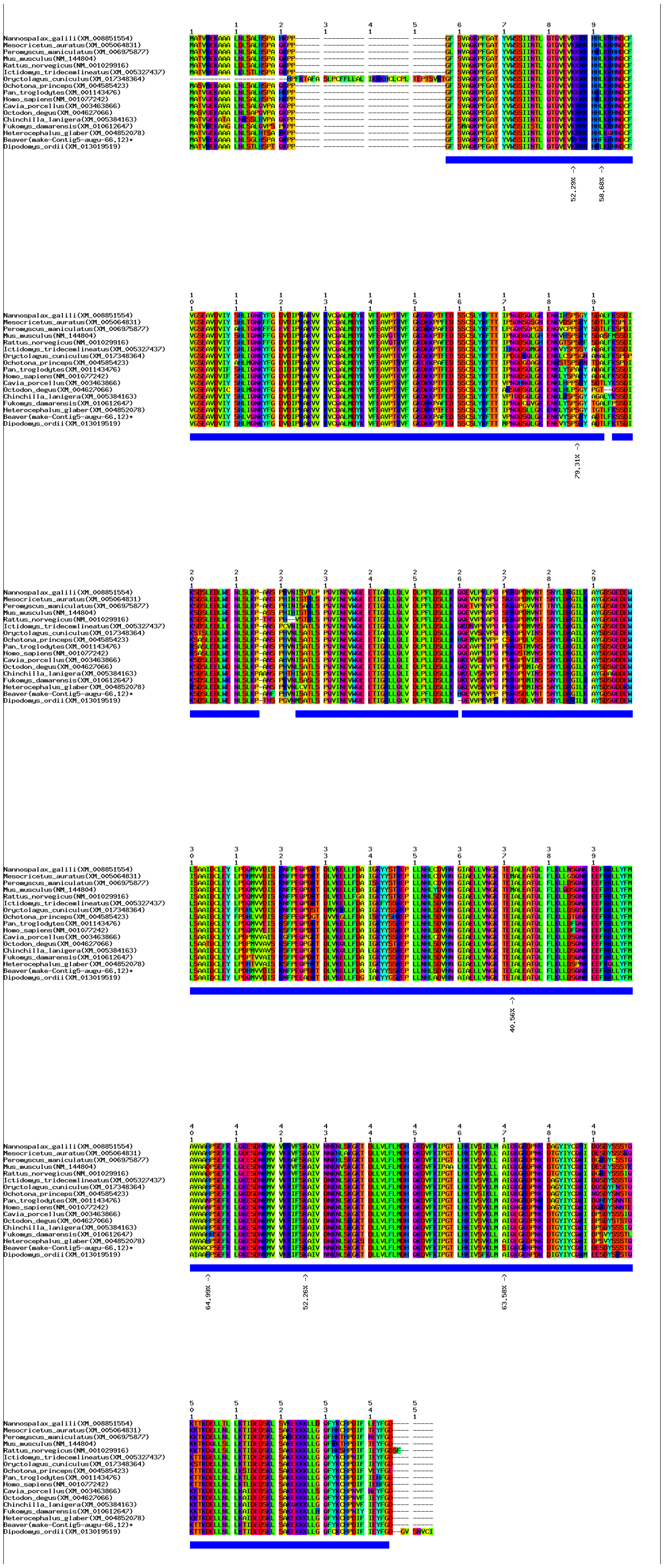

**(b)** Positive selection signals in *Depdc7* shown by multiple sequence alignment of protein sequences.
**Supplementary Figure 25. Positive selection signals in gene *Depdc7*.**

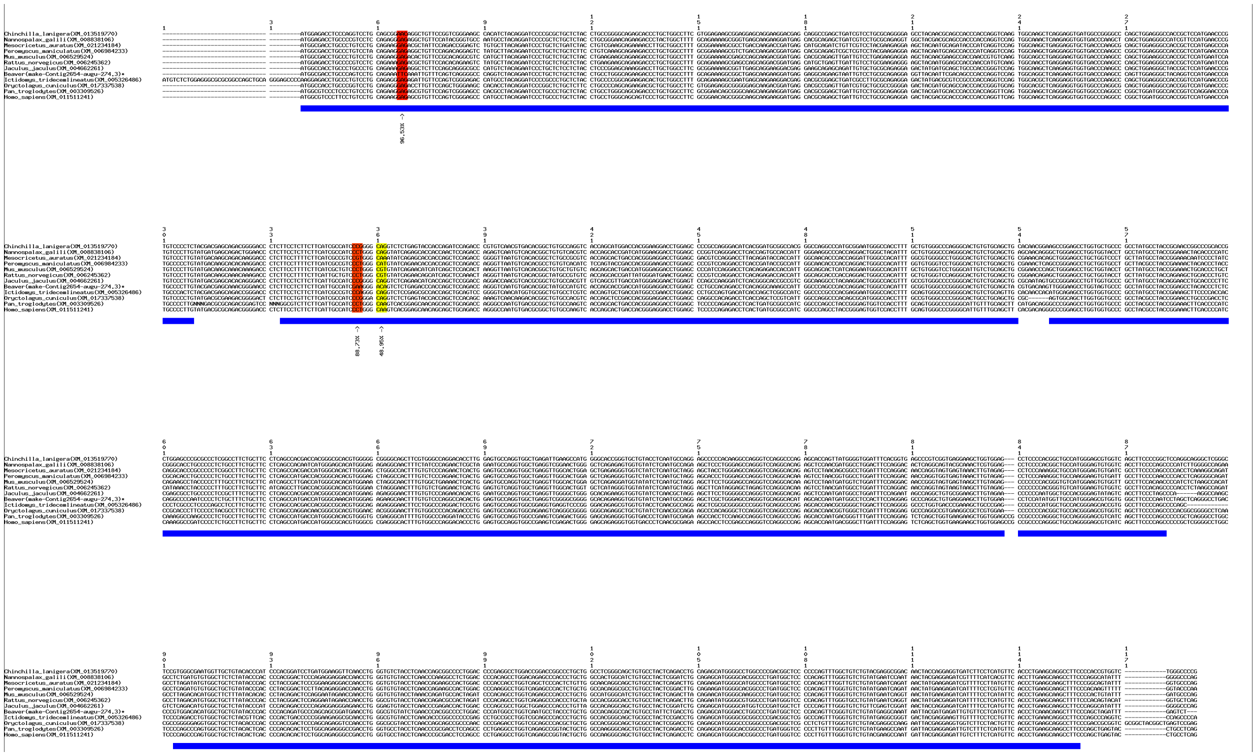

**(a)** Positive selection signals in *Neu2* shown by multiple sequence alignment of coding sequences.

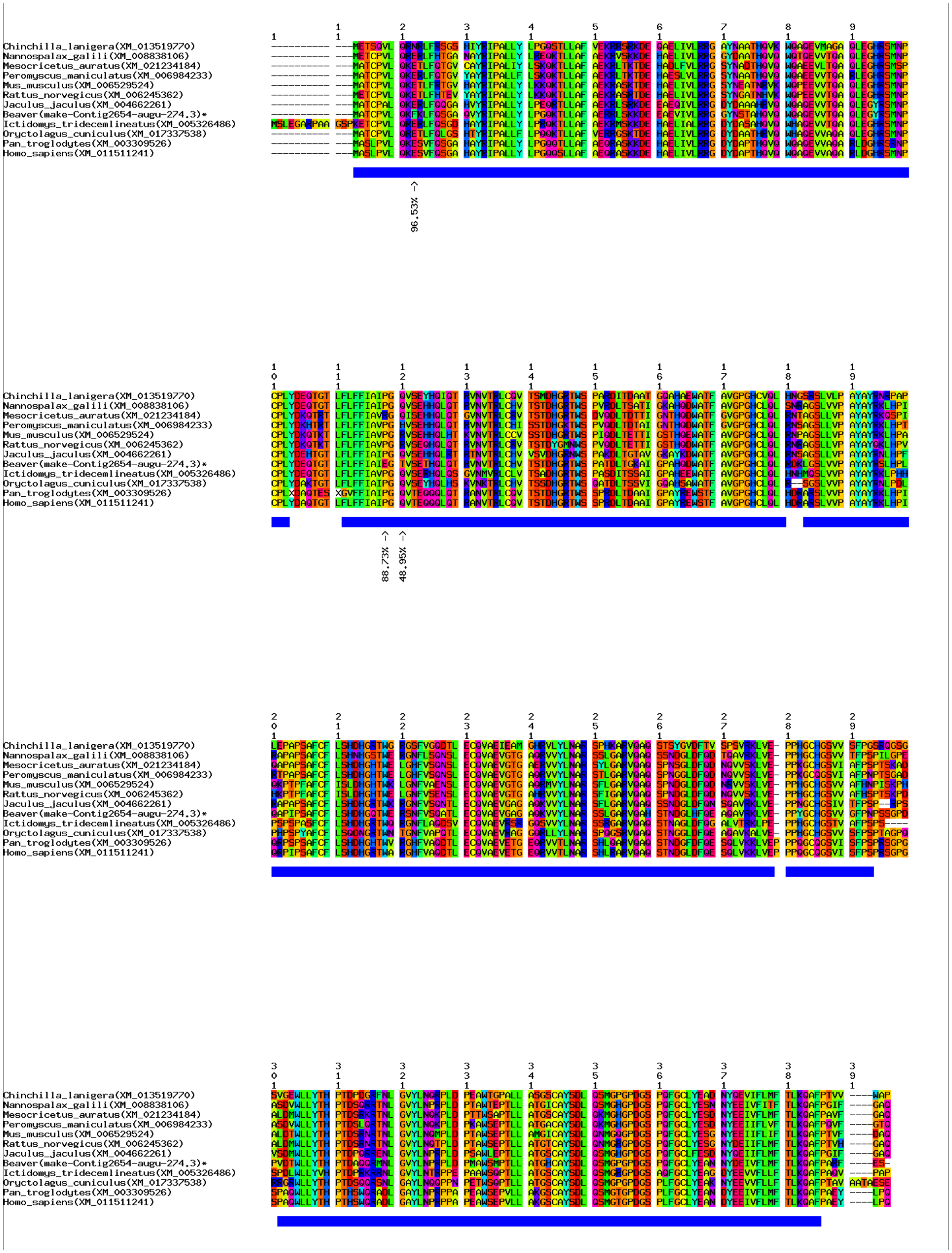

**(b)** Positive selection signals in *Neu2* shown by multiple sequence alignment of protein sequences.
**Supplementary Figure 26. Positive selection signals in gene *Neu2*.**

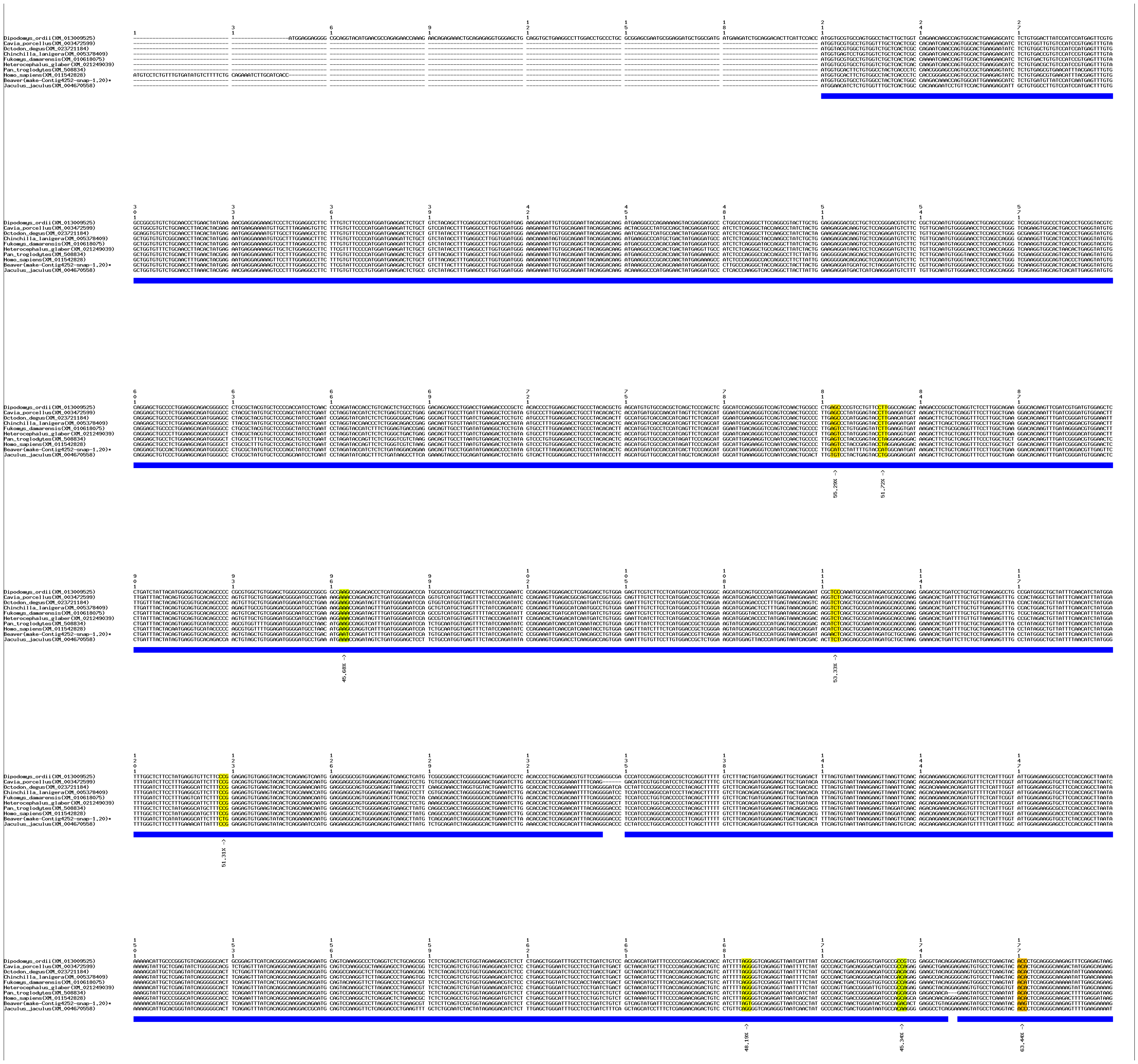

**(a)** Positive selection signals in *Vwa5a* shown by multiple sequence alignment of coding sequences.

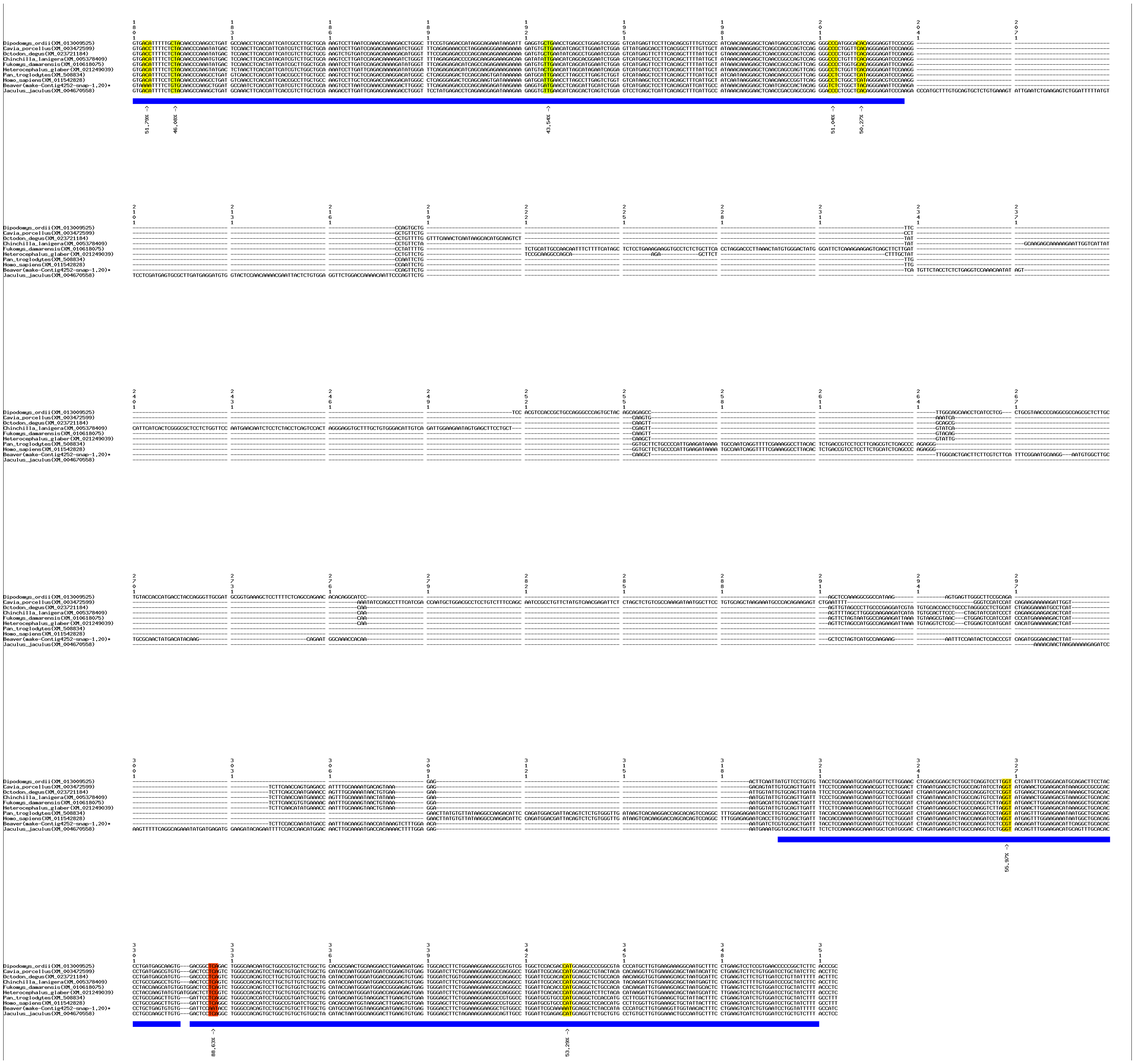

**(a)** Positive selection signals in *Vwa5a* shown by multiple sequence alignment of coding sequences (continue).

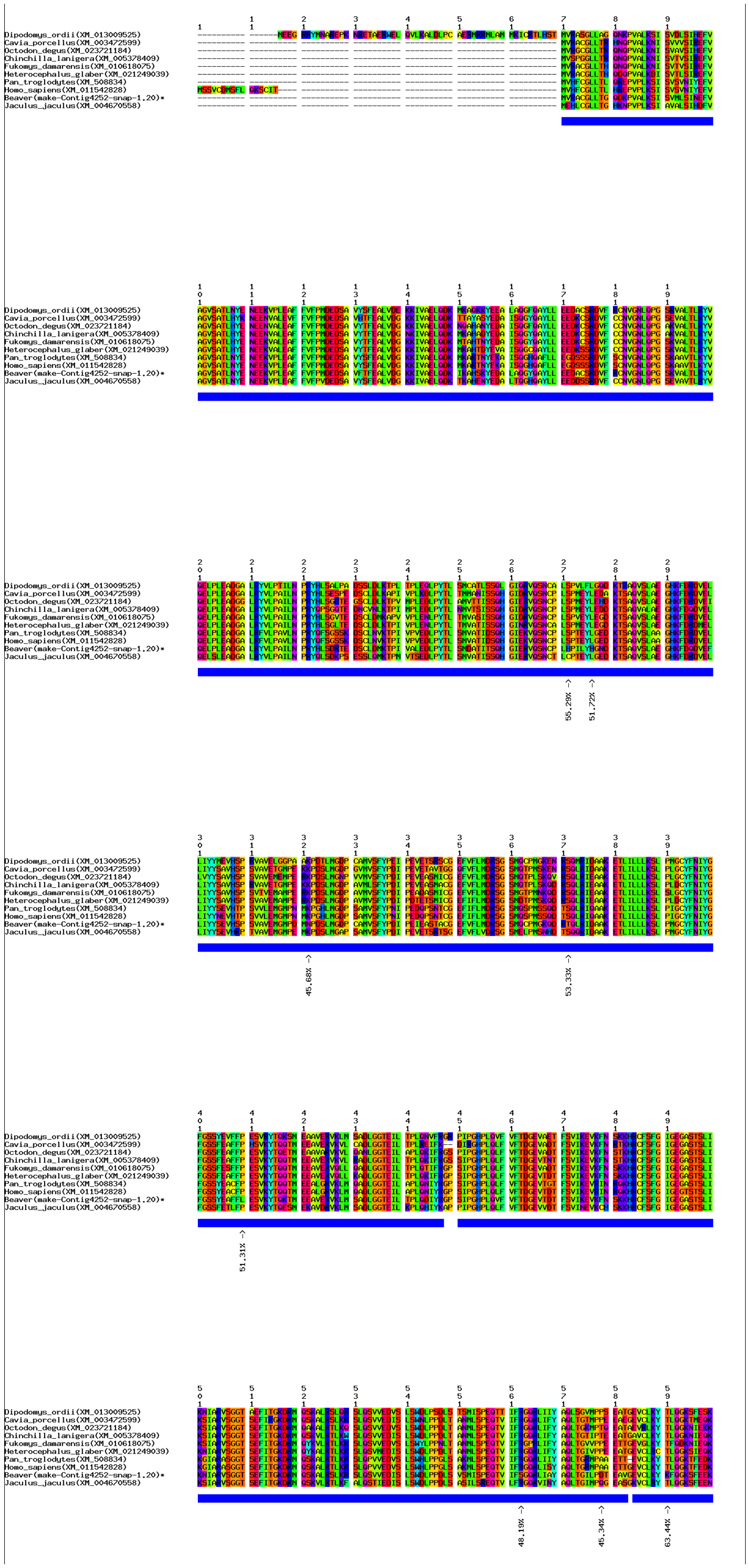

**(b)** Positive selection signals in *Vwa5a* shown by multiple sequence alignment of protein sequences.

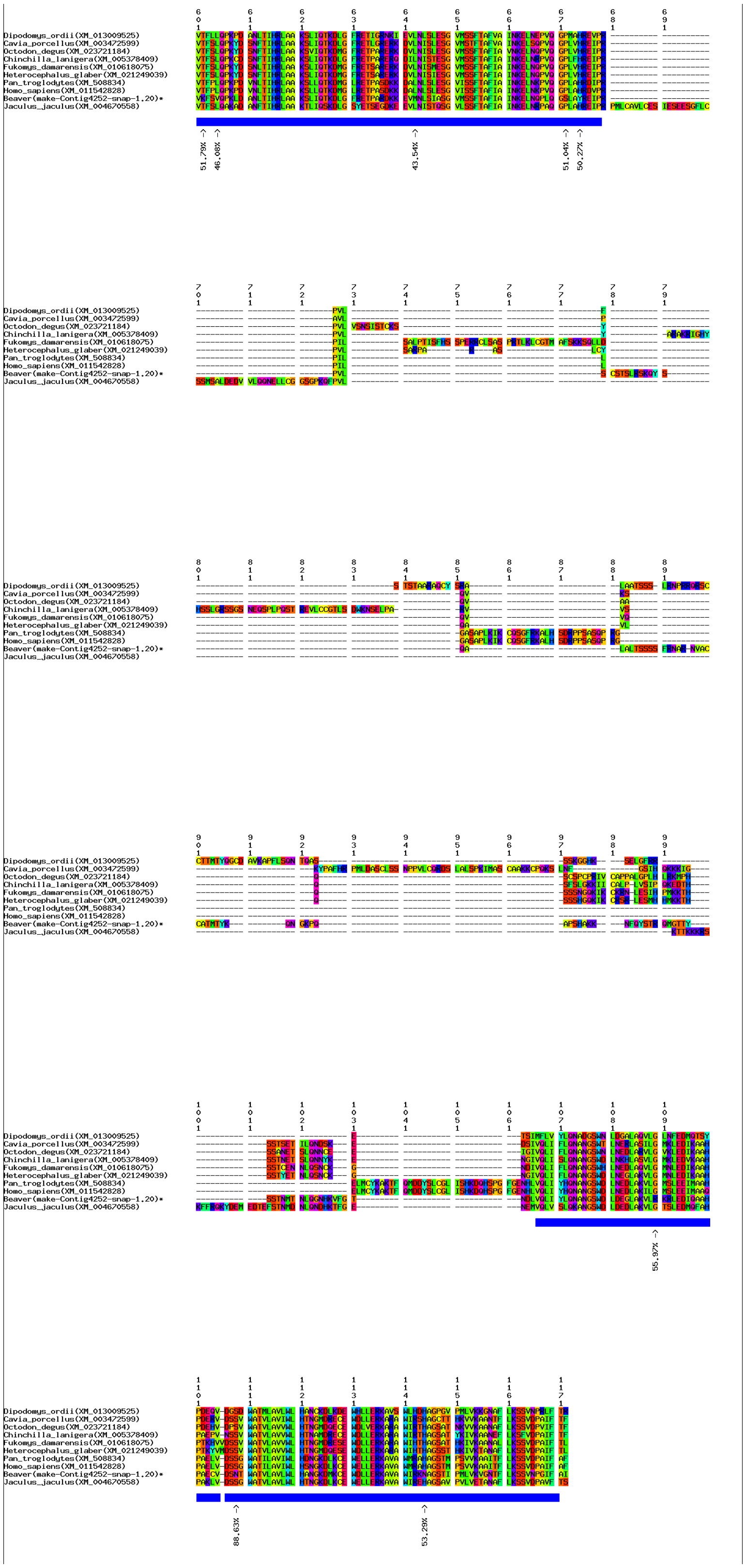

**(b)** Positive selection signals in *Vwa5a* shown by multiple sequence alignment of protein sequences (continue).

**Supplementary Figure 27. Positive selection signals in gene *Vwa5a*.**

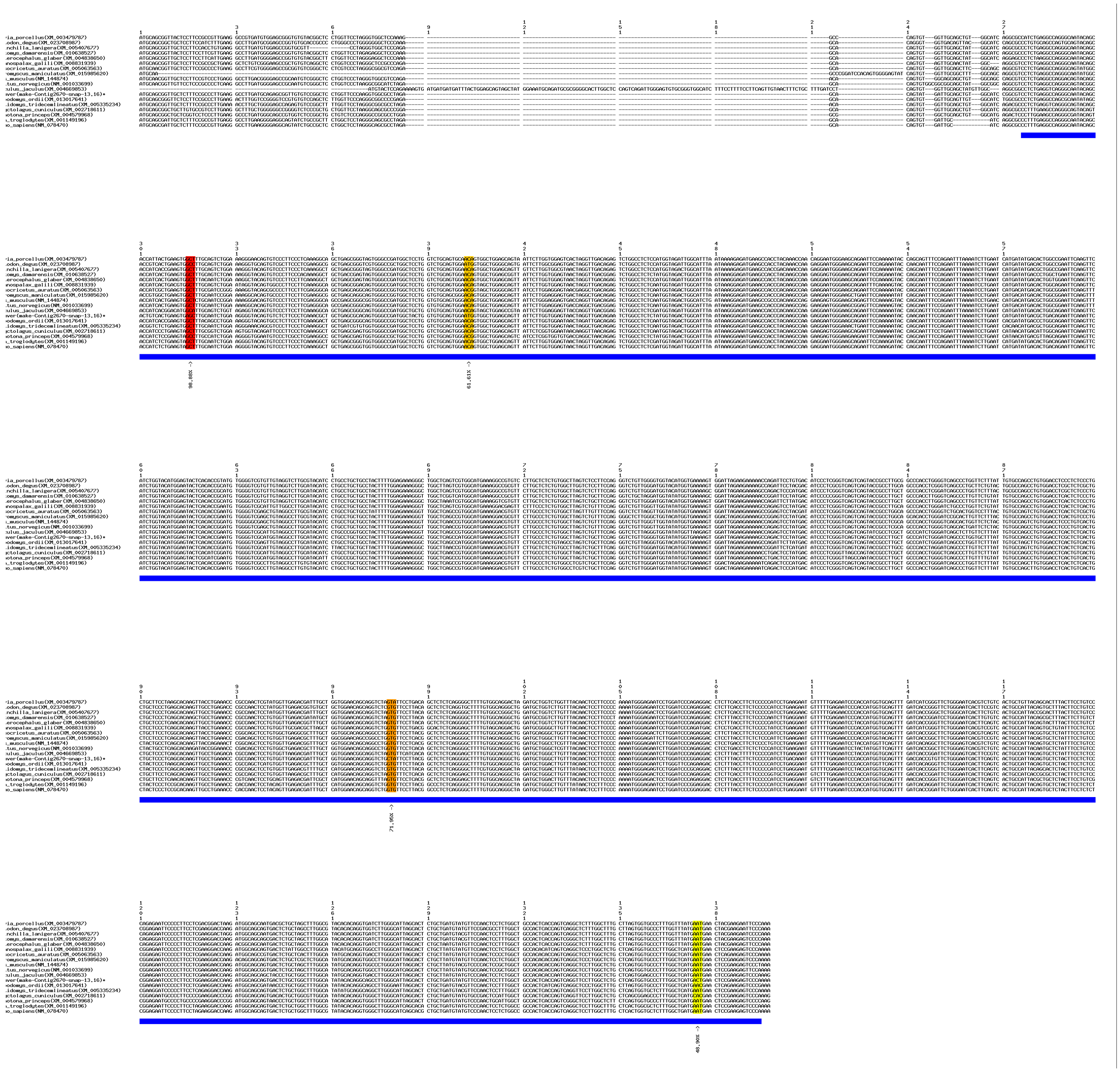

**(a)** Positive selection signals in *Cox15* shown by multiple sequence alignment of coding sequences.

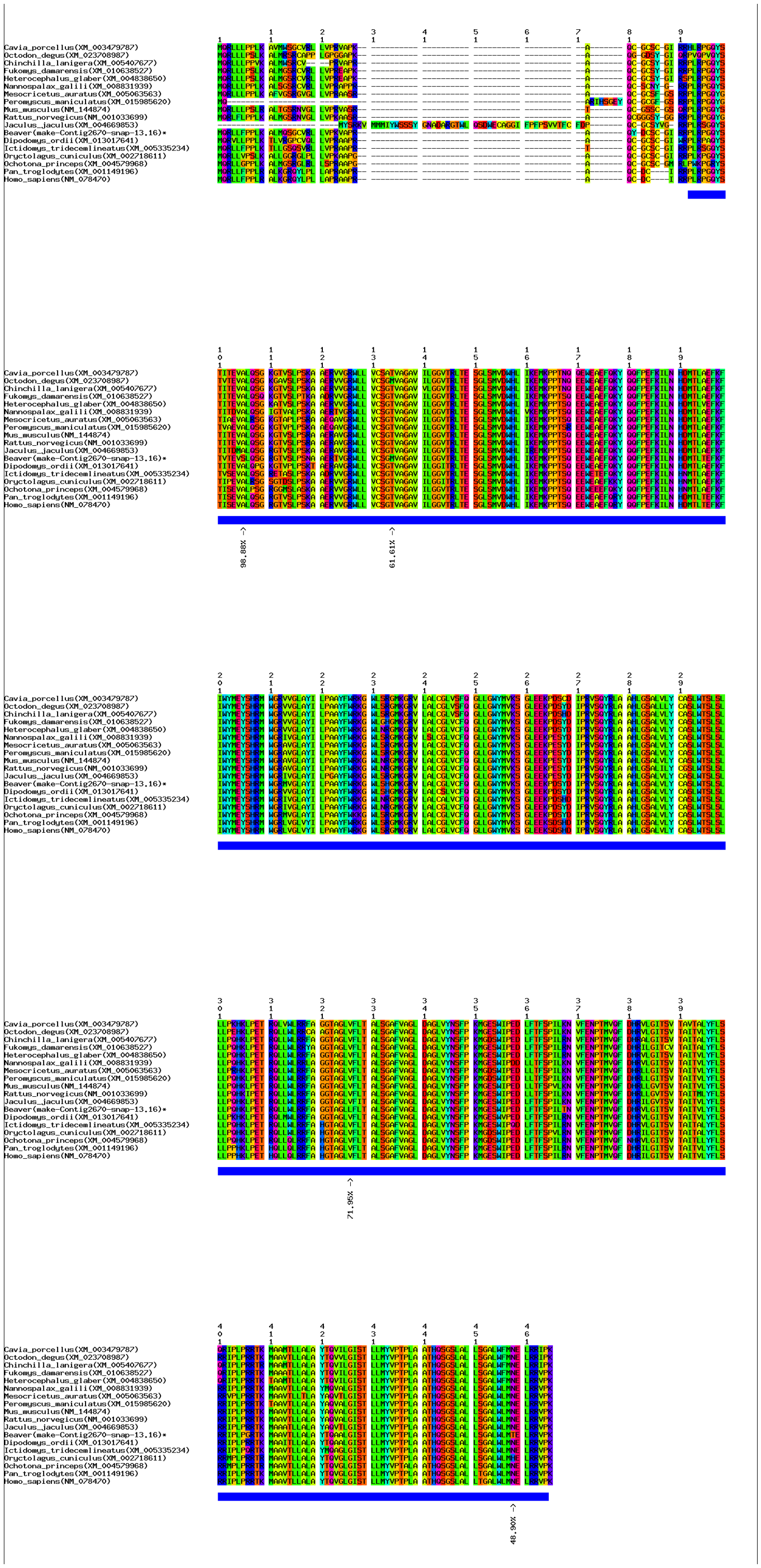

**(b)** Positive selection signals in *Cox15* shown by multiple sequence alignment of protein sequences (continue).

**Supplementary Figure 28. Positive selection signals in gene *Cox15*.**

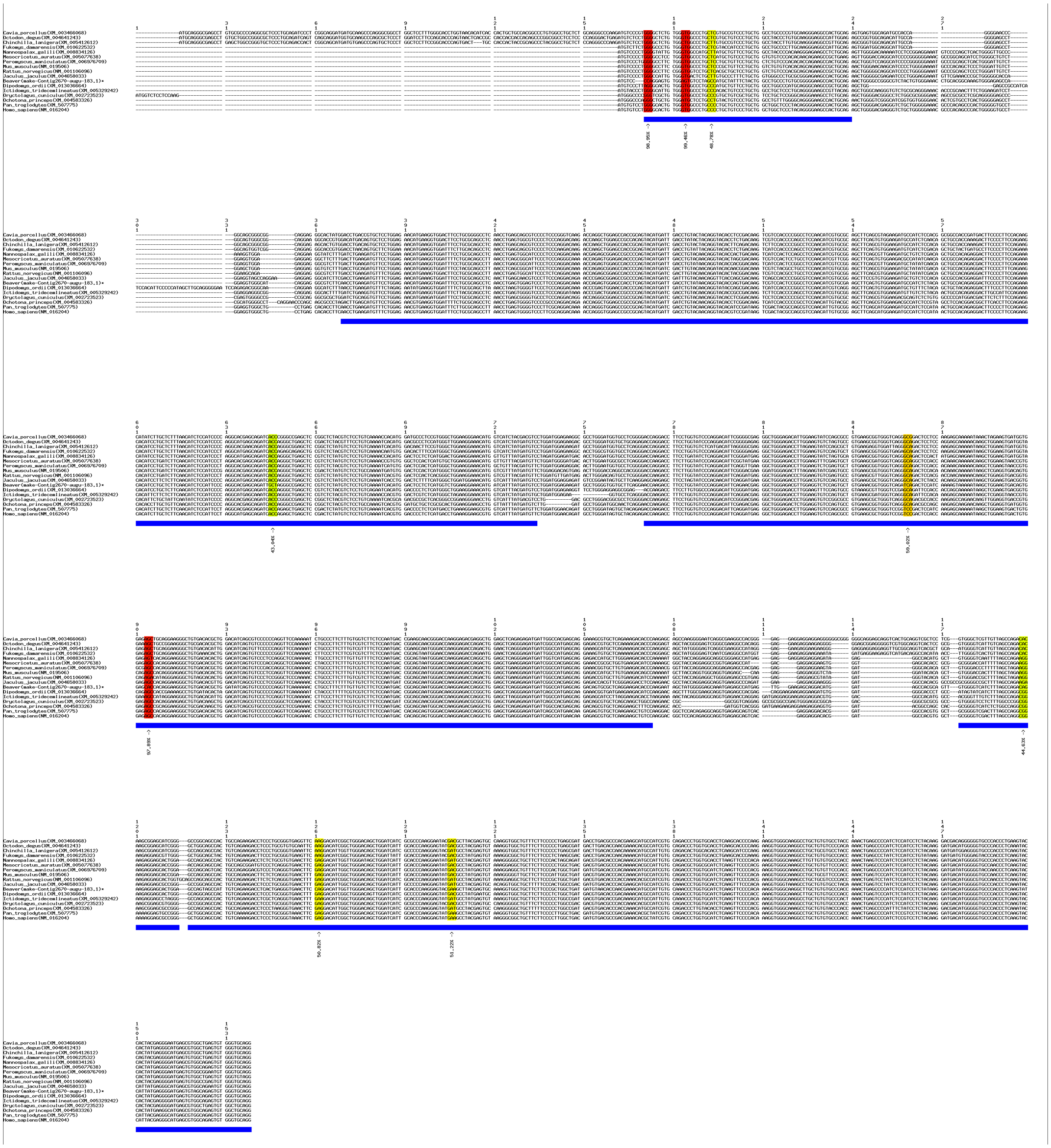

**(a)** Positive selection signals in *Gdf2* shown by multiple sequence alignment of coding sequences.

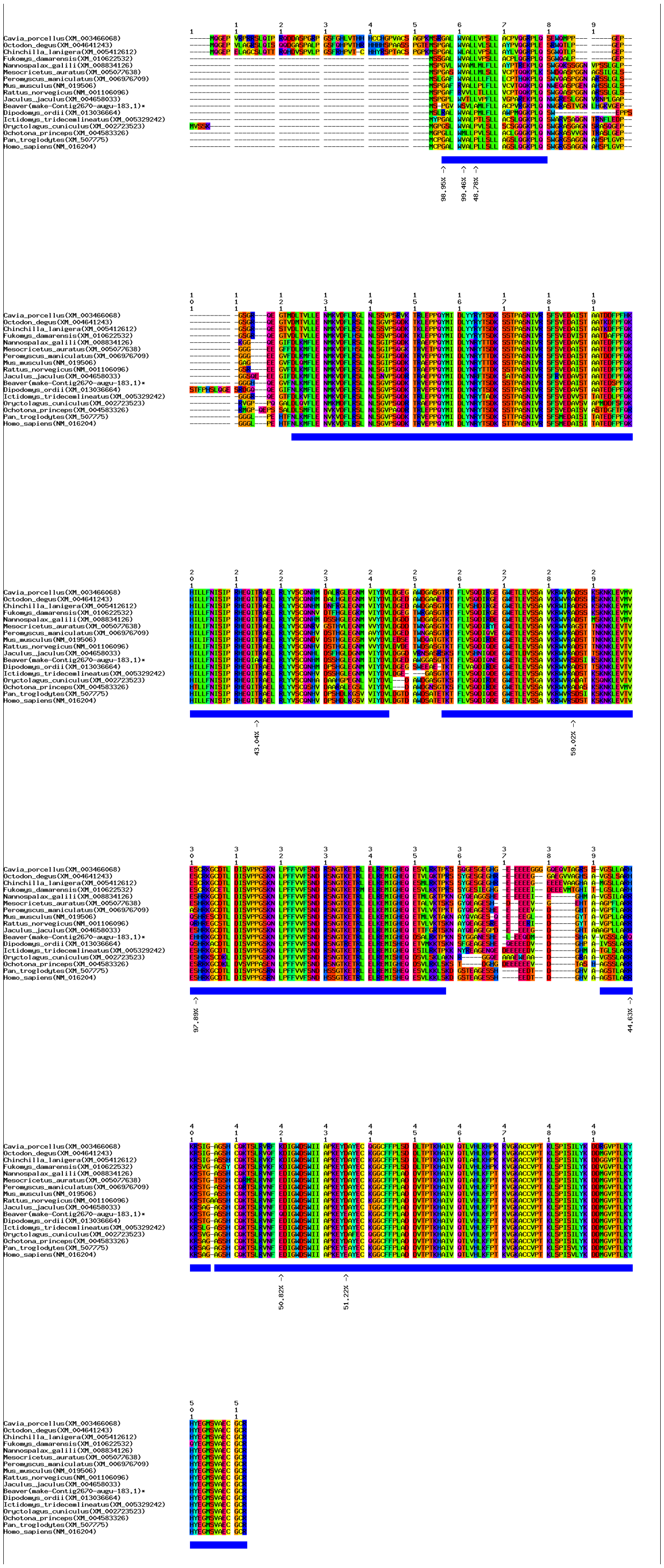

**(b)** Positive selection signals in *Gdf2* shown by multiple sequence alignment of protein sequences.
**Supplementary Figure 29. Positive selection signals in gene *Gdf2*.**

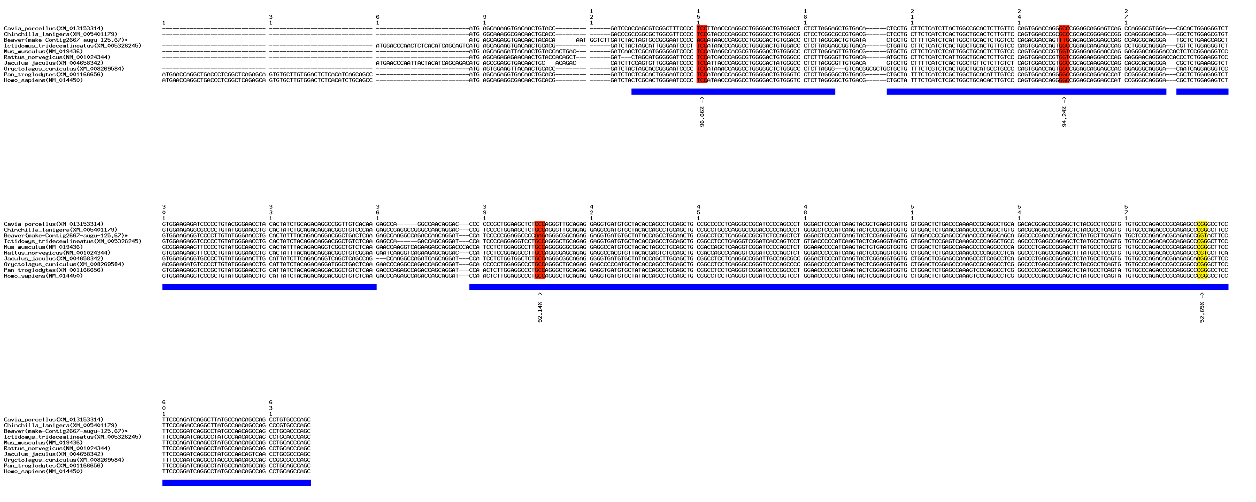

**(a)** Positive selection signals in *Sit1* shown by multiple sequence alignment of coding sequences.

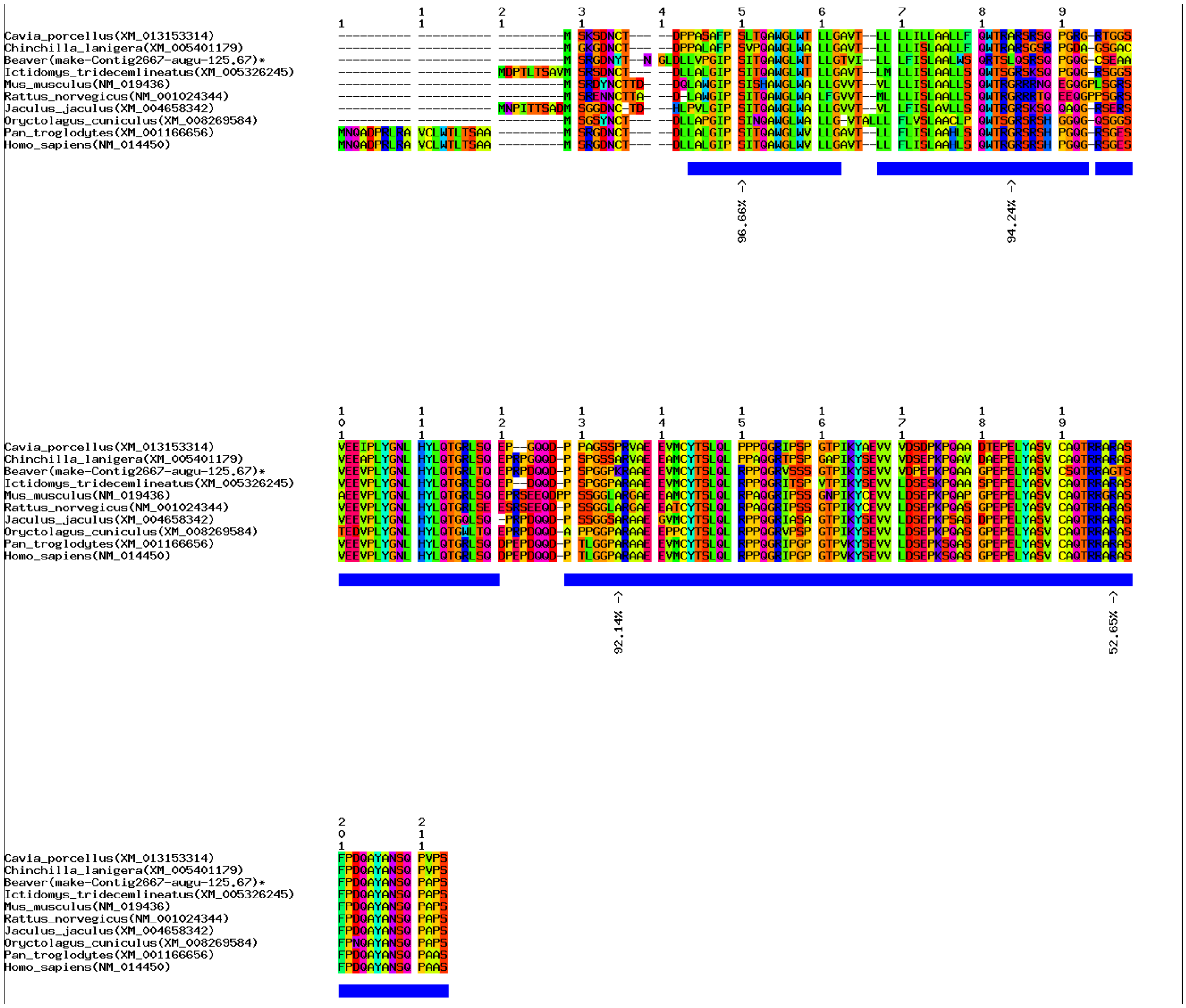

**(b)** Positive selection signals in *Sit1* shown by multiple sequence alignment of protein sequences.
**Supplementary Figure 30. Positive selection signals in gene *Sit1*.**

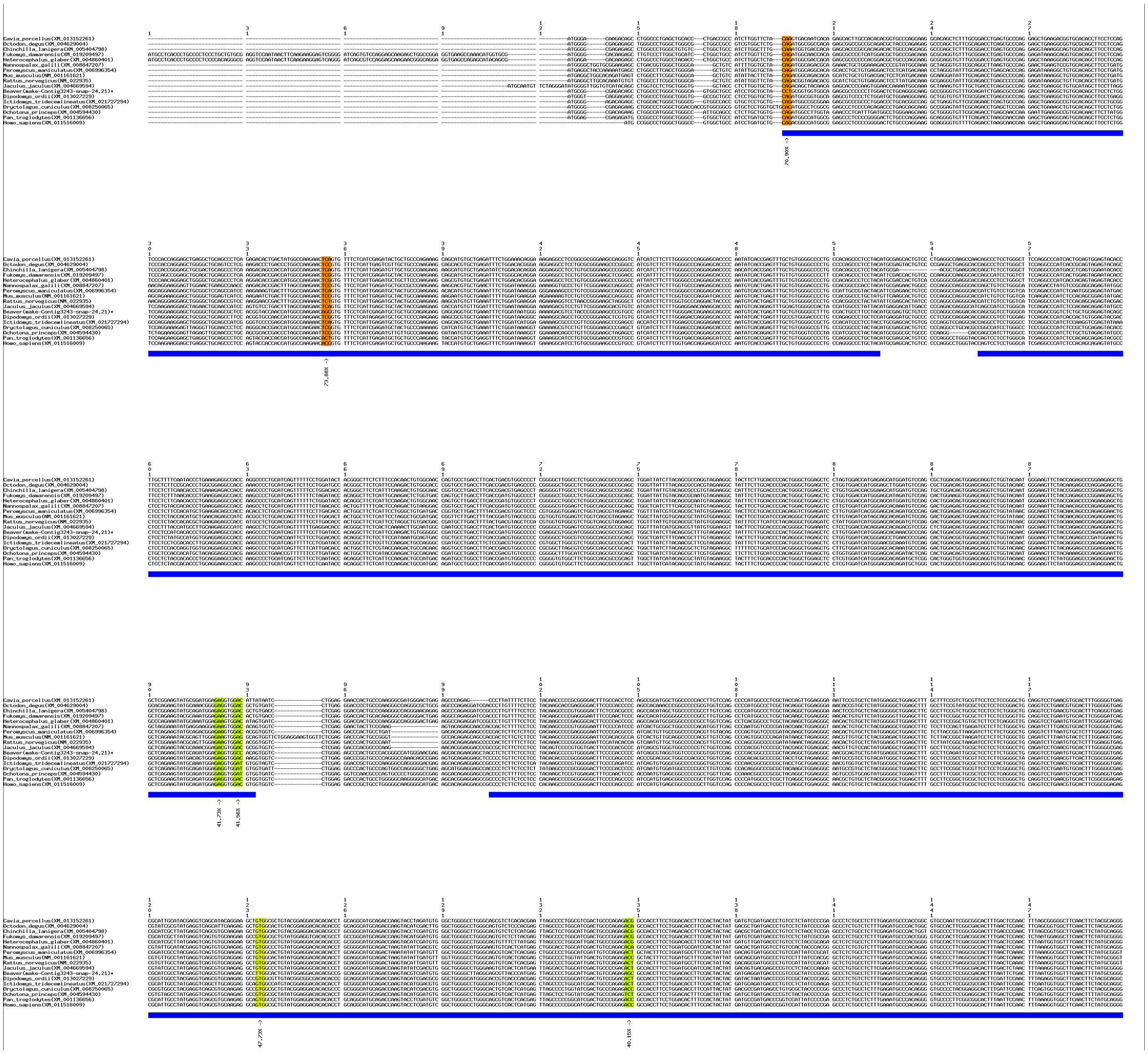

**(a)** Positive selection signals in *Aoc1* shown by multiple sequence alignment of coding sequences.

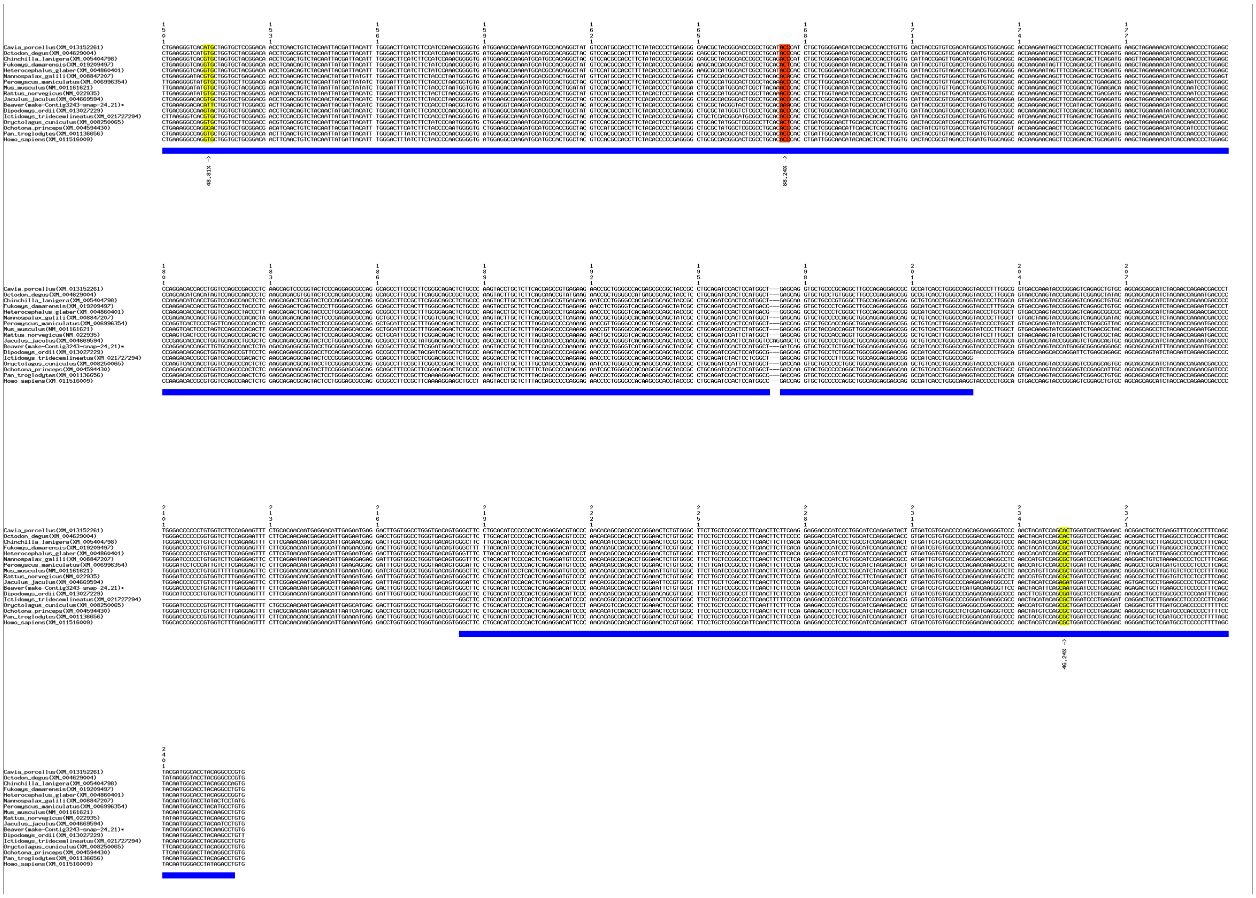

**(a)** Positive selection signals in *Aoc1* shown by multiple sequence alignment of coding sequences (continue).

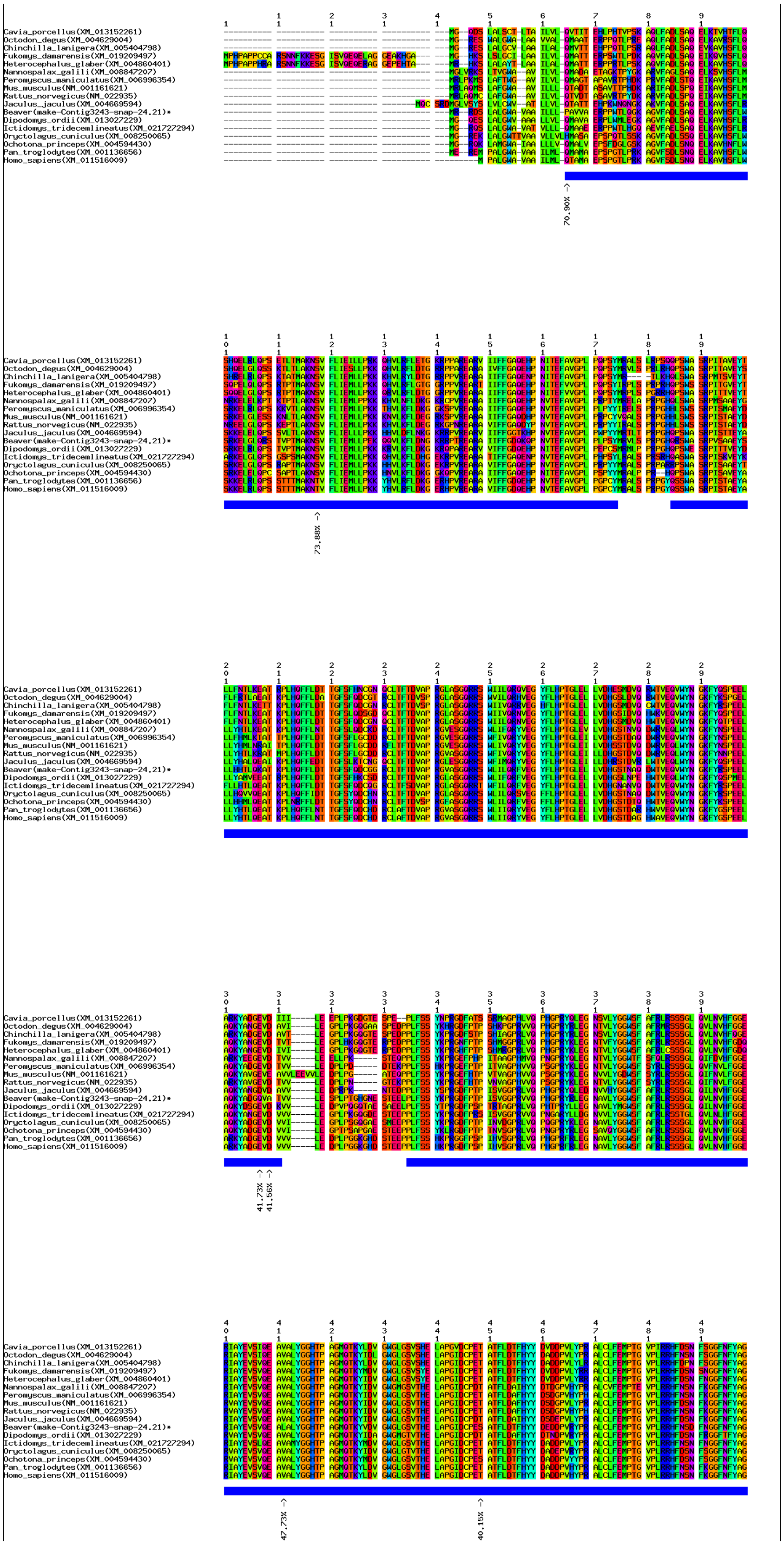


**(b)** Positive selection signals in *Aoc1* shown by multiple sequence alignment of protein sequences.

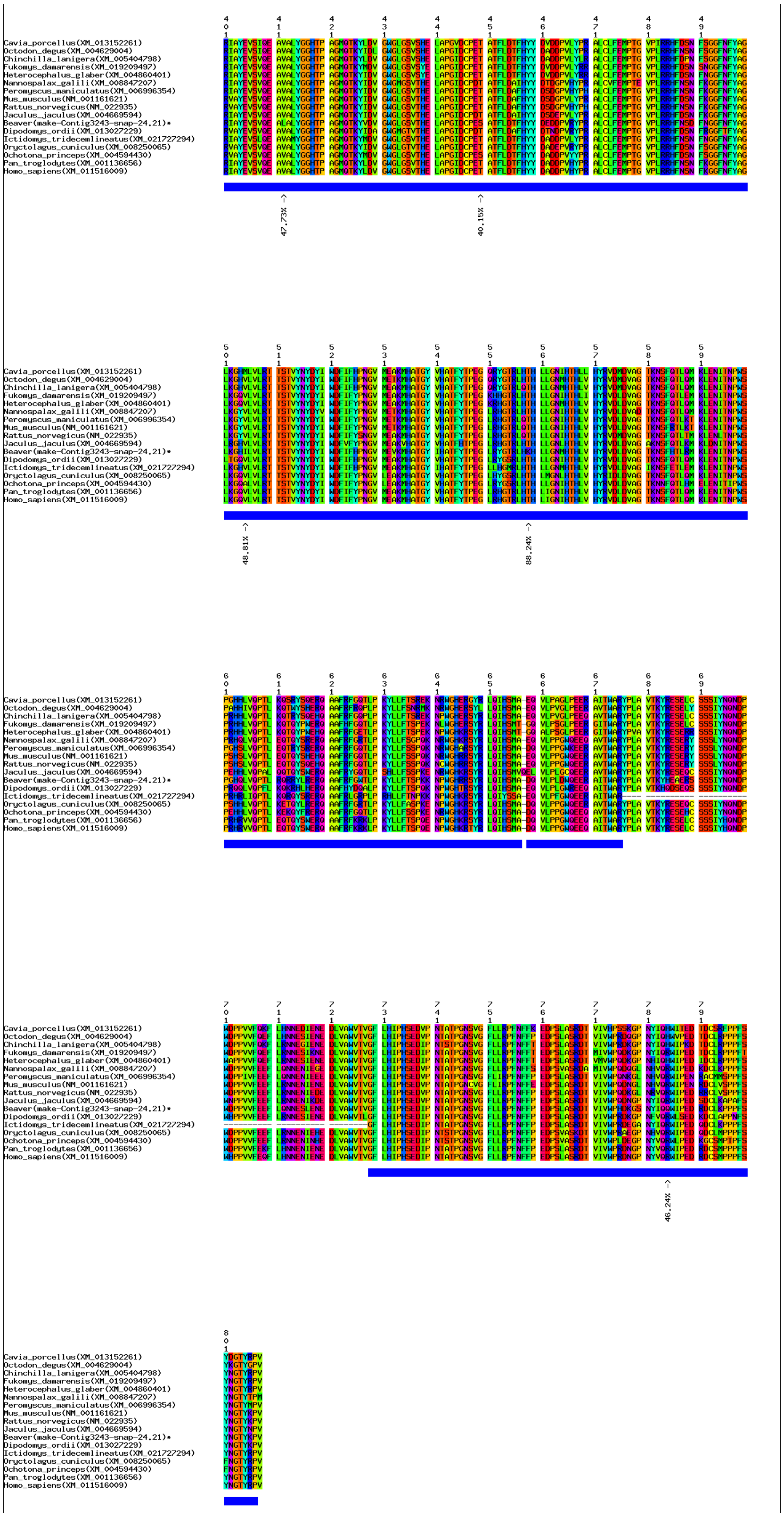

**(b)** Positive selection signals in *Aoc1* shown by multiple sequence alignment of protein sequences (continue).

**Supplementary Figure 31. Positive selection signals in gene *Aoc1*.**

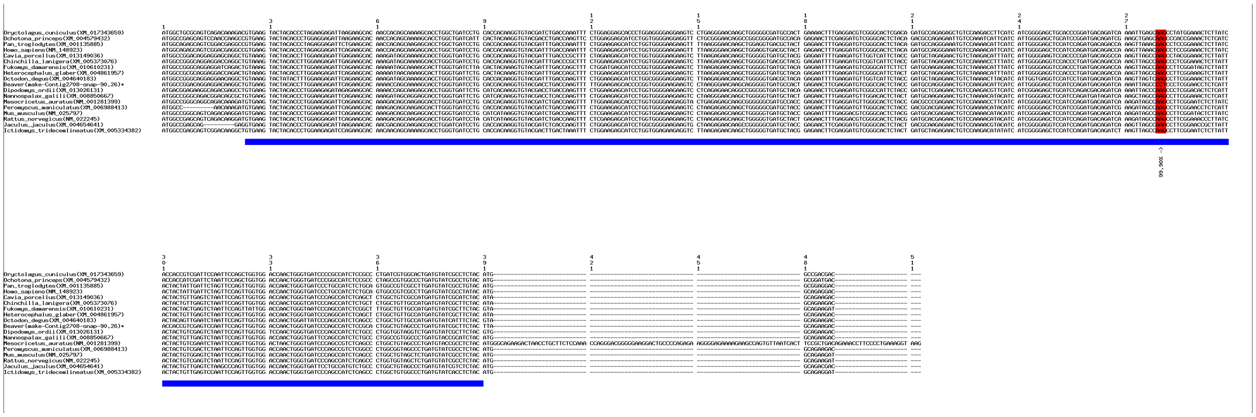

**(a)** Positive selection signals in *Cyb5a* shown by multiple sequence alignment of coding sequences.

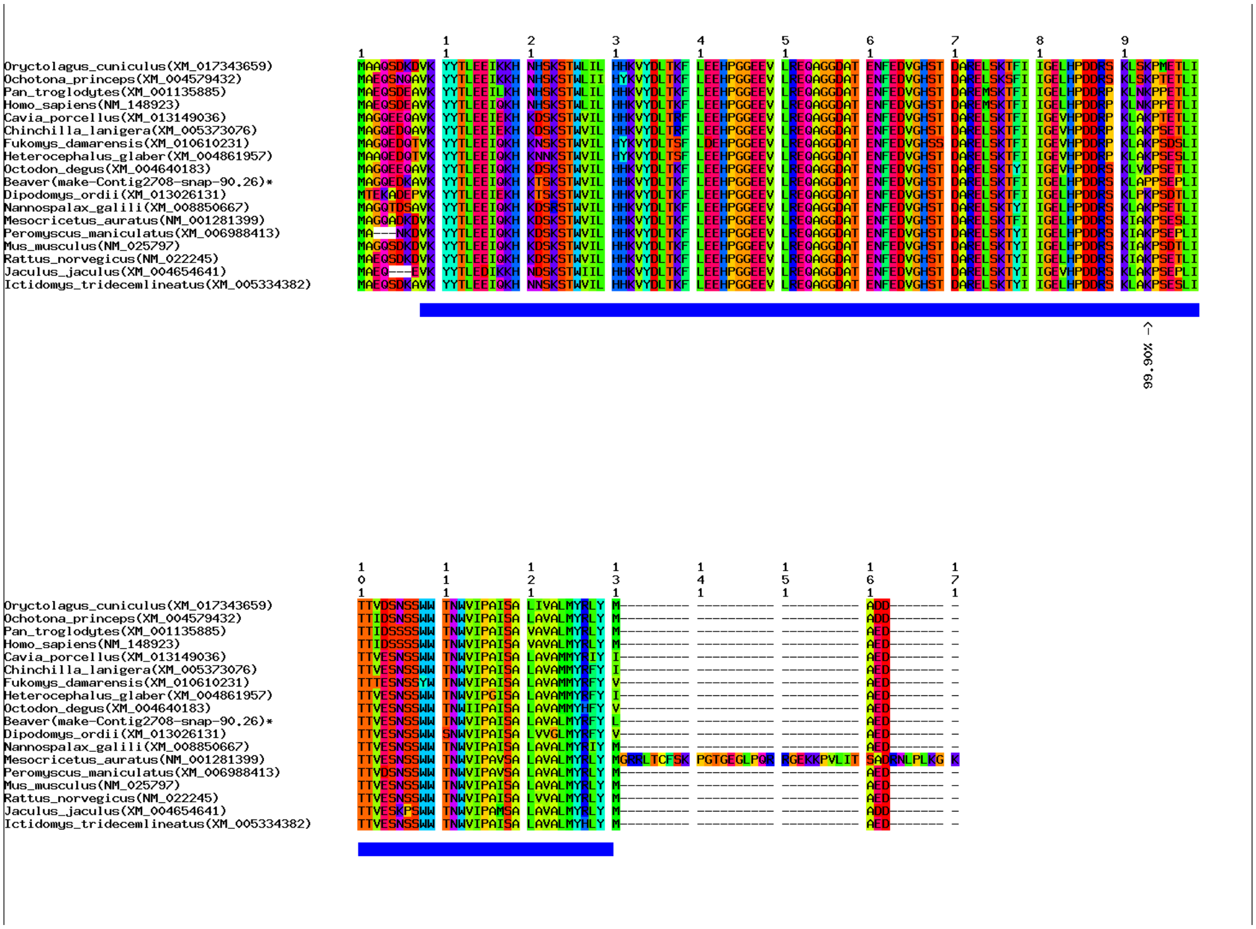


**(b)** Positive selection signals in *Cyb5a* shown by multiple sequence alignment of protein sequences.

**Supplementary Figure 32. Positive selection signals in gene *Cyb5a*.**

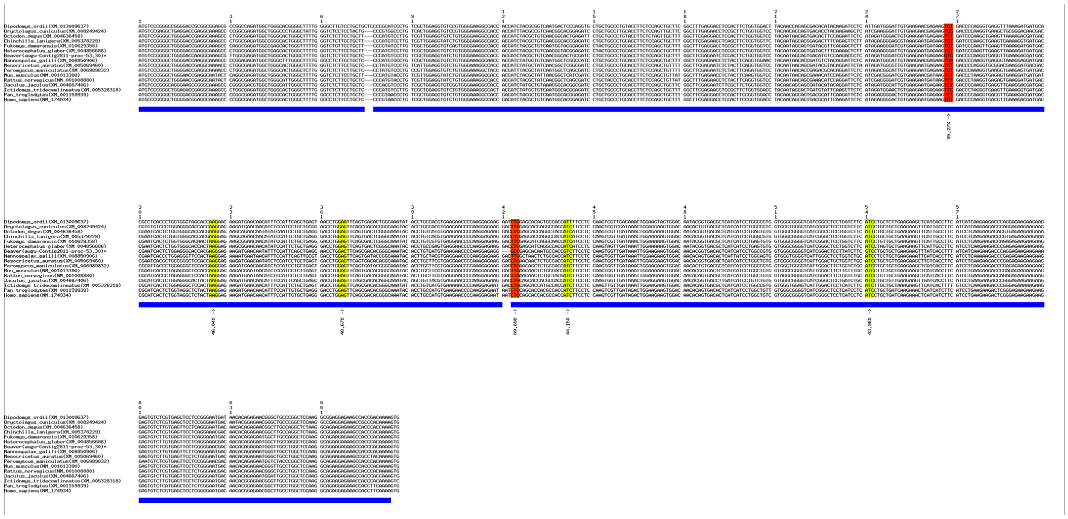

**(a)** Positive selection signals in *Scn4b* shown by multiple sequence alignment of coding sequences.

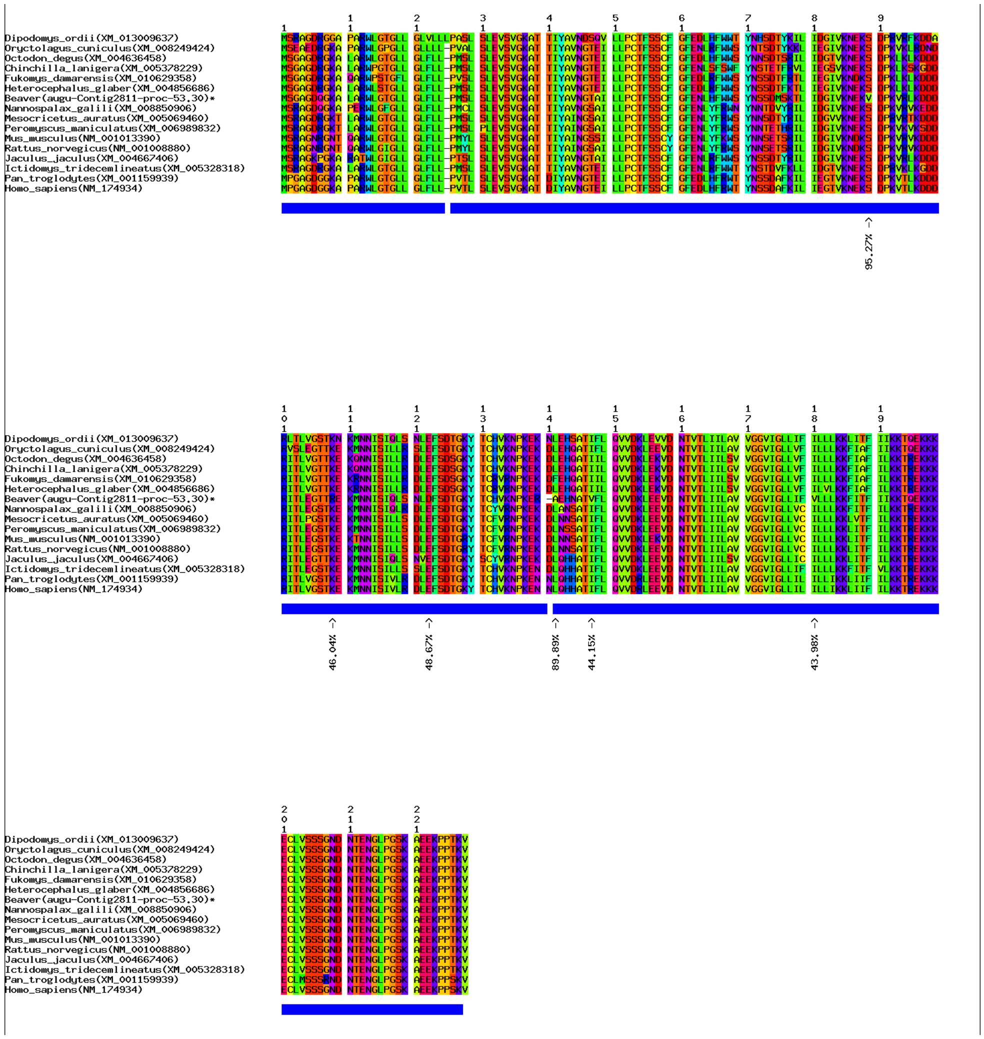

**(b)** Positive selection signals in *Scn4b* of gene shown by multiple sequence alignment of protein sequences.

**Supplementary Figure 33. Positive selection signals in gene *Scn4b*.**

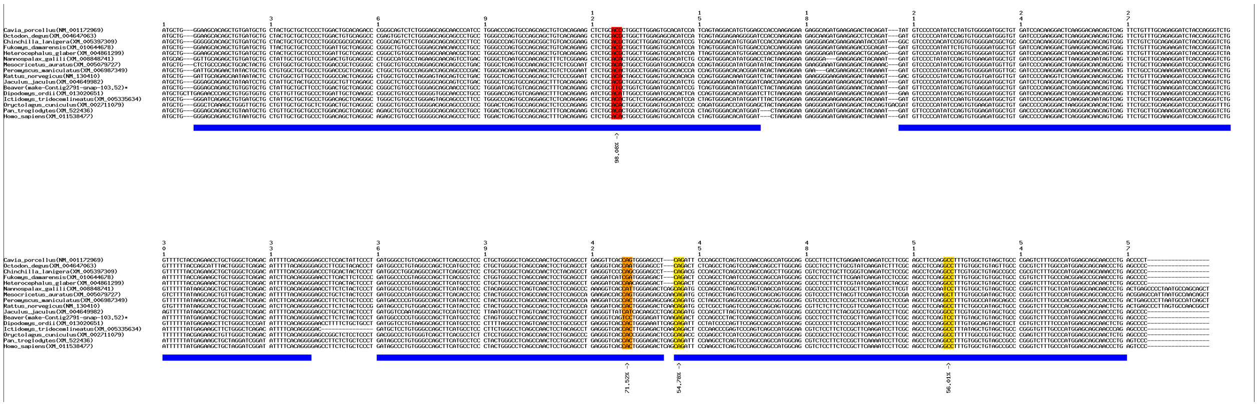

**(a)** Positive selection signals in *Il23a* shown by multiple sequence alignment of coding sequences.

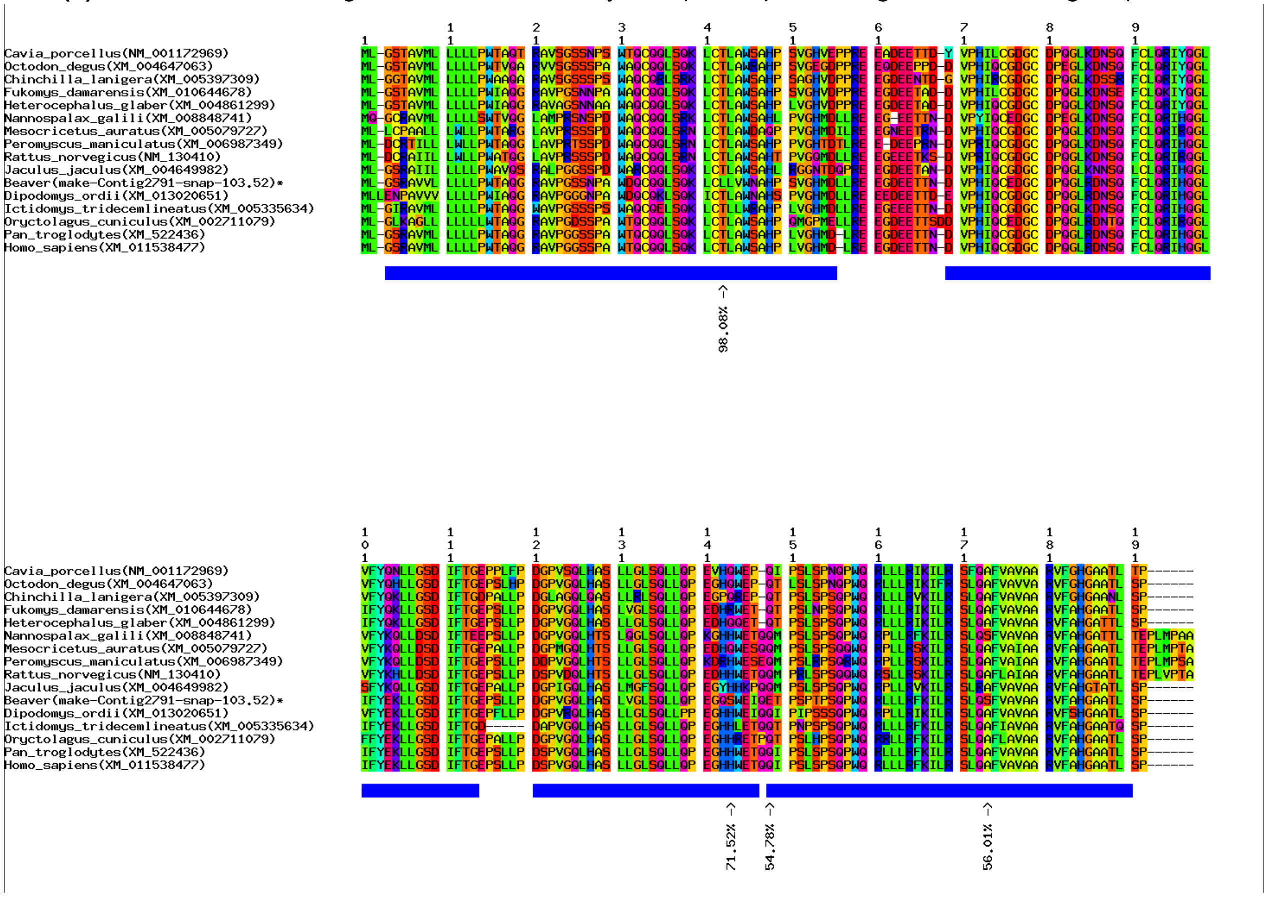


**(b)** Positive selection signals in *Il23a* shown by multiple sequence alignment of protein sequences.

**Supplementary Figure 34. Positive selection signals in gene *Il23a*.**
